# Missense mutations: Backbone structure positional effects

**DOI:** 10.1101/2024.12.23.630208

**Authors:** Ivan Perez, Ulrike Stege, Hosna Jabbari

**Affiliations:** Department of Engineering and Computer Science, University of Victoria, Victoria, BC, Canada; Department of Biomedical Engineering, University of Alberta, Edmonton, AB, Canada

**Keywords:** Missense mutation, Secondary structure, Protein backbone, Structure prediction

## Abstract

Human diversity often manifests through single nucleotide polymorphisms (SNPs). Among these, missense mutations, or SNPs that alter amino acids, can modify a protein’s three-dimensional (3D) structure. This impacts its function and can potentially elicit diseases or affect drug interactions. Thus, understanding protein single point mutations is crucial for precision medicine, as it helps tailor treatments based on individual genetic variations. As atomic locations can be susceptible to any number of changes that might or might not affect function, we focus on the secondary structure to provide concrete results on possible protein structural deformation that may occur from missense mutations.

We assess state-of-the-art structure prediction methods regarding backbone deformations caused by missense mutations. We categorize these deformations as **local, distant**, or **global** based on the proximity of structural changes to the mutation site. Our analysis utilizes a diverse dataset from the Protein Data Bank, comprising over 500 protein clusters with experimentally determined structures and documented mutations.

Our findings indicate that missense mutations can significantly affect the accuracy of structure prediction methods. These mutations often lead to predicted structural changes even when the actual secondary structures remain unchanged, suggesting that current methods overestimate the impact of missense mutations. This issue is particularly evident in advanced prediction algorithms, which struggle to accurately model proteins with stable mutations. We also found that the addition of low-performing prediction methods during structural analysis can positively impact the results on some proteins, particularly those with low homology. Furthermore, proteins that form complexes or bind ligands—such as membrane and transport proteins—are inaccurately predicted due to the absence of extra-molecular interaction data in the models, highlighting how missense mutations can complicate accurate structure prediction. All code and data are available at https://github.com/ivanpmartell/pdb-sam.

## Introduction

Variation in the human genome is commonly found via single nucleotide polymorphism (SNP) [7]. There are SNPs that occur in coding regions of the genome, causing translational effects by changing the amino acid sequence of proteins, and are known as *missense* mutations or non-synonymous SNP (ns-SNP) [41]. These mutations are often associated with changes in a protein’s ability to interact with other molecules, which can lead to one or more disease phenotypes [66, 39, 19, 17, 9]. It is also possible for these mutations to have an impact on the protein’s backbone and overall functionality by altering its structural integrity [71].

The three-dimensional structure of a protein is directly correlated with its function and is partly defined by its amino acid sequence [40]. Protein tertiary structure can be characterized mainly by the angles of certain backbone atoms. Ramachandran angles [51] are torsion angles *ϕ, ψ*, and *ω* that describe rotation of the protein backbone around the bonds between *N*-*C*_*α*_, *C*_*α*_-*C*^′^ and *C*^′^-*N* respectively. Common angle patterns found in protein backbones were first discovered by Pauling, Corey and Branson, with more patterns discovered afterward [22]. These patterns later became known as secondary structure elements and form the basis of the protein structure for self-assembly into its final conformation. This collection of secondary structure elements for a protein is known as the protein’s secondary structure. With a protein’s assigned secondary structure, its backbone atoms’ angles and distances can be inferred to obtain estimates of its sidechain atom locations.

Thus, a protein’s backbone and secondary structure contain essential structural information for its tertiary structure. Furthermore, the secondary structure of a protein can assist in the prediction of its functionality [63] and serve as an indicator for tertiary structure changes from missense mutations.

Secondary structure classification eliminates the atomic coordinate noise in side chains caused by atomic vibrations. This reduction in atomic-level detail aids in elucidating mutational effects without the confounding variations introduced by experimental and physical factors, thereby increasing the reliability of observed structural changes and removing atomic inconsistencies.

Recent studies [45, 38] have evaluated high-performing structure prediction models on missense mutations, but their analyses are limited to backbone-level assessments using *three-class secondary structure* (Q3). Distances between backbone atoms are used to determine backbone bonds, which can then be classified into specific secondary structure elements (SSEs) by assignment algorithms such as DSSP from Kabsch et al. [37]. As its name suggests, Q3 categorizes all SSEs into three structural classes—*α*-helix, *β*-sheet, and coil. However, as noted in the literature [37, 45], Q3 classification does not fully capture the complete structural complexity of the protein backbone. Furthermore, as existing methods approach the theoretical 90% accuracy limit for three-state predictions [76], efforts have shifted toward the more challenging task of *eight-class secondary structure* (Q8) prediction. Although the theoretical upper bound for Q8 accuracy is not well defined, current template-free methods achieve roughly 75% accuracy [56].

Evaluation of secondary structure prediction requires a consistent secondary structure assignment to use as a *gold standard*. However, most algorithms for protein secondary structure prediction have been found to produce differing results [6], especially in proteins with irregular conformations [80]. We selected DSSP [37], a pioneering algorithm for secondary structure assignment, as the gold standard because it has been extensively tested and utilized for machine learning tasks.

In this work, we investigate changes caused by missense mutations at three distinct scales:

- *Local* changes, which occur in the immediate vicinity of the mutated residue.
- *Distant* changes, which arise beyond the immediate neighborhood of the mutation site, but remain structurally linked to that specific region.
- *Global* changes, which can manifest anywhere in the overall protein structure, regardless of the mutation’s location.

We examine these distinct scales of backbone modification arising from a missense mutation to deepen our understanding of its impact on protein structure. The different vicinity scales allow us to learn how missense mutations affect the backbone structure within the experimental data, and highlight how prediction methods differ in determining the backbone structure after a mutation.

To the best of our knowledge, current structural prediction methods have not been evaluated on their ability to detect protein backbone structural changes induced by missense mutations at the local, distant, and global levels. To assess these structural predictions, we use experimental data to determine how a mutation affects a protein’s secondary structure. In this article, we examine the applicability and performance of nine state-of-the-art, methodologically diverse protein structure predictors and evaluate their capacity to distinguish local, distant, and global changes caused by missense mutations, using Q8. This procedure is shown in Fig. 1.

**Figure 1.**
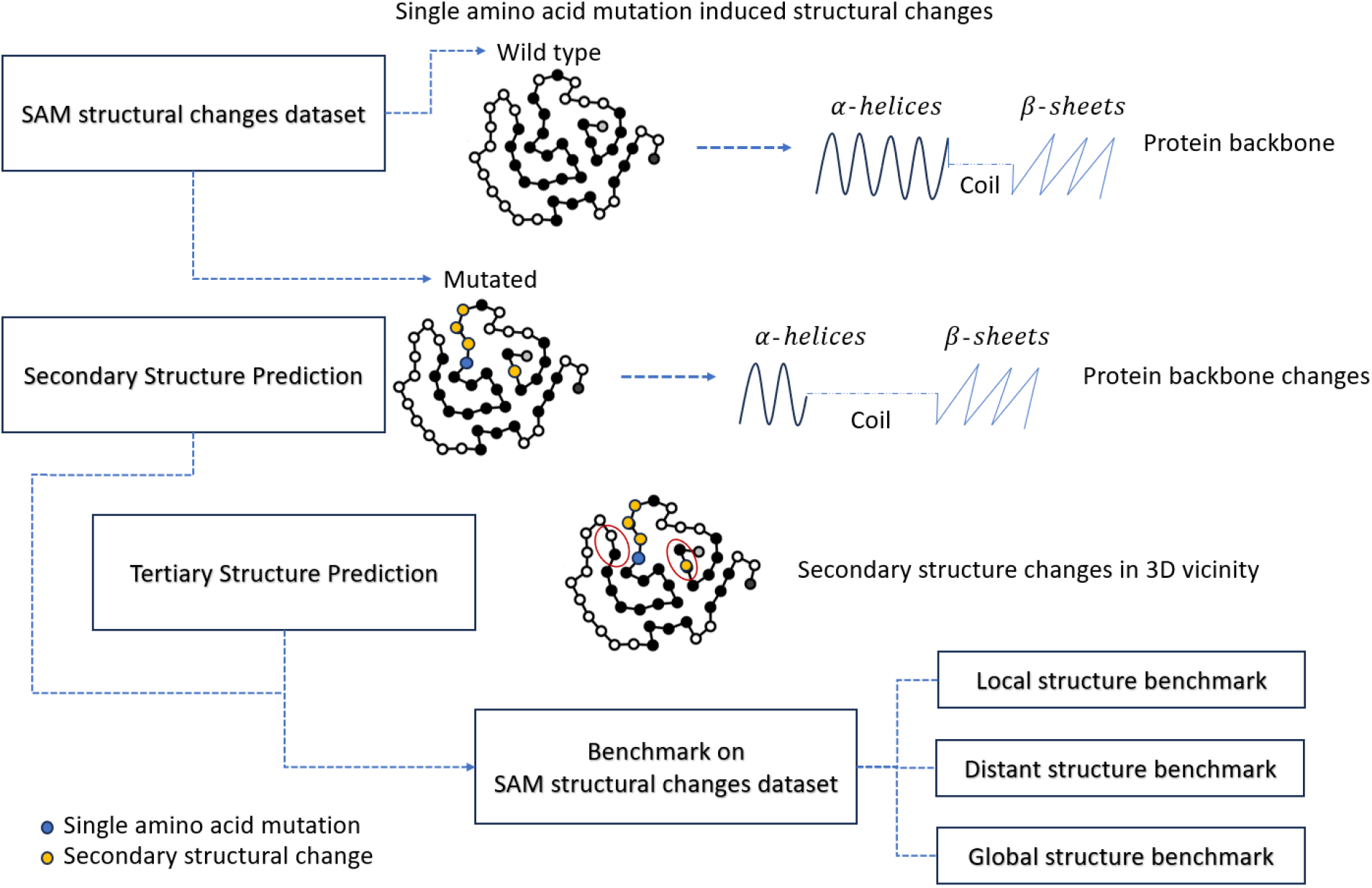
Protein secondary structure assessment. We assess both secondary and tertiary structure prediction methods on their eight-class secondary structure (Q8) prediction proficiency.

In addition, we extend our analysis to both secondary and tertiary structure prediction methods. Our focus on the protein backbone and its secondary structure elements is motivated by the strong correlation between backbone rigidity and protein thermostability [28, 64], as well as by their capacity to characterize the protein’s tertiary conformation without introducing atomic-level noise.

## Background

Protein structure is described through multiple differing levels of complexity. *Primary structure* refers to the linear sequence of amino acids, linked by peptide bonds, which make up the protein chain and dictates all subsequent levels of its structure. *Secondary structure* refers to the conformation of the protein backbone excluding the side chains. Regularly occurring secondary structures in proteins are called elements, e.g., *α* helices and *β* sheets. These secondary structure elements can vary widely in length, from as few as three to five residues, to over fifty residues. The connectivity between such secondary structure elements is often referred to as the protein topology. *Tertiary structure* is the overall three-dimensional shape of a protein chain, stabilized by physico-chemical interactions to its surrounding environment, and thus heavily influenced by it. Lastly, the *Quaternary structure* is the arrangement of multiple folded protein molecules in a multi-subunit complex.

Protein tertiary structure determines a protein’s overall shape, with its three-dimensional atomic arrangement tied to the precise spatial coordination of secondary structure elements. Because tertiary structure depends on a dynamic environment, its conformation is constantly fluctuating, albeit partially stabilized by interatomic bonds. The hierarchical model of protein folding supports this view, positing that pre-formed local secondary structures fold progressively into larger superstructures with native-like topology. Consequently, analyzing the protein backbone, its secondary elements, and overall topology can detect folding transitions and shape changes without requiring atomic-level resolution, which can be obscured through subtle alterations such as those caused by Brownian motion.

Significant advances in protein secondary structure prediction emerged from the Critical Assessment of Structure Prediction (CASP) competition. Over time, machine learning models became the primary predictors due to their improved accuracy, often leveraging co-evolutionary data for each protein. More recently, deep learning approaches have outperformed traditional machine learning methods, further increasing their prominence in the field [32, 62, 34].

We focus on locally installable protein secondary structure prediction software, prompted by the observation that many cloud-based tools lose support after a few years. As a recent review [32] notes, while most modern models achieve similar overall performance, they differ in Q8 predictions. With this in mind, we carefully selected publicly available top models for Q8 prediction, drawing from the reviewed methods and including additional models that have surpassed these leading tools. These models include Raptor-X Property [73], SSPro8 [44], SPOT-1D [29], SPOT-1D-Single [59], and SPOT-1D-LM [60].

We excluded models that are currently unavailable (including C8-SCORPION [77], and eCRRNN [79]) or require extensive training, particularly deep learning approaches lacking publicly available pre-trained models. Although historically significant since the inception of secondary structure prediction, PsiPred was excluded due to its Q3-only focus. Similarly, other Q3 prediction software not included in this study are PHD [53], SOPMA [27], SPINE-X [25], SPARROW [10], and JPRED [21].

We also did not consider certain Q8 predictors—namely SPIDER3 [31] and SPIDER3-Single [30]—because SPOT-1D and SPOT-1D-Single outperform them. Additionally, we excluded tools that did not have an accessible prediction capability for novel amino acid sequences, including MUFOLD-SS [24].

Deep learning algorithms have led to unprecedented progress in predicting tertiary protein structures directly from their amino acid sequences. A major breakthrough occurred in 2021 with AlphaFold2 [36], which achieved performance comparable to experimental methods, such as X-ray crystallography and NMR spectroscopy, for proteins with numerous homologs. Building on this, current deep learning approaches [48, 8, 42, 75] infer a protein’s 3D structure from its evolutionary relationships, identified through various techniques.

Ongoing efforts focus on creating highly accurate structure prediction methods for all types of molecules. With computing capabilities continuing to increase, machine learning models have become increasingly more feasible to train with larger amounts of data and parameters, which produce predictions for increasingly larger and more complex molcules [36, 1]. This facilitated the use of Alphafold’s deep learning methodology for subsequent models, with an emphasis on the exploration into their functionality.

For AlphaFold2 and its derivatives [36, 48, 8], homology is determined by comparing a protein’s amino acid sequence to known sequences via multiple sequence alignment (MSA). In contrast, language models like ESMFold [42, 75] capture homology during their training process, effectively encoding evolutionary information in their parameters.

The tertiary structure prediction methods we analyze were chosen for their outstanding performance and their distinct predictive approaches, despite all employing deep learning. These methods are AlphaFold2, ColabFold [48], ESM-Fold [42], and RGN2 [47]. The predictions from these methods contain confidence scores, which are calculated as the predicted Local Distance Difference Test (pLDDT) [36]. This metric measures the confidence in the local structure of a protein on a per-residue basis. It was originally developed for use in AlphaFold, but was also used by subsequent methods.

Due to challenges in understanding how deep learning models generate their results, additional investigation into the capabilities and limitations of these models is still necessary [14, 50]. This is particularly evident in mutational data, where a single mutation could substantially change the protein structure.

### Selected prediction methods

In this section, we describe the structure prediction methods selected for evaluation. Most of these approaches leverage a protein’s homology data, incorporating evolutionary information through multiple sequence alignment (MSA) against a database of known proteins. Two commonly used tools for this purpose are PSI-BLAST [4] and HHBLITS [52]. PSI-BLAST identifies similar proteins in the database and produces a position-specific scoring matrix (PSSM) from the resulting MSA, while HHBLITS employs a Hidden Markov Model to align the query protein against UniProtKB, providing a measure of sequence similarity.

We begin by introducing the secondary structure prediction methods. Further details on these methods are given in Section 8 of Supplementary Information.

- Raptor-X Property [73] uses a deep learning model that combines two powerful techniques: Conditional Random Fields and Convolutional Neural Networks. This model can capture complex patterns and correlations among protein sequences and structures.
- SSPro8 [44] combines PSI-BLAST, a Bidirectional Recurrent Neural Network (BRNN), and structural similarity to produce accurate results. SSPro8 consists of three steps: 1) align amino acid sequences and
- compute profile probabilities with PSI-BLAST, 2) use 100 BRNNs trained on data to predict probabilities for each structure class, and 3) use amino acid sequence-based structure similarity to refine predictions from the BRNNs ensemble.
- SPOT-1D [29] uses an ensemble of nine models composed of bidirectional recurrent neural networks and residual networks. These models are trained on evolutionary profiles derived from PSI-BLAST and HHBLITS, as well as predicted contact maps from SPOT-Contact and the physicochemical properties of amino acids.
- SPOT-1D-Single [59] employs three neural network models that integrate residual networks and bidirectional recurrent neural networks. In contrast to SPOT-1D, it does not depend on evolutionary information from multiple sequence alignments or contact maps.
- SPOT-1D-LM [60] leverages an ensemble of deep learning language models–ESM-1b [42] and ProtTrans [23]– to encode evolutionary information from protein amino acid sequences. The outputs of these language models are then processed by a deep learning architecture similar to SPOT-1D-Single.

We evaluated the following tertiary structure prediction methods. Further details concerning these methods are given in Section 9 of Supplementary Information.

- AlphaFold2 [36] is a neural network model integrating sequence and structural modules, leveraging MSAs as evolutionary features. Its predictions are iteratively refined in a process known as *recycling*. AlphaFold2 surpassed other methods in the CASP14 competition, yet its performance diminishes for proteins with few intra-chain (homotypic) contacts or when fewer than 30 aligned amino acid sequences are available [36].
- ColabFold [48] refines AlphaFold2 by using MMseqs2 [65] for more efficient MSA generation. It achieves performance comparable to AlphaFold2 while running 40 to 60 times faster.
- ESMFold [42] employs the ESM-2 deep learning language model, trained on millions of protein amino acid sequences, to capture evolutionary patterns and structural features. It then uses a folding module that iteratively refines sequence and pairwise representations, ultimately predicting atomic coordinates and their associated confidences. Like AlphaFold2, ESMFold utilizes recycling to enhance its predictions; however, unlike AlphaFold2, it does not rely on MSA features.
- RGN2 [47] employs AminoBERT [15], a language model that encodes evolutionary amino acid sequences into feature vectors. A recurrent network and Frenet-Serret structural frames [33] then decode these vectors into the protein’s backbone geometry. The resulting structure can be further refined using an energy-based protocol. RGN2 performs effectively even when no evolutionary information is available.

### Terminology

Let **A** = {*A, C, D, E, F, G, H, I, K, L, M, N, P, Q, R, S, T, V, W, Y*} be a finite set containing the alphabet of standard amino acids as defined by residue or side chain.

A **protein sequence** over **A** is a finite sequence of amino acids **S** = [*a*_1_, …, *a*_*n*_] of length *n*, where *a*_*i*_ ∈ **A**, for all 1 ≤ *i* ≤ *n. a*_1_ is the N-terminus residue and *a*_*n*_ is the C-terminus residue. For each amino acid *a*_*i*_ ∈ **S** at position *i* in **S**, there exists two 3D real-valued coordinates 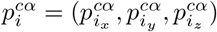 and 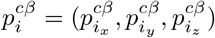, where 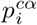 and 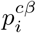 are associated to *C*_*α*_-atoms and *C*_*β*_-atoms respectively of the amino acid in Angstrom (Å) units. The list of coordinates for all amino acids in **S** is called the **3D structure** of **S**.

For 1 ≤ *i* ≤ *j* ≤ *n*, **S**_*ij*_ = [*a*_*i*_, *a*_*i*+1_, …, *a*_*j*_] is called a **protein substring** of length *L* = *j* − *i* + 1 of **S** from the amino acid at position *i* to *j*. The lists of the corresponding *C*_*α*_-coordinates and *C*_*β*_-coordinates associated with the amino acids in sequence **S**_*ij*_ are called 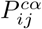 and 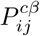, respectively.

For the protein sequence **S** each amino acid *a*_*i*_ is assigned a secondary structure *r*_*i*_ resulting in **R** = [*r*_1_, *r*_2_, …, *r*_*n*_]. Similarly, for a substring **S**_*ij*_ = [*a*_*i*_, *a*_*i*+1_, …, *a*_*j*_] the secondary structure is then given by **R**_*ij*_ = [*r*_*i*_, *r*_*i*+1_, …, *r*_*j*_]. As we investigate Q8 using DSSP, each SSE in **R**_*ij*_ must be an assignment *r*_*l*_ ϒ_8_, where ϒ_8_ ={𝒞, ℋ, ℰ, 𝒢, ℐ, 𝒯, 𝒮, ℬ is the set of possible SSE classes in DSSP.

If the amino acid *a*_*m*_ ∈ **S**_*ij*_, *i* ≤ *m* ≤ *j* is mutated to *â*_*m*_, this results in the modified sequence *Ŝ*_*ij*_ and the corresponding *C*_*α*_-structure 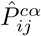 and *C*_*β*_-structure 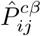.

For any amino acid *a*_*k*_ ∈ **S**, we define its **primary structure vicinity** (1D vicinity) 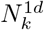 as the subsequence of amino acids surrounding *a*_*k*_. Formally, 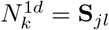 where 1 ≤ *j* = *k* − ϵ_1*d*_ ≤ *l* = *k* + *ϵ*_1*d*_ ≤ *n* and 0 ≤ *ϵ*_1*d*_ ≤ 20. This range was chosen to reflect the average length of secondary structure elements, which most commonly span from 10 to 40 amino acids [61]. See Fig. 2.A for a visual representation of the 1D vicinity.

**Figure 2.**
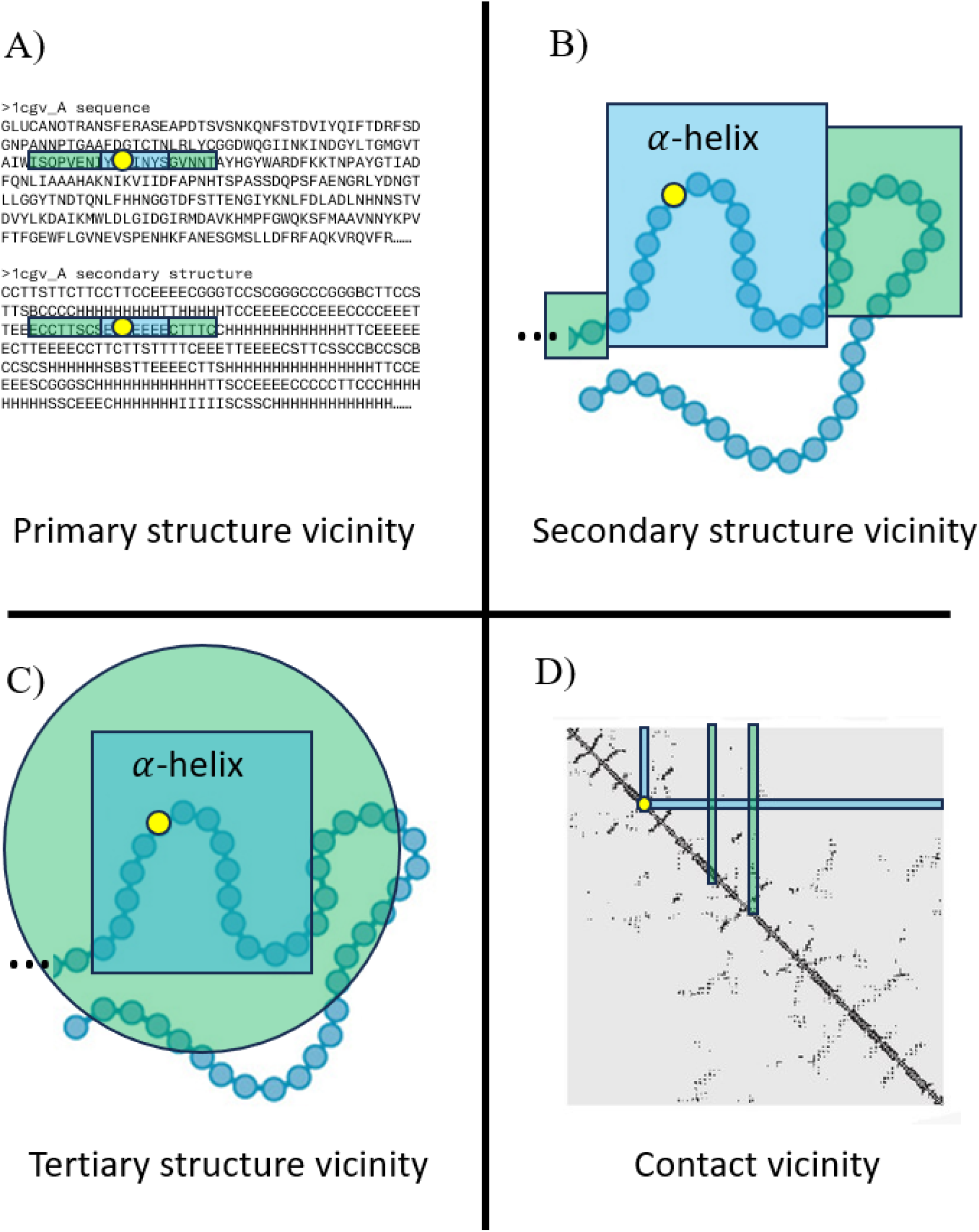
Vicinity Measurements. Measurement of vicinity from a missense mutation location shown with a yellow circle. The missense mutation is associated with an *α*-helix, shown in blue. Vicinity is shown in green. A) 1D vicinity. 2D vicinity. C) 3D vicinity. D) contact vicinity

The **secondary structure vicinity** (2D vicinity) 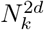 of an amino acid *a*_*k*_ ∈ **S** is defined as the concatenation of three subsequences: 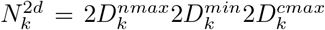, where 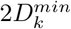, the minimal 2D vicinity, 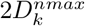, the N-terminus extension and 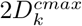, the C-terminus extension are defined as follows.

- The **minimal 2D vicinity** is defined as 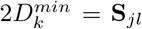 where 1 ≤ *j* = *k* − *ϵ*_2*d*_ ≤ *l* = *k* + *ϵ*_2*d*_ ≤ *n* with 0 ≤ *ϵ*_2*d*_ ≤ 10. This minimal vicinity includes a fixed number of amino acids on both sides of *a*_*k*_. We extend this minimal vicinity to capture the largest possible contiguous secondary structure blocks at each terminus.
- **N-terminus extension** 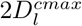: Consider secondary structure *r*_*j*_, assigned to the leftmost amino acid *a*_*j*_ in **S**_*jl*_. Then 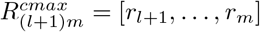 is called an *nmax*-block if
  1. 1 ≤ *i* = *k* − *γ*_2*d*_ *< j*,
  2. *r*_*i*_ = … = *r*_*j*−1_ = *r*_*j*_, and
  3. *r*_*i*−1_ ≠ *r*_*j*_.
  Then, 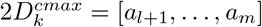 is the largest contiguous subsequence on the N-terminus side of 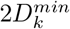 that has the same secondary structure as *a*_*j*_.
- **C-terminus extension** 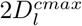: Similarly, consider secondary structure *r*_*l*_, assigned to the rightmost amino acid *a*_*l*_ in **S**_*jl*_. Then 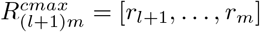 is called *cmax*-block if
  1. *l < m* = *k* + *γ*_2*d*_ ≤ *n*,
  2. *r*_*l*+1_ = … = *r*_*m*_ = *r*_*l*_, and
  3. *r*_*m*+1_≠ *r*_*l*_.
  Then, 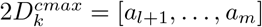 is the largest contiguous subsequence on the C-terminus side of 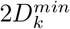 that has the same secondary structure as *a*_*l*_.

We set *γ*_2*d*_ = 30, defining the maximum possible extension at each terminus.

Therefore, every amino acid in an extension must maintain the same secondary structure as their respective minimal vicinity boundary amino acids *a*_*j*_ or *a*_*l*_; if not, the extension ceases where the structure changes. The resulting secondary structure vicinity is 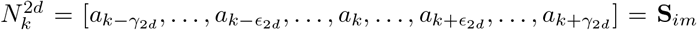. The length of 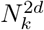 is constrained by *γ*_2*d*_ = 30 to achieve vicinity lengths comparable to the other vicinity types. However, the maximum vicinity length for 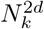 is rarely reached due to the typically small span of secondary structure elements. We also enforce a lower bound *ϵ*_2*d*_ to prevent the vicinity from becoming too small. See Fig. 2.B for a visual representation of the 2D vicinity.

For any amino acid *a*_*k*_ ∈ **S**, its **tertiary structure vicinity** (3D vicinity) 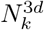 is a collection of subsequences of **S**, defined by the set of positions of the amino acids in its 3d neighborhood {*j* | *d*_*α*_(*k, j*) ≤ *ϵ*_3*d*_}, with *ϵ*_3*d*_ = 13 and *d*_*α*_ is a distance metric given by Eq 1. A visual representation can be seen in Fig. 2.C.

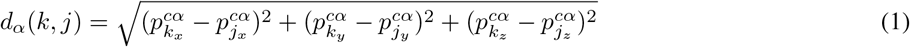

Analogously, we define **contact vicinity** 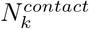 for any amino acid *a*_*k*_ ∈ **S** as the collection of subsequences of **S**, defined by the set of positions of the amino acids in its contact vicinity {*j* | *d*_*β*_(*k, j*) ≤ *ϵ*_*contact*_}, where *ϵ*_*contact*_ = 8 and *d*_*β*_ is a distance metric given by Eq 2. A visual representation can be seen in Fig. 2.D.

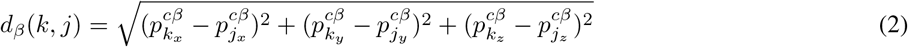

The maximum vicinity length thresholds for 3D and contact vicinities are set to match previous definitions. The threshold for 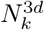, *ϵ*_3*d*_ is based on a previously established threshold by Alphafold2 [45], while *ϵ*_*contact*_ for 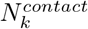 aligns with the accepted definition of contact prediction [55].

A single amino acid mutation can trigger a structural change within a protein. We categorize the structural changes into three categories for each of the aforementioned vicinities. A **local** structural change occurs if a mutation at position *l* alters the secondary structure within its respective vicinity 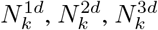, or 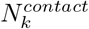. A **distant** structural change occurs if a mutation at *l* modifies the structure outside its respective vicinity (i.e.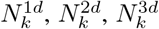, or 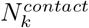) but not within the vicinity. Finally, a **global** structural change occurs if the altered secondary structure is neither confined within nor solely outside its respective vicinity.

When a mutation induces a structural change, we refer to it as *disruptive*. In contrast, if no secondary structural change occurs, the mutation is considered *stable*. The structural changes described above are illustrated in Fig 3.

**Figure 3.**
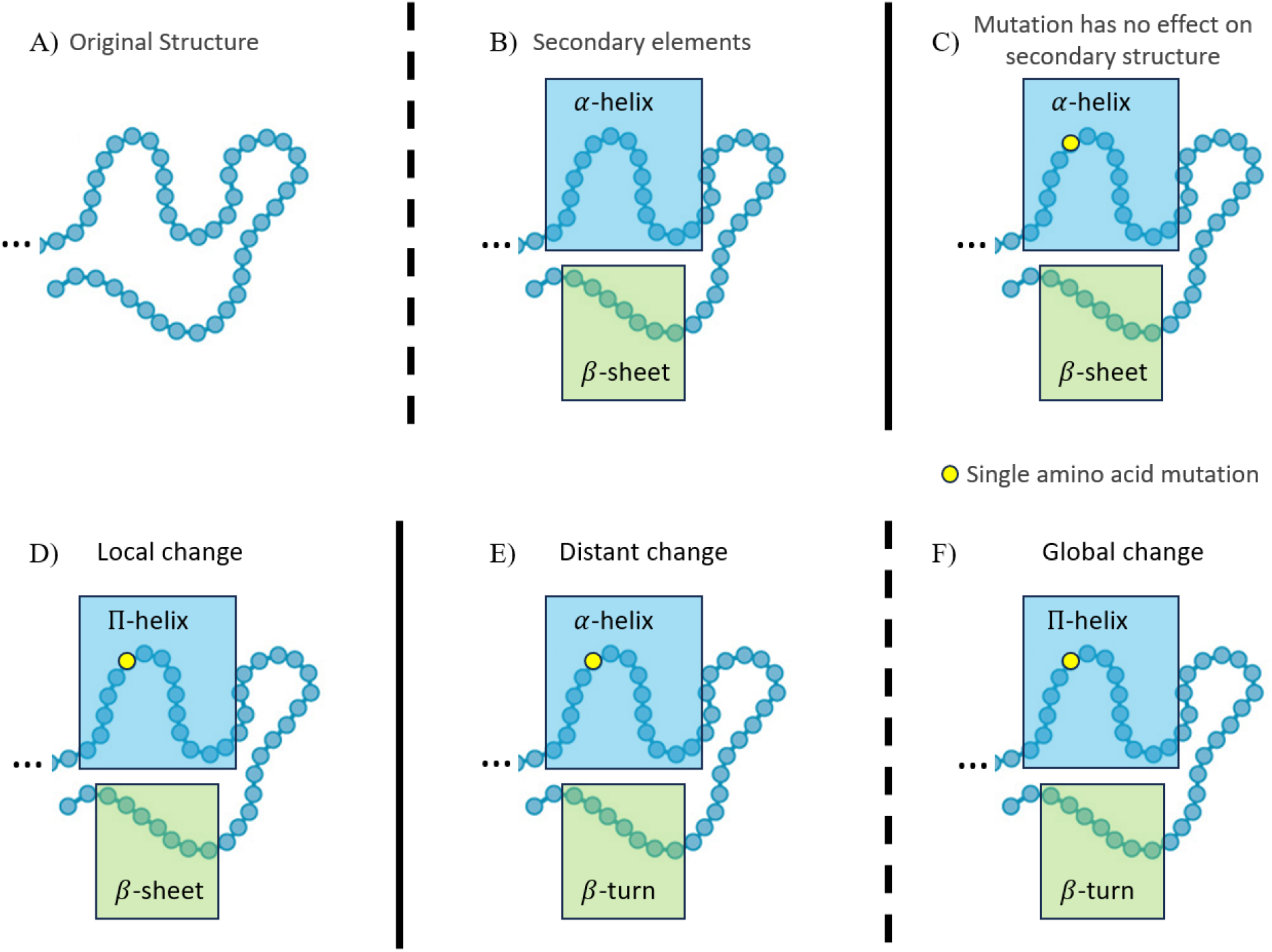
Types of Backbone Changes. Measuring backbone changes in proteins requires pinpointing specific locations within their secondary structures, relative to the mutation site. This allows us to observe how a mutation impacts the protein’s backbone. Yellow circles indicate the amino acid mutation location. Blue regions show the secondary structure in the mutated region. Green regions contain the amino acids that are part of the vicinity. A) Original protein backbone structure. B) Secondary elements in the protein backbone. C) No structural change due to mutation. D) Local structural change due to mutation. E) Distant structural change outside the local structural vicinity of the mutation. F) Global structural change occurring anywhere in the protein backbone.

## Materials and methods

In order to assess the performance of our selected structure prediction methods on protein backbone changes induced by missense mutations, we created a mutation dataset with experimentally obtained structures. In the following section, we describe how we created our benchmark dataset as well as caveats on this data regarding the assessed methods.

### Data acquisition and processing

Protein sequences (PDB_SEQRES.TXT) and their experimentally derived 3D structures (PROTEIN.CIF) were obtained from the Protein Data Bank (PDB) as of April 2023. We excluded non-protein sequences and duplicates, retaining only proteins composed of the standard 20 amino acids. Consequently, any sequences containing ambiguous amino acids were also removed.

We clustered the protein sequences using CD-HIT [26] with a 99% sequence similarity threshold, producing groups of mutated proteins. Within each cluster, we performed multiple sequence alignments using Clustal Omega [57] to identify amino acid substitutions and discard incomplete sequences. After alignment and filtering, each cluster retained only full-length sequences of uniform length.

However, while retrieving the structural data files for these sequences, we sometimes encountered structures with missing amino acids, resulting in gaps where the atoms’ locations are inconclusive. Such gaps can obscure the true effects of missense mutations, as the 3D structures may not fully represent the corresponding protein sequence associated to the structure. To ensure that all relevant atomic positions are accounted for, we excluded any proteins whose structure files contained these gaps. This approach guarantees that the mutation effects that we analyze are accurately represented.

After this preprocessing step, we applied DSSP to assign secondary structures to each protein sequence using their corresponding experimental structures. As resulting protein sequences had uniform length within a cluster and differ through missense mutations, the proteins could be considered aligned. Wild-type and mutated sequences were identified through mutation extraction that is detailed in Section 3 of Supplementary Information. The identification procedure follows the same logic as Weblogos [18], which display the most common amino acid at each position of the sequence as the largest symbol in a figure.

We ran each method on our dataset and normalized the results to ensure consistent Q8 predictions, as different methods may use distinct symbols for identical secondary structure classes. After completing these steps, we obtained the final preprocessed dataset used for evaluating the methods. A summary of this process is illustrated in Fig. 4.

**Figure 4.**
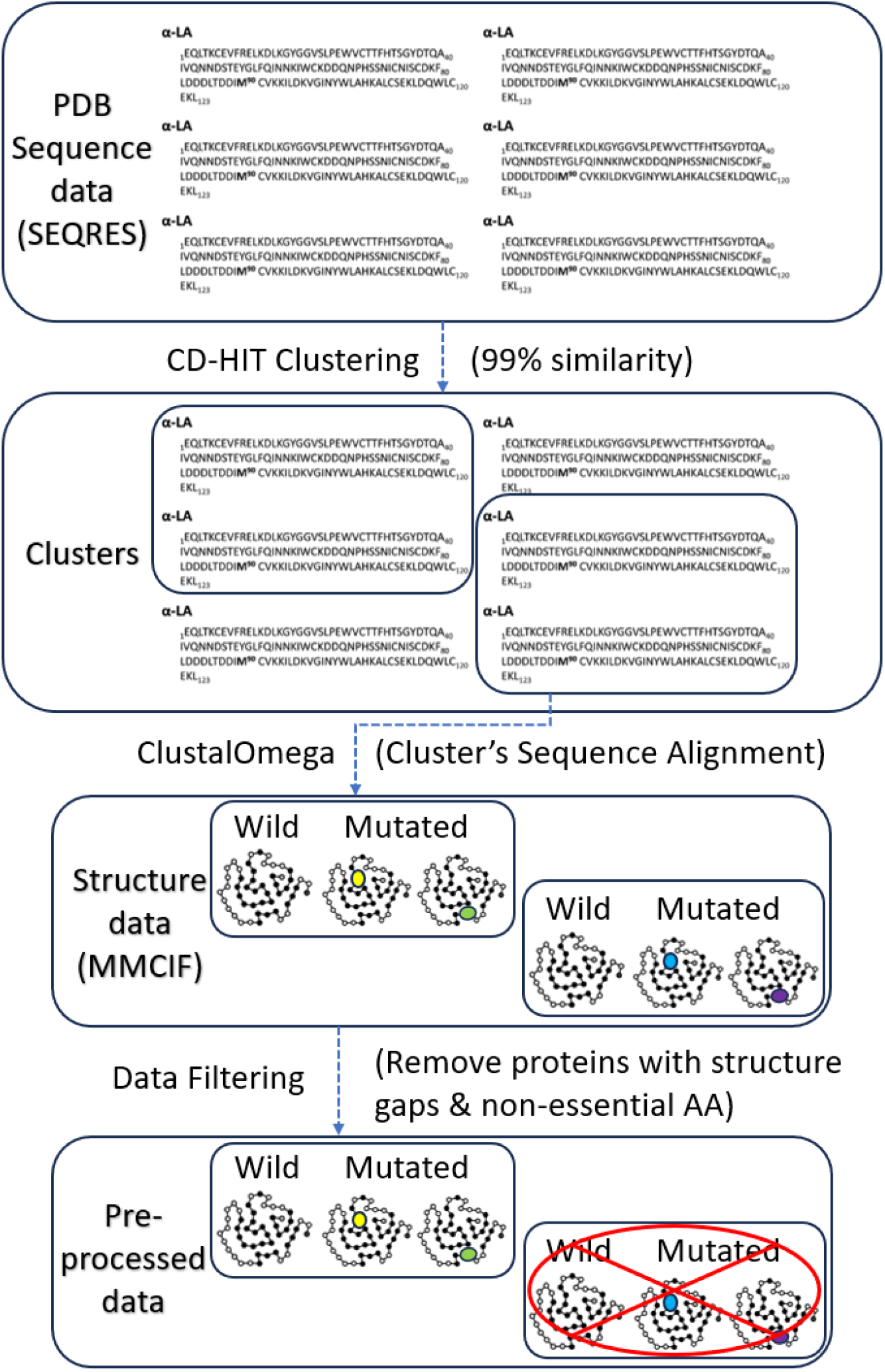
Data processing. Top to bottom: Begin with “SEQRES’ data containing PDB molecular sequences (keeping proteins only). Cluster the sequences by 99% similarity using CD-HIT. Align each cluster’s sequences using Clustal Omega. Finally, filter the data to ensure all amino acids appear in their associated PDB structure files.

### Dataset statistics

Our preprocessed dataset comprises 579 clusters encompassing 1, 414 proteins. It serves as a simple mutational dataset, where at most 1% of the amino acids are mutated and an average length of 238 amino acids. Further filtering of the dataset ensures a dataset containing single amino acid mutations only. The protein sequences range from 100 to 858 amino acids in length. Any proteins longer than 1, 024 amino acids were excluded, as several prediction methods cannot handle such length.

Each cluster may include multiple single amino acid mutations, depending on the number of proteins it contains. Approximately 60% of the clusters have just one mutation, while the remainder is evenly split—between those with two mutations and those with three or more. A wide range of sequence length is represented across all clusters, as illustrated in Fig. 5.

**Figure 5.**
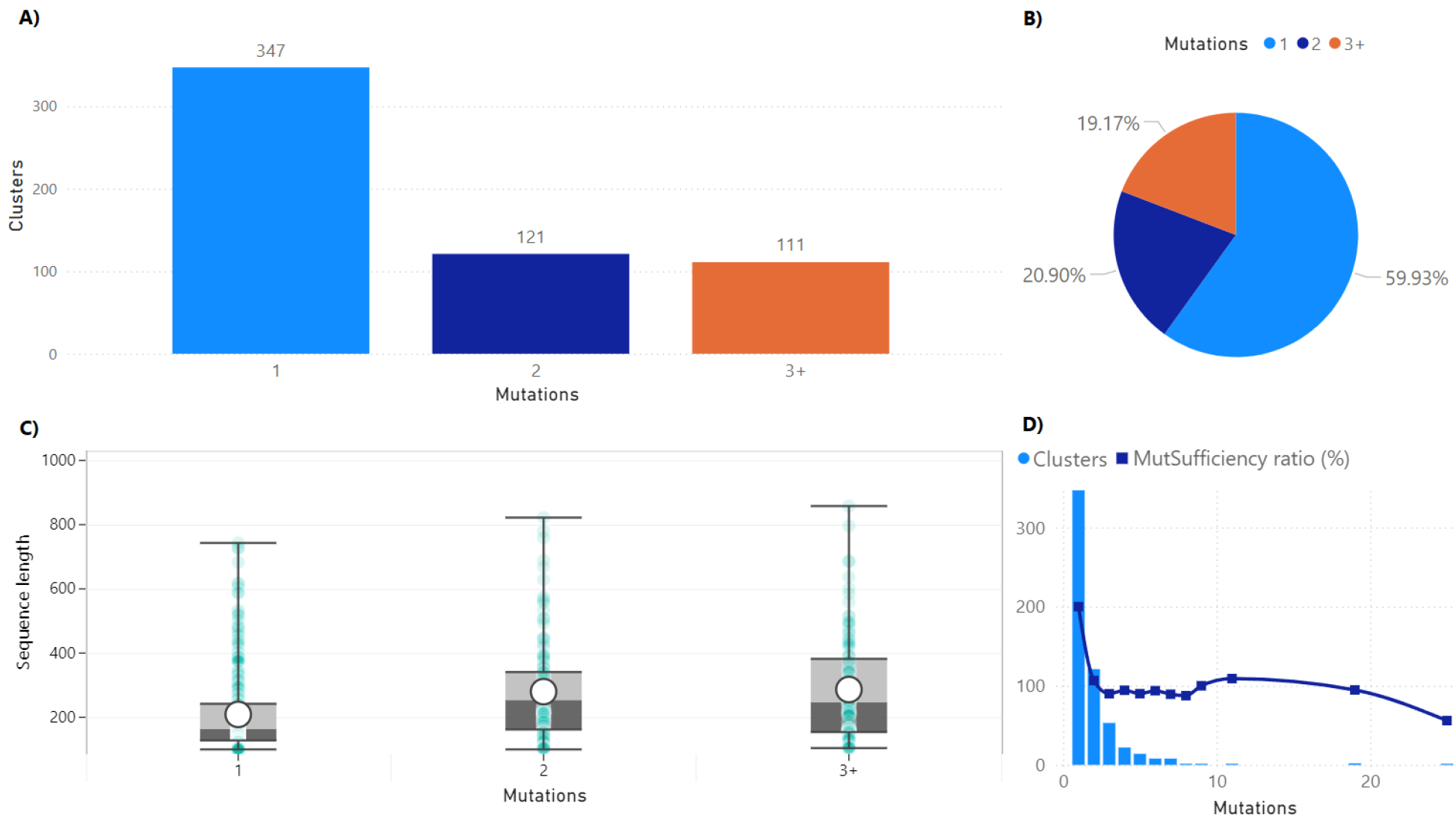
Protein mutation statistics. A) Amount of clusters containing 1, 2, or 3+ protein mutations. B) Percentage of clusters in the dataset for certain mutations. C) Protein sequence lengths in clusters with 1, 2, or 3+ protein mutations. We can see an even spread of sequence lengths among all clusters regardless of mutations D) Amount of clusters with a certain amount of mutations and mutation sufficiency ratio as a percentage.

Because the clusters were formed at 99% sequence similarity, individual proteins within a cluster may carry more than one amino acid mutation. To identify clusters containing only a single unique mutation per protein, we applied the *Mutation Sufficiency* (`MutSufficiency`) metric defined in Eq. 3. This metric uses the ratio of the total number of mutations to the total number of proteins in a cluster (recall that each protein has at least one mutation).

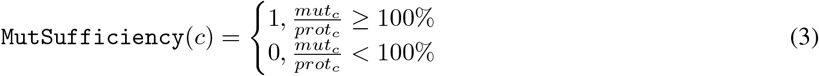

where *mut*_*c*_ is the total number of mutations in cluster *c* and *prot*_*c*_ is the total number of proteins in *c*. After applying the Mutation Sufficiency metric to filter the clusters, we retained 542 clusters and 1, 291 proteins, with an average length of 226 amino acids. (See Fig. 6).

**Figure 6.**
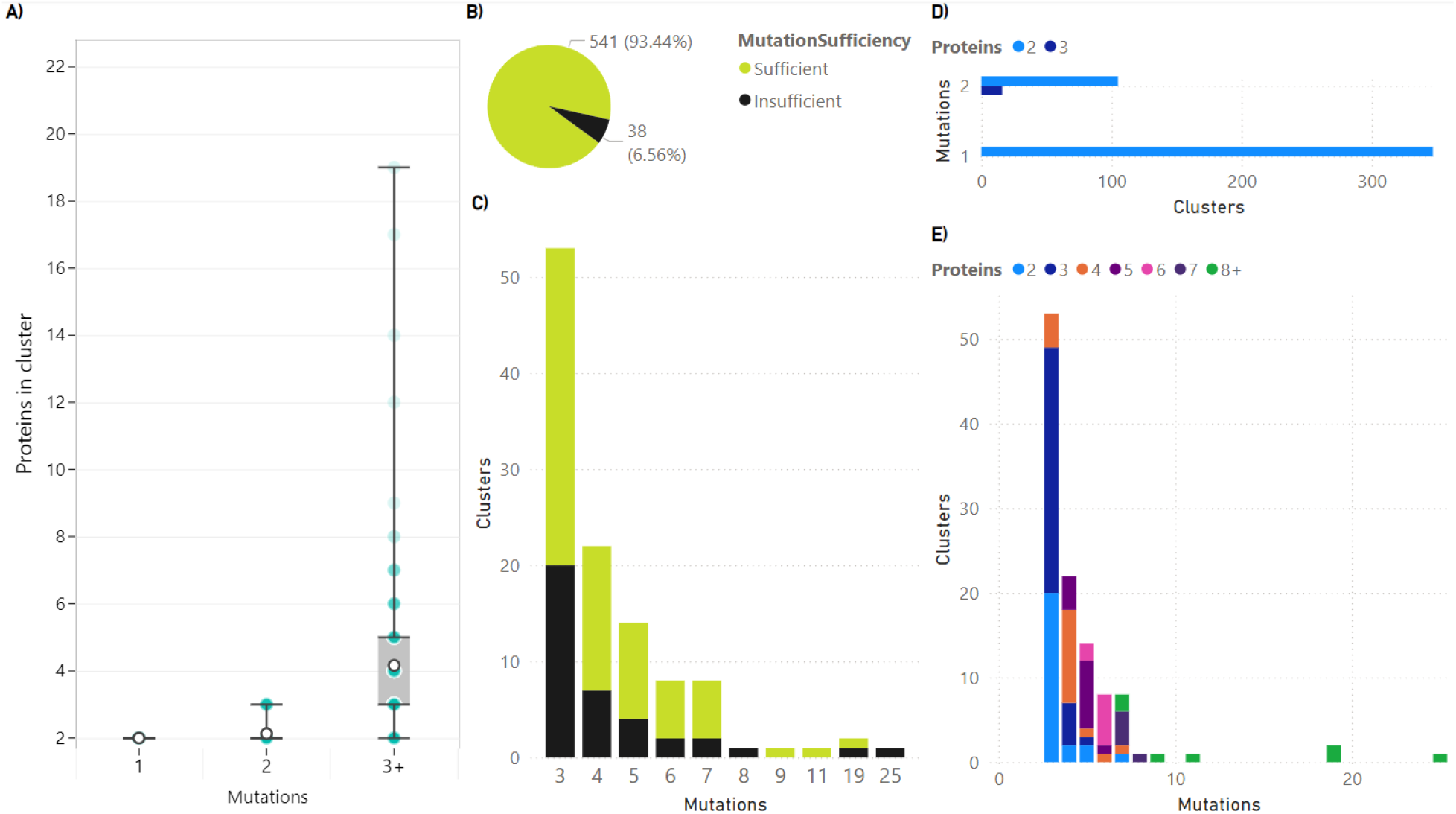
Protein mutation sufficiency statistics. All mutation insufficient clusters contain 3 or more mutations. Clusters were separated according to their amount of proteins from 2 to 8 or more (8+). A) Bar plot showing the mutations that occur for certain amount of proteins in a cluster. White circle represent the mean value. The box indicates the 25th and 75th percentiles. The transparency of the gray circles indicate the amount of clusters (more transparent, more clusters with that amount of proteins). B) Percentage and values of mutation sufficient and insufficient clusters. C) Amount of clusters containing a specific amount of mutations. D) Clusters with 1 or 2 mutations (All of these clusters are mutation sufficient). E) Detailed graph of proteins and mutations for clusters

Most of the removed clusters from the Mutation Sufficiency filtering were those with three or more mutations per cluster, as well as those with longer protein sequences and higher mutation rates. These changes are detailed in Section 4 of Supplementary Information. This filtration primarily removed mutational outliers, where long protein sequences carried an exceptionally high proportion of mutations. As we delve into single amino acid mutations, the Mutation Sufficiency filtering was applied to remove any clusters with an insufficient amount of mutations arising from possible duplicate mutations.

We were also interested in any evolutionary structural information found in our protein dataset, as well as functional annotations that this evolutionary information can provide for the proteins. For this task, we utilized structural homology datasets, which contain a collections of proteins that are structurally similar due to shared ancestry. These datasets identify conserved regions, and predict the function of unknown protein sequences based on known ones from experimental data.

SCOP [5] (Structural Classification of Proteins) and CATH [58] (Class, Architecture, Topology, Homologous super-family) are databases that categorize proteins based on their structural and evolutionary relationships. Applying these classifications to our dataset reveals that not all proteins have matching entries: SCOP lacks data for about 25% of our clustered proteins, and CATH is missing data for approximately 15%.

Among the proteins with SCOP data, more than 98% are globular, around 1.5% are membrane proteins, and about 0.5% are fibrous proteins. In the CATH classifications, roughly 43% feature a mixture of alpha and beta secondary structures, about 36% have predominantly beta (sheet) structures, around 20% primarily contain alpha (helical) structures, and the remaining 1% exhibit few secondary structures.

Properties shared by more than 50% of our proteins primarily include top-level SCOP and CATH classifications, as well as common protein functions such as “binding’. The removal of these prevalent properties is done to minimize bias from frequently appearing characteristics in our subsequent analyses.

Each mutation vicinity differs in length due to the varying number of amino acids it encompasses. The contact vicinity is the longest, followed by the 1D and 3D vicinities. The 2D vicinity is the shortest, averaging at least 10 fewer amino acids than any other vicinity type.

These protein and mutation statistics are illustrated in Fig. 7.

**Figure 7.**
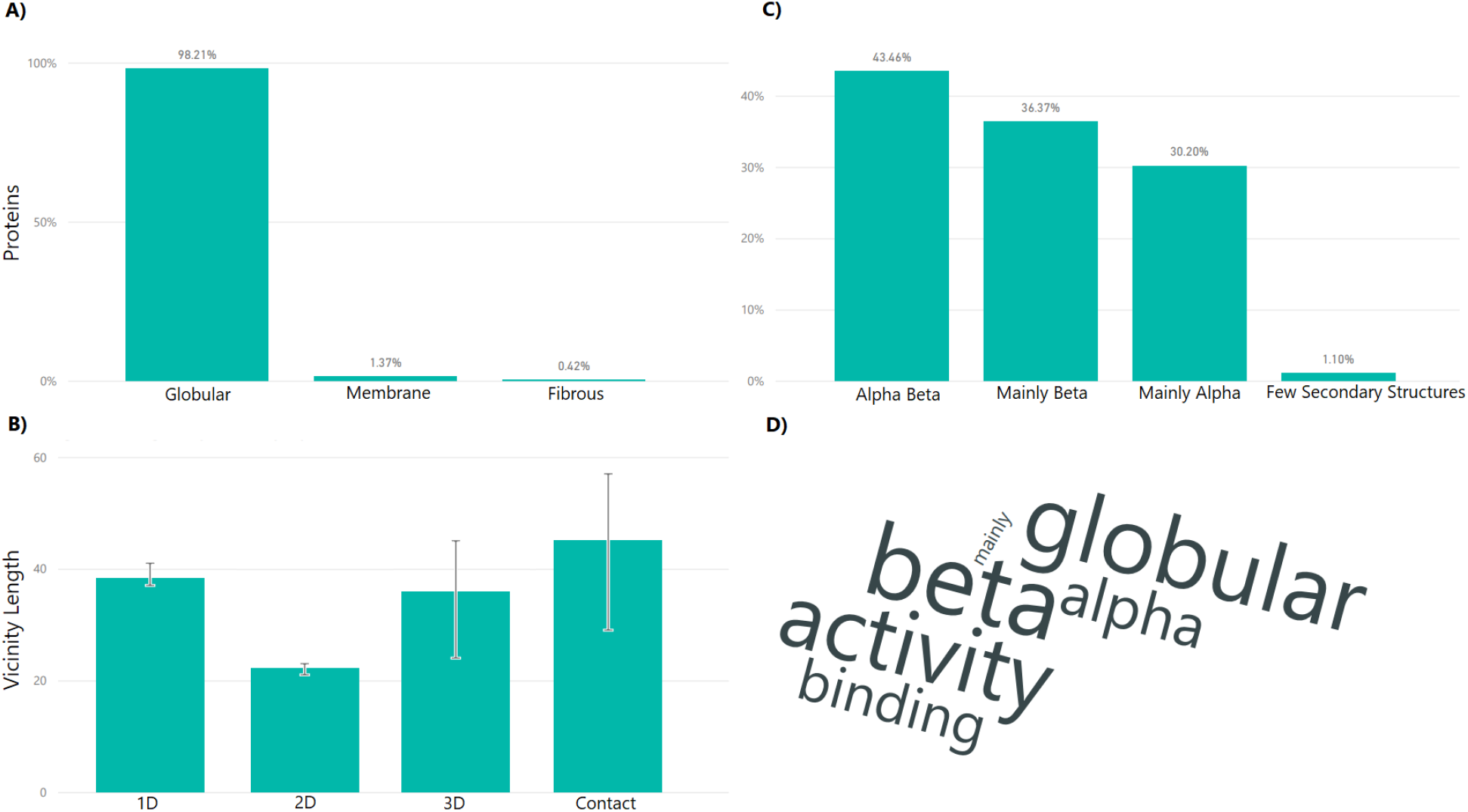
Protein statistics. Percentage proteins containing SCOP (A) and CATH (B) top level classifications in our data. C) Average vicinity length according to the mutation vicinity type. D) Most common protein properties in our dataset.

### Dataset limitations

Some of the original training data used by these methods are included in our dataset, introducing a positive performance bias that cannot be fully mitigated due to the limited availability of experimentally obtained single-amino acid mutation structures. To gauge the extent of the data overlap for each method, even though most methods do not provide the respective exact training dataset, we assembled a dataset closely resembling their training data by following the published methodologies.

Raptor-X Property was originally trained using a dataset described in Wang et al. [74], which included protein chains resolved at better than 2.5 Å, with less than 30% protein sequence identity, lengths between 50 and 700 residues, and no chain discontinuities. That dataset contained approximately 5,600 proteins.

To approximate these conditions, we selected the May 2012 CullPDB dataset (“cullpdb_pc30_res2.5_R1.0_d120428_chains9175’). We filtered it by length and excluded any proteins published after 2010, using publication dates obtained from the Protein Data Bank Japan ^1^. This process yielded 5, 972 proteins, among which 40 clusters and 40 individual proteins overlap with our dataset.

SSPro8 was originally trained on a CullPDB dataset from the PISCES [72] server, utilized for culling sets of protein sequences from the Protein Data Bank (PDB) based on sequence identity and structural quality criteria. The resulting dataset (Cullpdb_pc30_res2.5_ R1.0_d100716’), contained roughly 8,000 high-resolution protein structures. Similar to Raptor-X Property, we used the “cullpdb_pc30_res2.5_R1.0_d120428_chains9175’ dataset as the closest available resource. After discarding protein sequences shorter than 50 residues or longer than 1,500 residues and removing proteins published after 2011, we obtained a set of 8, 039 proteins. Of these, 45 proteins—distributed across 45 clusters—overlap with our dataset.

SPOT-1D utilizes a dataset of around 10,000 proteins. This dataset was based on the data used for SPIDER3-Single, a method that precluded SPOT-1D, which its methodology was upgraded to become SPOT-1D. This dataset has 38 clusters and 38 proteins in common with our dataset.

SPOT-1D-Single and SPOT-1D-LM utilize the same dataset which is publicly and readily available ^2^. Their dataset has 39119 proteins, which has 114 clusters and 126 proteins in common with our dataset.

Alphafold2 and ColabFold use a training dataset derived from PDB with proteins published before 30 Apr. 2018. This dataset contains 139,417 proteins with 1174 proteins in common with our dataset covering 503 clusters.

Similarly, ESMFold uses a training dataset derived from PDB with proteins published before 30 Nov. 2022. This dataset contains 198,618 proteins which cover 1, 371 proteins and 570 clusters in our dataset.

RGN2 was trained on the “astral-rapid-access-1.75.raf’ dataset, which includes 87, 061 proteins. Within our dataset, this training set overlaps with 269 clusters and 609 proteins.

Additionally, RGN2 predictions omit the last two residues of each protein sequence. To align with the lengths used by other methods, we appended two coil secondary structures to the end of each prediction. Comparing results with and without this adjustment (Fig. 8) shows a slight overall performance increase, likely because most protein termini consist of coils.

**Figure 8.**
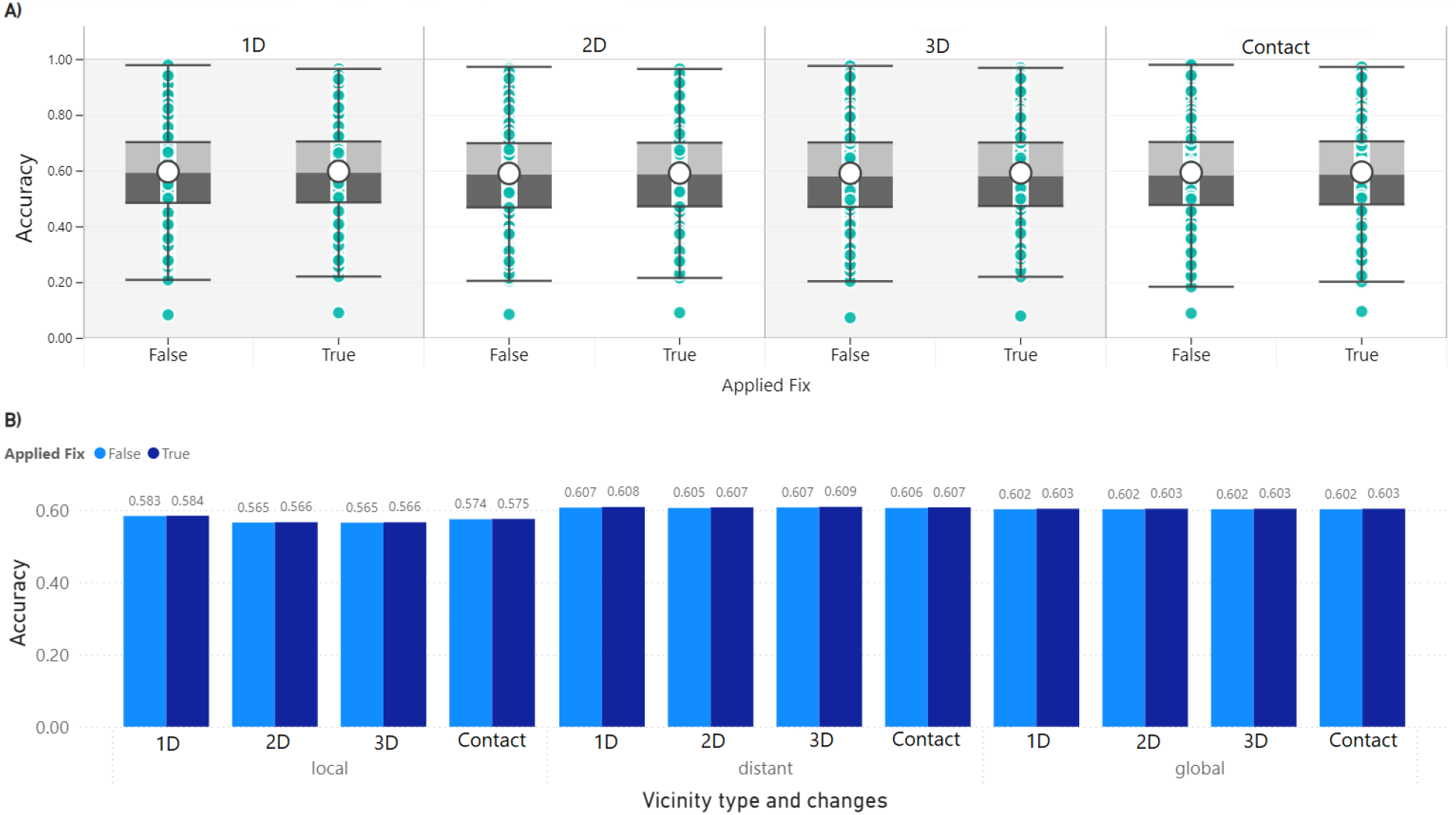
Adjustment to the RGN2 secondary structure prediction. Difference in performance with (True) and without (False) adjusting for the length. A) Box plot showing performance of RGN2 for each mutation vicinity with boxes spanning from 25 to 75 percentile, as well as whiskers of 1.5 IQR. B) Performance of RGN2 for each mutation vicinity and type of backbone change. These graphs show that the results remain unchanged after the adjustment was performed, thus the RGN2 results remaining valid.

Furthermore, all protein structure prediction methods draw on protein sequence homology from databases that include the majority of known protein sequences at the time of their release. Since these methods rely on databases published between 2013 and 2022, it is reasonable to assume that any sequence published prior to a method’s release year may have been part of its training data.

### Protein descriptors

We obtained protein descriptors from multiple sources, including gene ontology (GO) annotations, structural file descriptions, and structural homology classifications. GO annotations were sourced from UniProtKB [69].Structural descriptions were extracted from each protein’s mmCIF files in the PDB.Structural homology classifications were retrieved from the CATH [58] and SCOP [5] databases. These descriptors were aggregated for each protein, as no single source included all the necessary information for our dataset.

In the Results section, we mention the protein descriptors as properties related to a corresponding protein. These properties are obtained by identifying the most commonly found protein descriptors for the protein within our previously mentioned aggregated descriptors. These properties are filtered to eliminate recurrent descriptors found in all proteins to prevent non-specific descriptors for each protein. The process is done through word clouds which are shown in Section 6 of Supplementary Information.

### Secondary structure metrics

To assess the performance of secondary structure prediction methods on Q8, three commonly used metrics are employed: Accuracy (*Q*^*Acc*^), Segment Overlap (*SOV*), and SOV_REFINE. As *SOV* has been improved since its inception, the most common version, SOV99 [78], is the version that we refer to as Segment Overlap. Its most recent modification is referred to as the refined version SOV_REFINE [43]. These metrics are computed using our “Secondary Structure Metrics Calculator” software^3^, which implements previously published secondary structure metrics, such as Accuracy, SOV_REFINE, SOV99, but also includes older *SOV* versions like SOV94 [54].

The metrics require two secondary structures of length *n* as input:

Reference structure: **R**^*ref*^ = [*r*_1_, *r*_2_, …, *r*_*n*_]

Predicted structure: 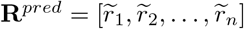

For each SSE class *r*, a *reference segment* (or *reference r-block*) in **R**^*ref*^ is defined as a contiguous substructure 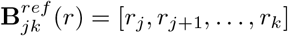 of **R**^*ref*^ that satisfies the following conditions:

Uniform Structure: All residues in the substructure are of SSE class *r* (i.e., *r*_*j*_ = *r*_*j*+1_ = … = *r*_*k*_ = *r*). Boundary Conditions:

-*r*_*j*−1_ ≠ *r* and *r*_*k*+1_ ≠ *r*, thus the SSEs immediately before *r*_*j*_ and after *r*_*k*_ must not be SSE class *r*.

Similarly, for each SSE class *r*, a *prediction r-block* in **R**^*pred*^ is defined as 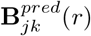 following the same criteria. The sets of all such blocks are defined as:

Reference Blocks:

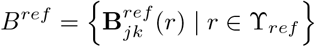

Prediction Blocks:

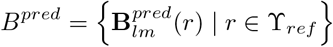

Here, ϒ_*ref*_ represents the set of all secondary structure element (SSE) classes present in the reference structure sequence **R**_*ref*_.

In this study, we evaluate Q8 prediction by measuring the following metrics for each DSSP-assigned secondary structure class within ϒ_8_ ={ℬ, 𝒞, ℰ, 𝒢, ℋ, ℐ, 𝒮, 𝒯}, where ϒ_*ref*_ ⊆ ϒ_8_. Further details on DSSP assignments are given in Section 1 and 2 of Supplementary Information.

All secondary structure metrics evaluate SSEs at each position of both reference and predicted structure sequences using the *identity* function

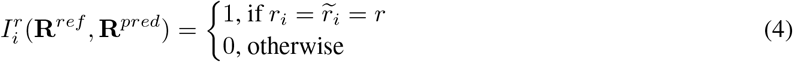

where *i* is the position of the SSE in both secondary structures **R**^*ref*^ and **R**^*pred*^, and *r* is the SSE class.

The Accuracy metric is defined as the ratio of matching SSE pairs between the reference and predicted structures to the total number of SSE pairs. Since the reference and predicted structures have equal lengths, the total number of SSE pairs is equal to the length of the structure sequences, i.e., |*R*^ref^ |= |*R*^pred^| = *n*. This is more formally defined as:

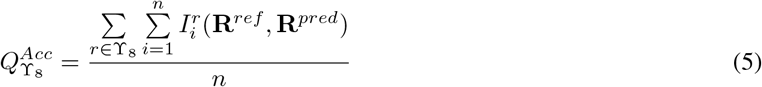

Accuracy, which measures exact matches between two secondary structures at each position, may not be able to sufficiently capture the structural details of slightly misaligned secondary structure elements (SSEs) that extend across the structure sequence. To address this limitation, a more informative metric called Segment Overlap (*SOV*) was introduced [54] and consequently improved [78].

The Segment Overlap metric is a weighted sum over overlapping pairs of segment blocks for each SSE class *r* ∈ ϒ_*ref*_, accounting for slight misalignments in SSEs. Formally, for *r*-blocks 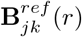 and 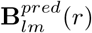, let 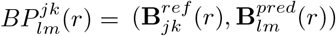 be an **overlapping segment block pair** for **R**^*ref*^ when *j* ≤ *m* and *l* ≤ *k*. Then, let 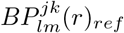 denote the first element of 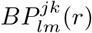 (i.e.,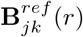) for that overlapping segment block pair.

We denote 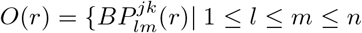 and 1 ≤ *j* ≤ *k* ≤ *n*} as the **set of all overlapping pairs** of *r*-blocks between **R**^*ref*^ and **R**^*pred*^ (i.e., the set of all tuples 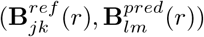 for all *r* ∈ ϒ_*ref*_). If a reference *r*-block 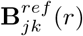 has no overlap with any predicted *r*-block in *B*^*pred*^, we define 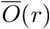 as the set of non-overlapping segment blocks in **R**^*ref*^ for SSE class *r*:

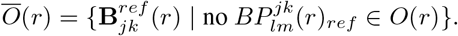

We now specify the functions that take part in the *SOV* definition.

`Norm`(*r*) is the **normalization value** for SSE class *r* defined as,

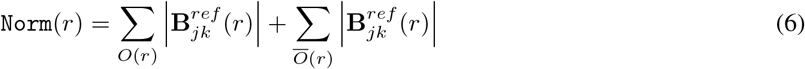

where 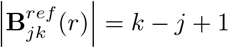 is the length of the *r*-block. It is important to note that any particular reference *r*-block can appear multiple times across different block pairs in *O*(*r*).

LenOv_*r*_ is the number of identical SSEs of class *r* for a pair of segments:

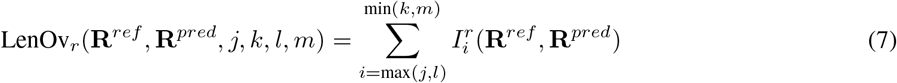

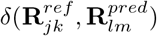 is the amount of **allowable misalignment** given to a pair of segments and defined as,

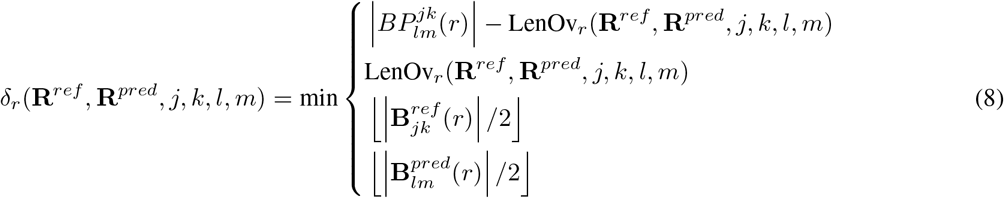

where 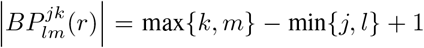 is the combined overlap length of an overlapping pair of segment blocks. An example of the way overlapping segments can obtain the different *δ*_*r*_ values is shown in Fig. 9.

**Figure 9.**
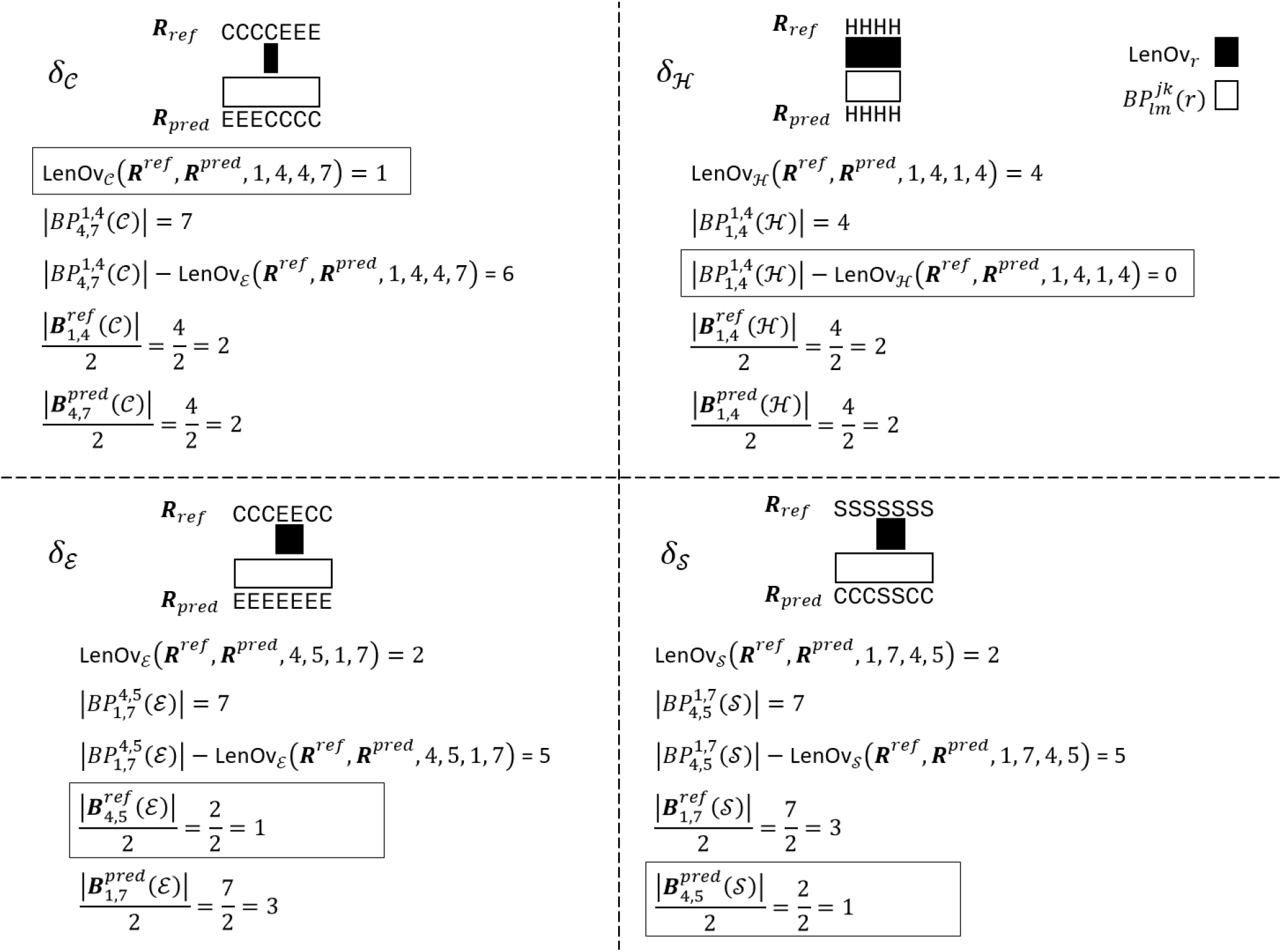
Calculating *δ*_*r*_. Example shows all four possible minimum values depending on the overlapping segments. The resulting value is the misalignment allowed for the *r*-block pair

To define **Segment Overlap**, it is important to note that the set of all overlapping pairs of *r*-blocks *O*(*r*) contain *r*-blocks with their respective starting and ending indices (e.g., 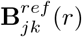, where *j* is the starting index of the *r*-block for **R**^*ref*^, and *k* is the ending index). Then for SSE class *r*, we define Segment Overlap *SOV* (*r*):

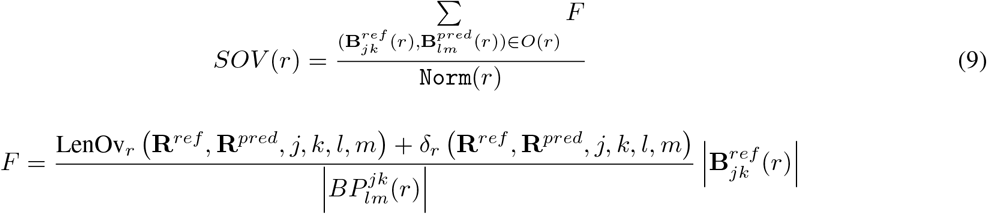

Lastly the overall *SOV* score for all SSE classes is defined as,

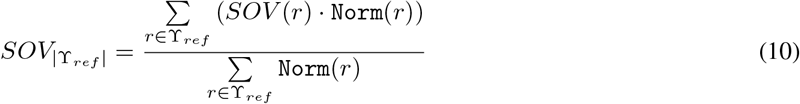

An example for calculating Segment Overlap can be seen in Fig. 10.

**Figure 10.**
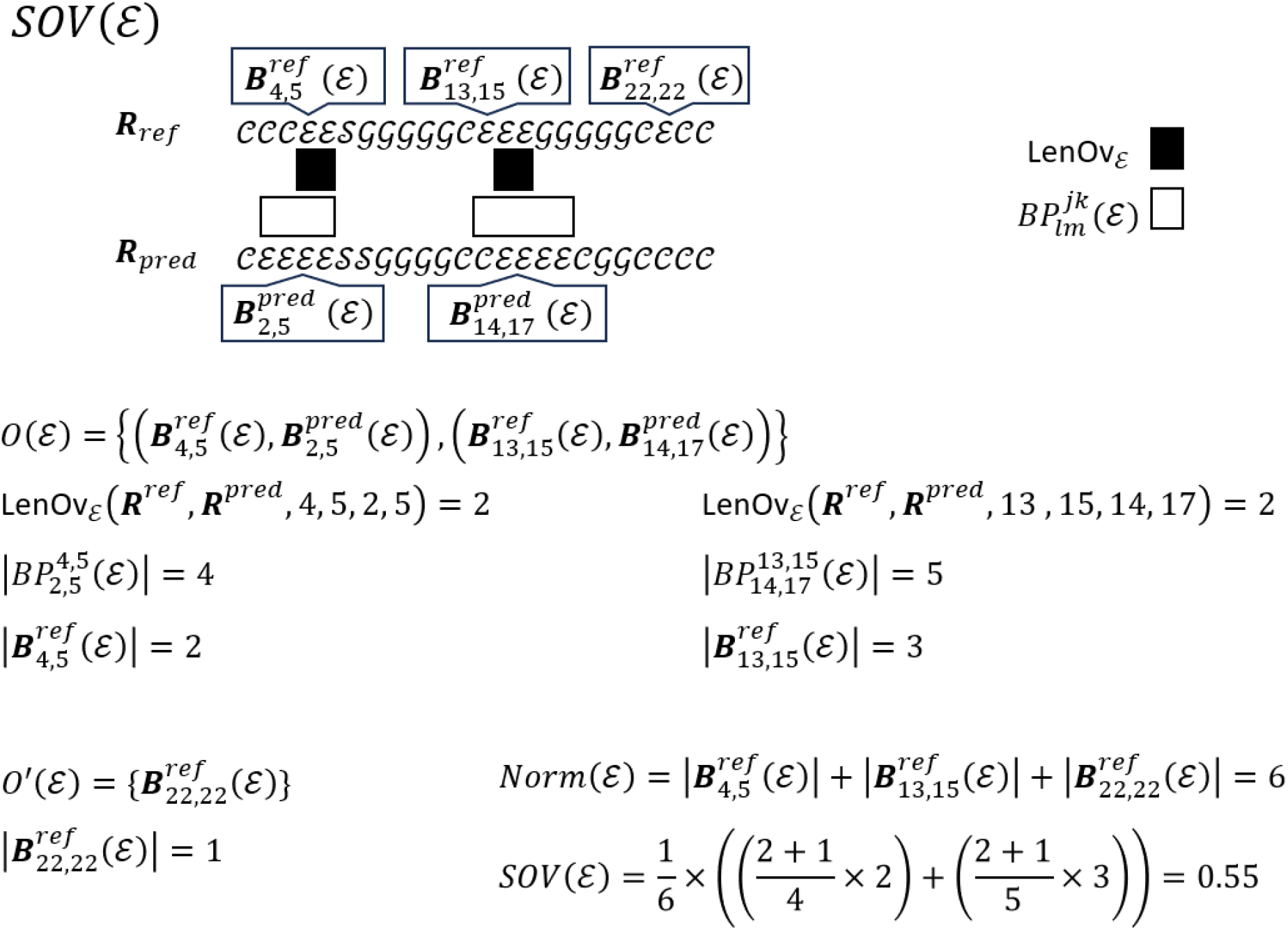
Calculating. *SOV* (ℰ). Example following the nomenclature from our definitions in the Secondary structure metrics section.

The refined version of *SOV*, termed as SOV_REFINE, changes the allowance function as follows,

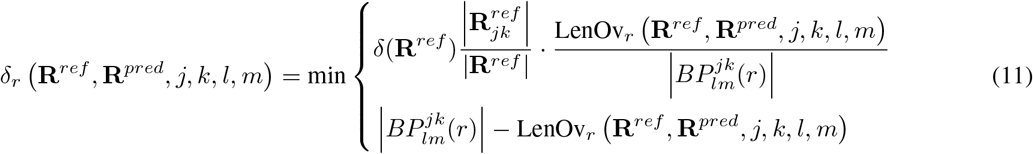

where *δ*(**R**^*ref*^) is the **total misalignment allowance** given for all segment blocks of the reference structure sequence as

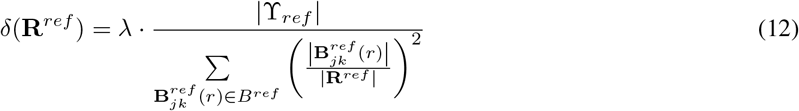

where ϒ_*ref*_ ⊆ ϒ_8_ and |ϒ_*ref*_ | is the number of SSE classes that appear in the reference structure sequence, and *λ* ∈ ℝ, 0 ≤ *λ* ≤ 1 is an adjustable scale parameter that is used to limit the range of *δ*(**R**^*ref*^).

### Metrics calculation

Existing tools for calculating the previously defined metrics are script-based and lack efficiency, particularly for large datasets or longer structure sequences. To address this, we developed a more efficient metric calculation tool that outperforms currently available options. Secondary structure metrics for this project were calculated using our custom software, **SSMetrics** ^4^. The results were validated against the Perl script SOV_refine.pl developed by Liu and Wang [43]. The performance comparison between our SSMetrics and existing methods is shown in Fig. 11.

**Figure 11.**
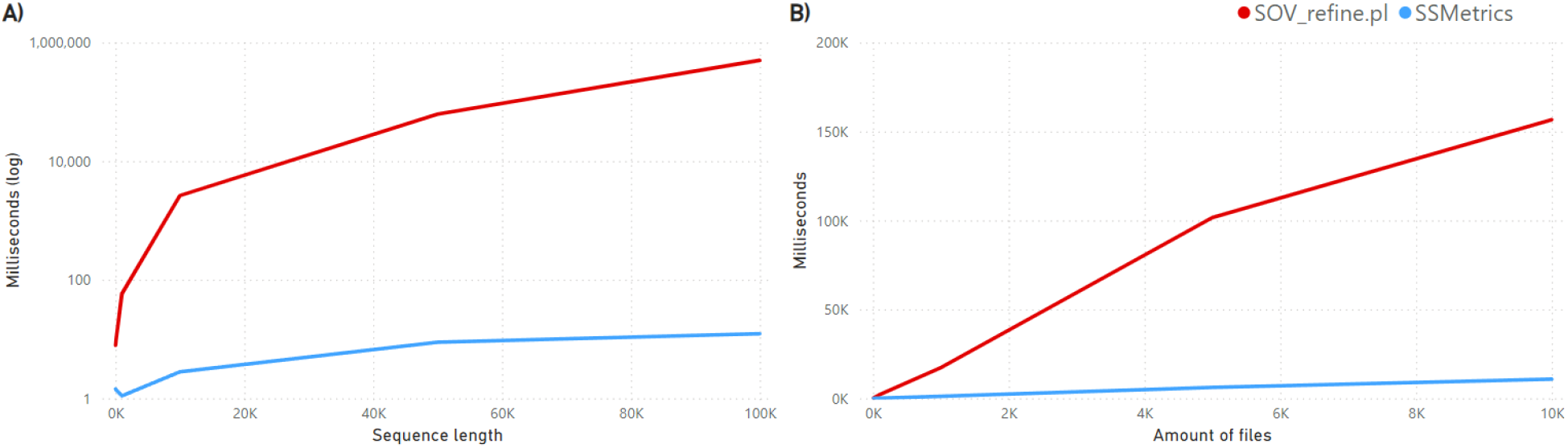
Performance comparison between SSMetrics and SOV_refine.pl. A) Performance measured with the “perf stat” profiling tool across varying structure sequence lengths. B) Performance measured with the “time” tool for runs involving 10, 100, 1,000, 5,000, and 10,000 files of 500 amino acids each.

### Mutational metrics

Mutational data clusters consist of a wild-type protein sequence along with its associated mutations. The previously discussed metrics are specifically designed to compare two secondary structures: a reference structure and a predicted structure. For each protein, the reference structure is derived using DSSP, while the predicted structure is generated by a structure prediction method.

To evaluate the performance of prediction methods on mutational data, it is essential to account for mutational changes. To achieve this, we introduce three mutational metrics, described below.

**Mutational consistency** measures the Accuracy between mutational changes in the reference and predicted structures. To compute this, we:

1. compare the wild-type and mutated reference structures element-wise for each secondary structure element (SSE). This yields the **reference mutational change** sequence, indicating whether each SSE is preserved (“N’) or changed (“C’) after mutation.
2. perform the same comparison for the wild-type and mutated predicted structures, producing the **predicted mutational change** sequence.
3. calculate Accuracy for the reference and predicted mutational change sequences, as described in the Secondary structure metrics section, treating the two possible states—’preserved’ and “changed’—as binary classes.

By framing the problem as a binary classification task, additional binary classification statistics [12] can be used to further evaluate mutational consistency.

**Mutational accuracy** measures Accuracy for SSE mutations in the reference and predicted structures. The calculation involves the following steps:

1. Compare the wild-type and mutated reference structures element-wise for each secondary structure element (SSE). This generates the **reference SSE mutation** sequence, indicating the type of change, e.g., “EE’ for no change in a *β*-strand or “EI’ for a *β*-strand changing into a *π*-helix.
2. Perform the same comparison for the wild-type and mutated predicted structures to produce the **predicted SSE mutation** sequence.
3. Calculate Accuracy for the reference and predicted SSE mutation sequences, as outlined in the Secondary structure metrics section, with the SSE mutation change types replacing the standard secondary structure classes.

For Q8, there are 64 possible SSE mutation classes, as each of the eight secondary structure classes can either remain the same or change into any other class after mutation.

**Mutational precision** is calculated as Accuracy, Segment Overlap, or SOV_REFINE between interlaced SSE sequences of the wild-type and predicted secondary structures. The process involves:

1. Interlacing the equal-length wild-type and mutated structures element-wise for each SSE. For a wild-type secondary structure **R**^*rep*^ = [*r*_1_, *r*_2_, …, *r*_*n*_] and a mutated structure 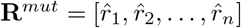, the **interlaced SSE sequence** is defined as:

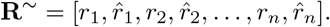
2. This procedure is applied to both the reference and predicted structures, yielding the **reference interlaced SSE** and **predicted interlaced SSE**.
3. Finally, Accuracy, Segment Overlap, or SOV_REFINE is computed between the reference and predicted interlaced SSE sequences, as described in the Secondary structure metrics section.

### Tertiary structure assessment

Tertiary structure prediction methods typically output results in “PDB’ format, though some, like AlphaFold2, also support mmCIF format. For consistency, we used the “PDB’ output for all methods. To align with the process used for experimental structures, we converted the predictions to mmCIF format using MAXIT^5^. We then applied DSSP to assign secondary structures to the mmCIF-formatted predictions. We normalized the output as outlined in the Secondary structure metrics section to produce secondary structure predictions from tertiary structure predictions.

Similar to secondary structure prediction methods, tertiary structure predictions were evaluated based on local, distant, and global structural changes. Mutation types were determined uniformly across all methods, using the protein sequence most similar to the consensus (wild-type sequence) within each cluster.

As Alphafold2 took a considerably longer amount of time than other tertiary structure prediction methods, we utilized batch processing to increase its throughput. The details on batch processing are detailed in Section 10 of Supplementary Information.

RGN2 was originally developed as a Colab notebook, which is designed to be used online. We transformed the online scripts to be utilized locally for our experiments and for better ease of access to others. Details on the changes can be seen in Section 11 of Supplementary Information.

## Results

For each structure prediction method, we evaluated their performance using the Accuracy, Segment Overlap, and SOV_REFINE metrics. These metrics were calculated for different structural and mutation vicinities. The methods evaluated include: AlphaFold2 (“af2’), ColabFold (“colabfold’), ESMFold (“esmfold’), RGN2 (“rgn2’), SSPro8 (“sspro8’), Raptor-X Property (“raptorx’), SPOT-1D (“spot_1d’), SPOT-1D-Single (“spot_1d_single’), and SPOT-1D-LM (“spot_1d_lm).

Metrics were analyzed for the following backbone vicinities: primary structure (“1d’), secondary structure (“2d’), tertiary structure (“3d), and contact/distance vicinity (“contact’). Additionally, results were grouped by mutation vicinity types: local, distant, and global.

Performance results for top-, average-, and low-performing methods are provided as tables in Section 5 of Supplementary Information. These categories are based on performance trends, as shown in Fig. 12. In the figure, the error bars represent the 1st to 99th percentiles. This reveals the performance spread of the methods, indicating that even top-performing methods struggle with some proteins. Further details of the performance spread can be seen as a box plot in Section 7 of Supplementary Information.

**Figure 12.**
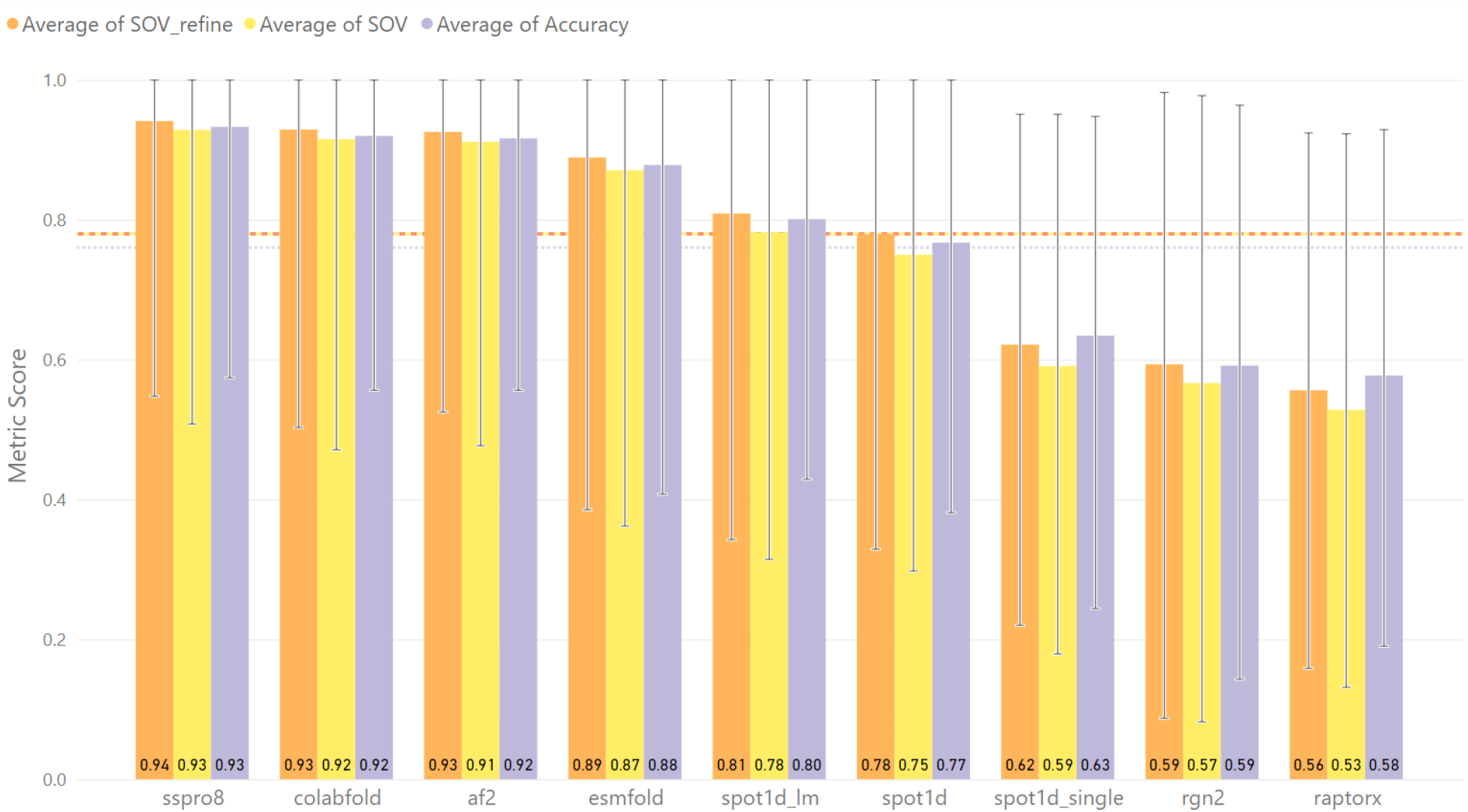
Performance of each structure prediction method. SSPro8, ColabFold, Alphafold2, and ESMFold perform higher than average. SPOT-1D-LM and SPOT-1D perform close to average. SPOT-1D-Single, RGN2, and Raptor-X Property perform below average. Therefore, methods are categorized according to their performance as “Top’, “Avg’, or “Low’ respectively.

### Mutational metrics

Since mutational consistency reduces structural changes to a two-class problem, we were expecting high scores across all prediction methods. Surprisingly, low-performing methods achieve similar scores to average- and top-performing methods. The exception is RGN2, which performs lower overall but still has higher minimum scores compared to other methods.

When binary classification statistics are applied, treating structural change (“C’) as the positive class and structural preservation (“N’) as the negative class, mutational consistency scores reveal more variation. However, since structural changes are rare, the classification problem is highly imbalanced.

Fig. 13 reveals that all methods exhibit high False Discovery Rates and False Negative Rates, along with low Positive Predictive Values and Sensitivity. This indicates that, regardless of their overall mutational consistency scores, the methods struggle to accurately predict mutational changes, often failing to detect structural changes when they occur. The high mutational consistency scores are largely due to the rarity of structural changes, resulting in a strong bias toward preserved structures.

**Figure 13.**
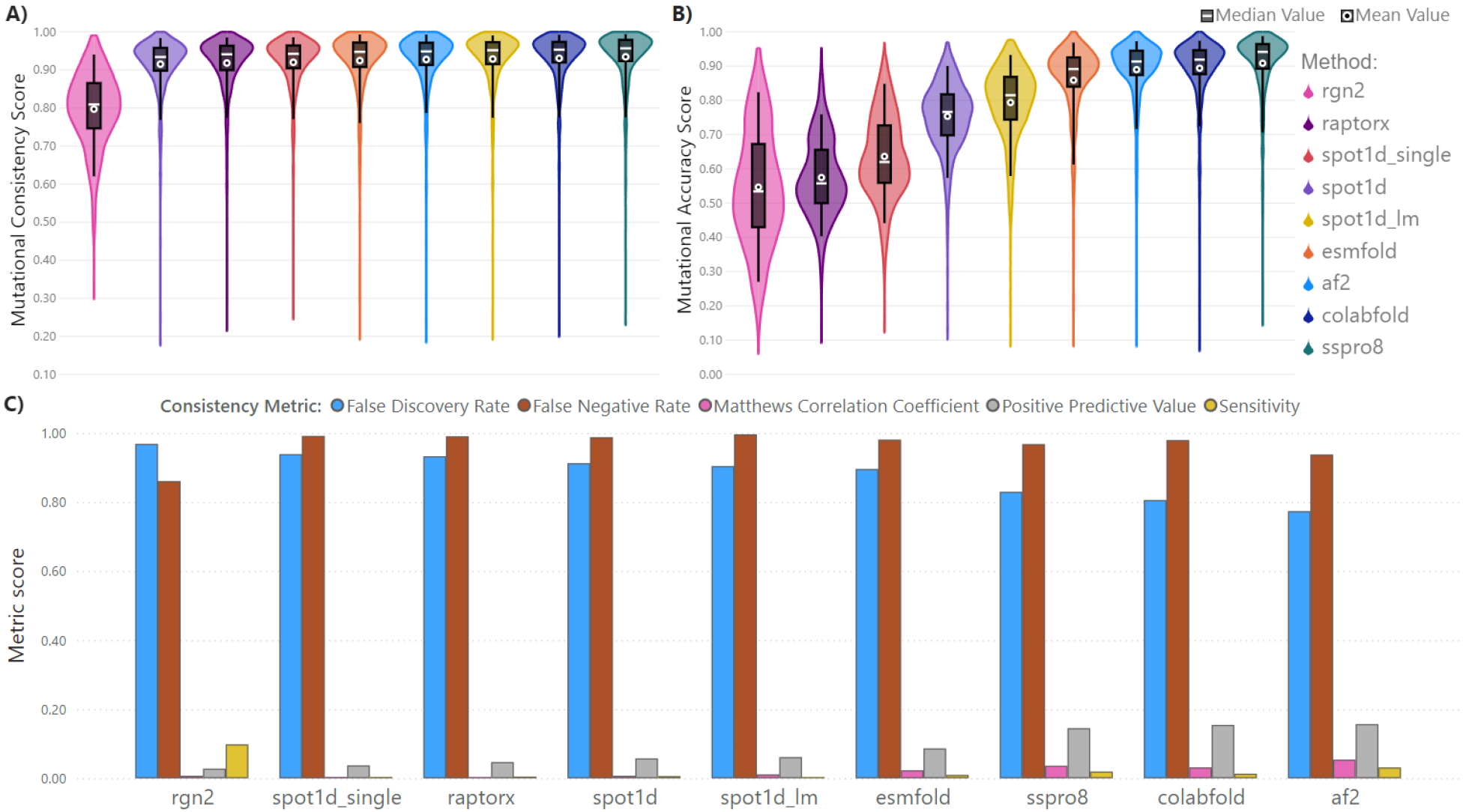
Mutational consistency and Mutational accuracy. Violin plots display the mutational consistency (A) and mutational accuracy (B) results for each structure prediction method. C) A bar graph presents the binary classification metrics for mutational consistency across all prediction methods. This shows that all methods have deficiencies predicting if and when a mutational change will occur. The high scores in A and B come from the data imbalance of very few mutational secondary structure changes occurring.

Mutational accuracy follows a scoring trend similar to SOV_REFINE benchmark results, which calculate secondary structure metrics on a protein-by-protein basis without focusing on mutations. However, mutational accuracy reduces scores across all methods by approximately 4%. For example, SSPro8 achieves a mean mutational accuracy of 91%, compared to 95% in the protein-by-protein SOV_REFINE benchmark, providing a more realistic assessment of predictive performance. The mutational consistency and accuracy results are shown in Fig. 13.

Mutational precision using Accuracy follows a similar trend to SOV_REFINE in Fig. 12, with Raptor-X Property scoring lowest, followed by RGN2, SPOT-1D-Single, SPOT-1D, and SPOT-1D-LM. When using Segment Overlap or SOV_REFINE, the order changes, with RGN2 scoring lowest, followed by Raptor-X Property. Additionally, most metrics show lower mean values when calculated with a Segment Overlap metric. Notably, mutational accuracy strongly correlates with mutational precision when using a Segment Overlap metric. The mutational precision results are shown in Fig. 14.

**Figure 14.**
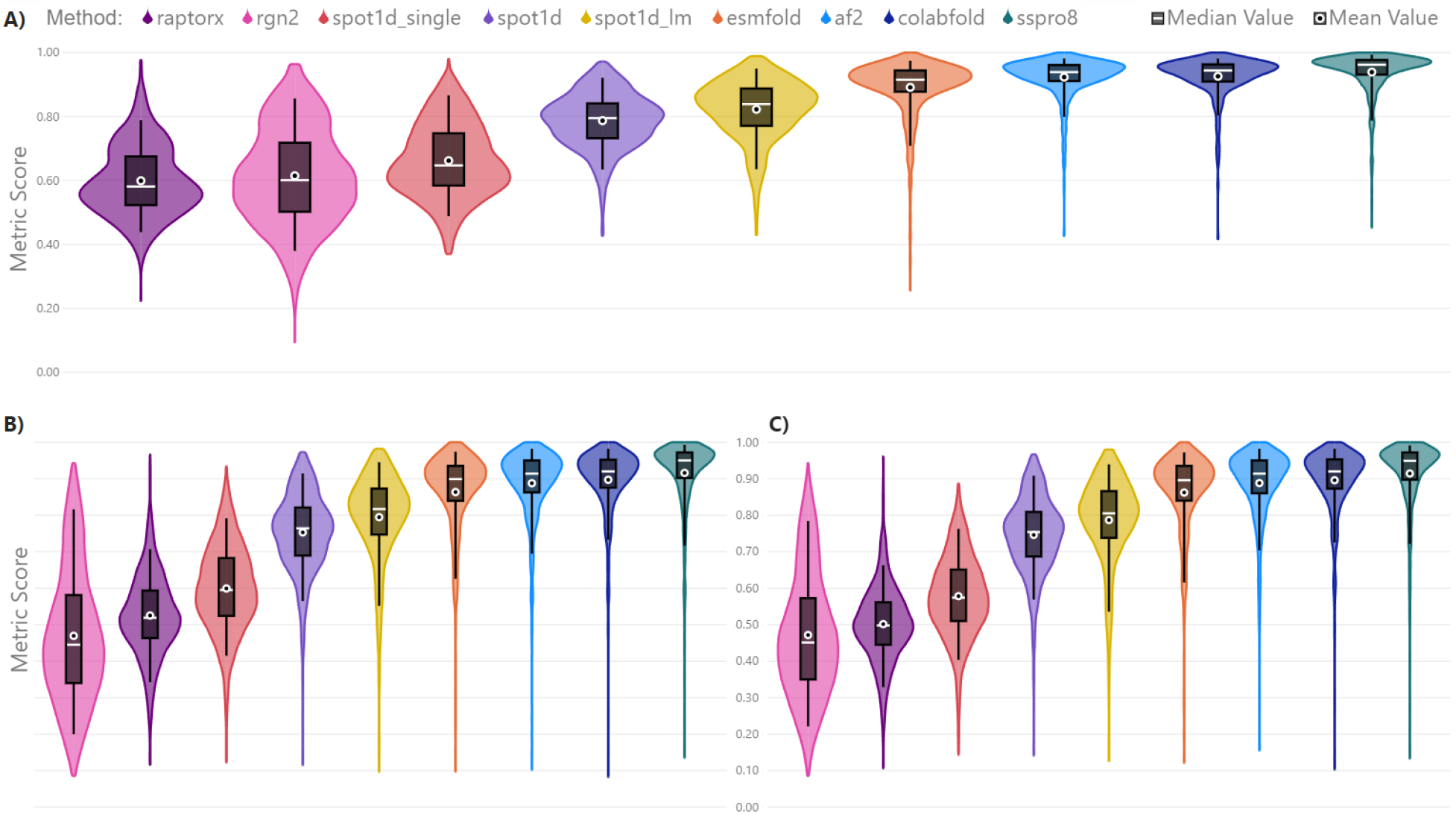
Mutational precision. Violin plots showing mutational precision for each prediction method using three different secondary structure metrics: A) Accuracy, B) Segment Overlap, and C) SOV_REFINE. Results for mutational precision metrics are very similar to their respective individual protein secondary structure metrics.

### Mutation stability

Single amino acid mutations can produce stable, without a structural change, or disruptive, inducing a structural change, results. In our mutational dataset, containing experimental structure from the PDB, stable mutations account for 128 out of 915 mutated proteins. Disruptive mutations were considered by their 1D vicinity: local and distant. Local disruptive mutations constitute 429 of the 787 total disruptive mutations. Distant disruptive mutations are found in 675 mutated proteins. In contrast to the experimental data, structure prediction methods tend to infer a disruptive mutation for almost all proteins in our dataset. This can be of interest to structure prediction researchers, as even the best methods are still lacking on predicting stable mutations. In the PDB data, stable mutations are found mostly in transport proteins. This transport property of a protein is also found in mutations that cause a structural change in its distant vicinity, but not in its local vicinity, This is logical since there is evolutionary pressure on keeping important functional structures, e.g. transport and complex-forming capabilities, as they affect downstream processes in the cell. From our selected prediction methods, SSPro8 is the most capable method for stable mutations. It is able to predict stable mutations in transport proteins, but struggles on disruptive mutations that only affect its local vicinity. This type of disruptive mutation deficiency for SSPro8 can be found in complex-forming proteins. This discrepancy may arise because some experimental PDB structures are resolved in their ligand-bound state, potentially biasing the data toward bound conformations, which may differ from unbound structures. Prediction methods, however, do not account for ligands and are designed to predict unbound structures. It is also possible for the extraordinary results from SSPro8 to be caused by its use of templates that contain most of our mutational dataset. The results for mutation stability data can be seen in Fig 15.

**Figure 15.**
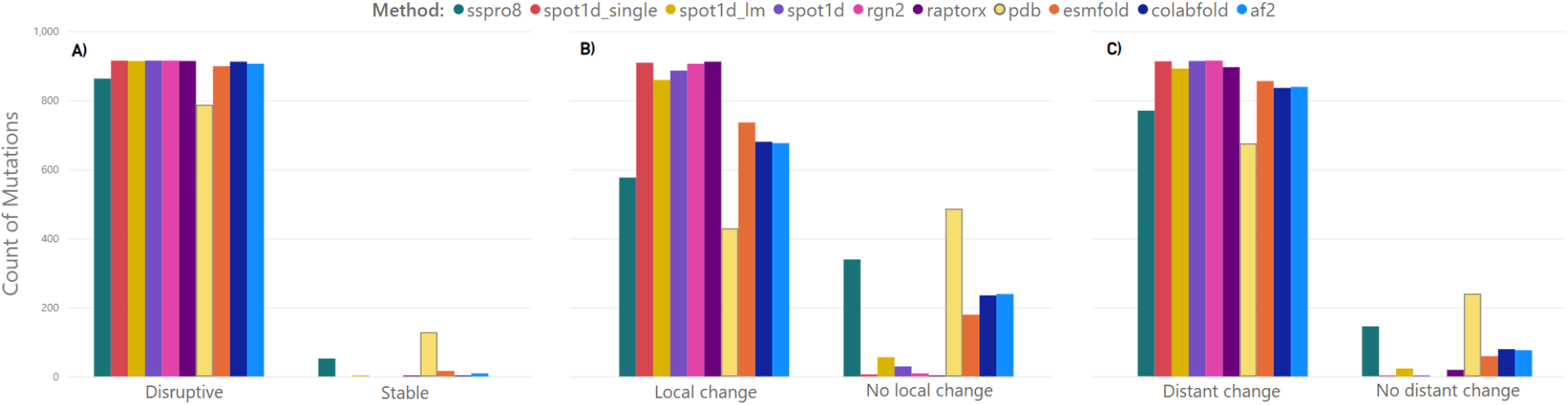
Mutation stability results. Amount of mutations, for all prediction methods and experimental PDB data, with a disruptive or stable result in the A) complete secondary structure, B) Local vicinity, C) Distant vicinity of a mutation. Stable mutations occur more often in PDB data than in prediction methods, as the latter almost always predicts destabilizing mutations. The exception is SSPro8 while still missing two thirds of stabilizing mutations. PDB data also show that the local vicinity is more stable than not when a mutation occurs.

### Mutation vicinity

Examining the vicinity results for each type of backbone change reveals that methods struggle more with accurately predicting local changes. The shorter maximum length of local changes compared to distant and global changes likely contributes to greater variability in performance. The lower average metrics for local changes result from all methods performing slightly worse on these changes, as reflected in Fig. 16.

**Figure 16.**
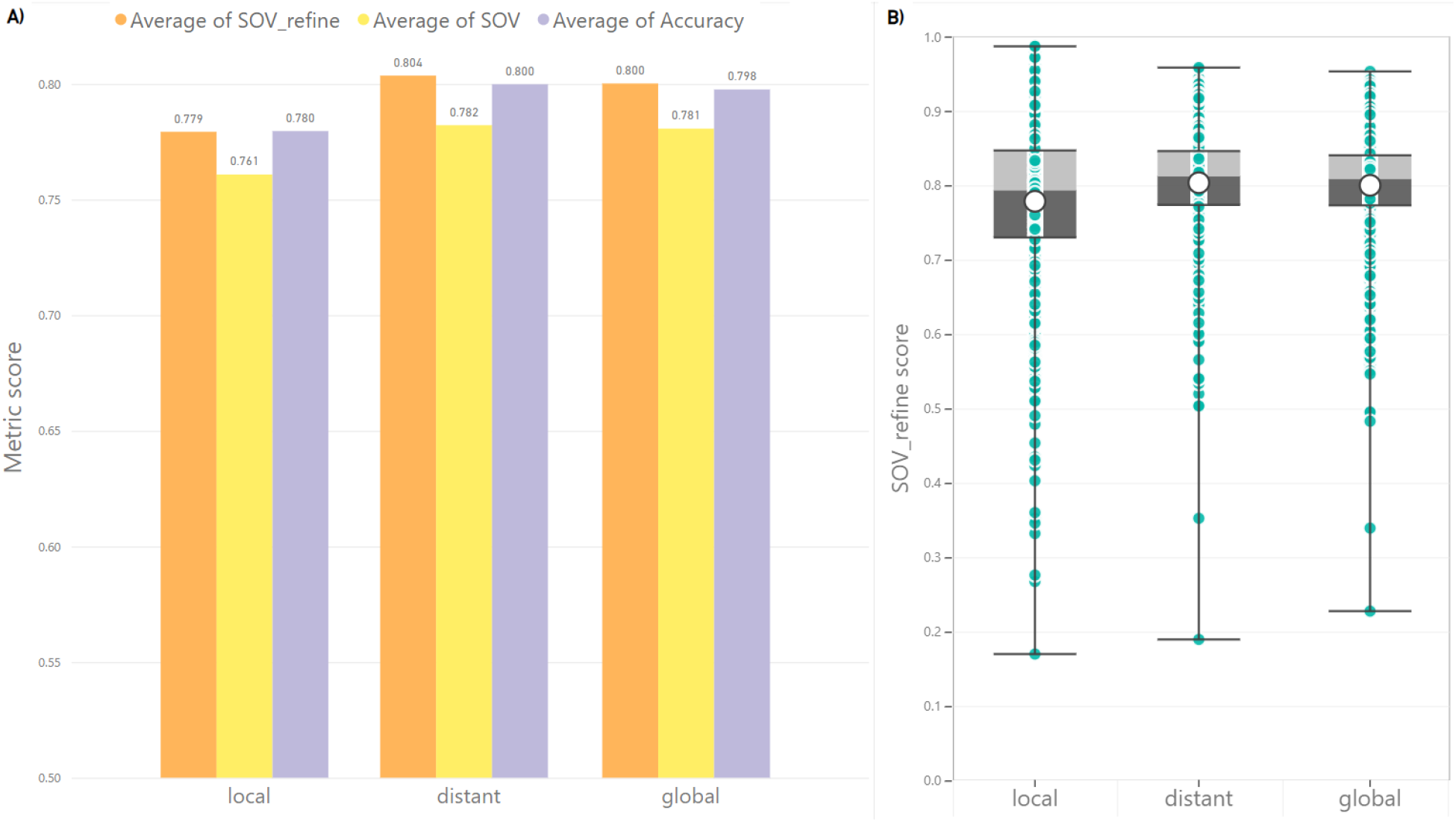
Statistics for Type of Backbone changes. Accuracy, Segment Overlap and SOV_REFINE metrics for each type of backbone change. A) Bar graph showing the average performance across the three metrics, highlighting lower accuracy in predicting local changes. B) Box plot of SOV_REFINE values for each type of backbone change, illustrating a wider spread in local change predictions, ranging from the highest to the lowest overall results.

Focusing on disruptive mutations, PDB data reveals a few common single amino acid mutations that cause secondary structure changes, such as Asparagine (polar) to Histidine (positively charged), Serine (polar) to Threonine (polar), and Alanine (non-polar) to Leucine (non-polar). In contrast, many mutations rarely caused more than one secondary structure change, as shown in Fig. 17.

**Figure 17.**
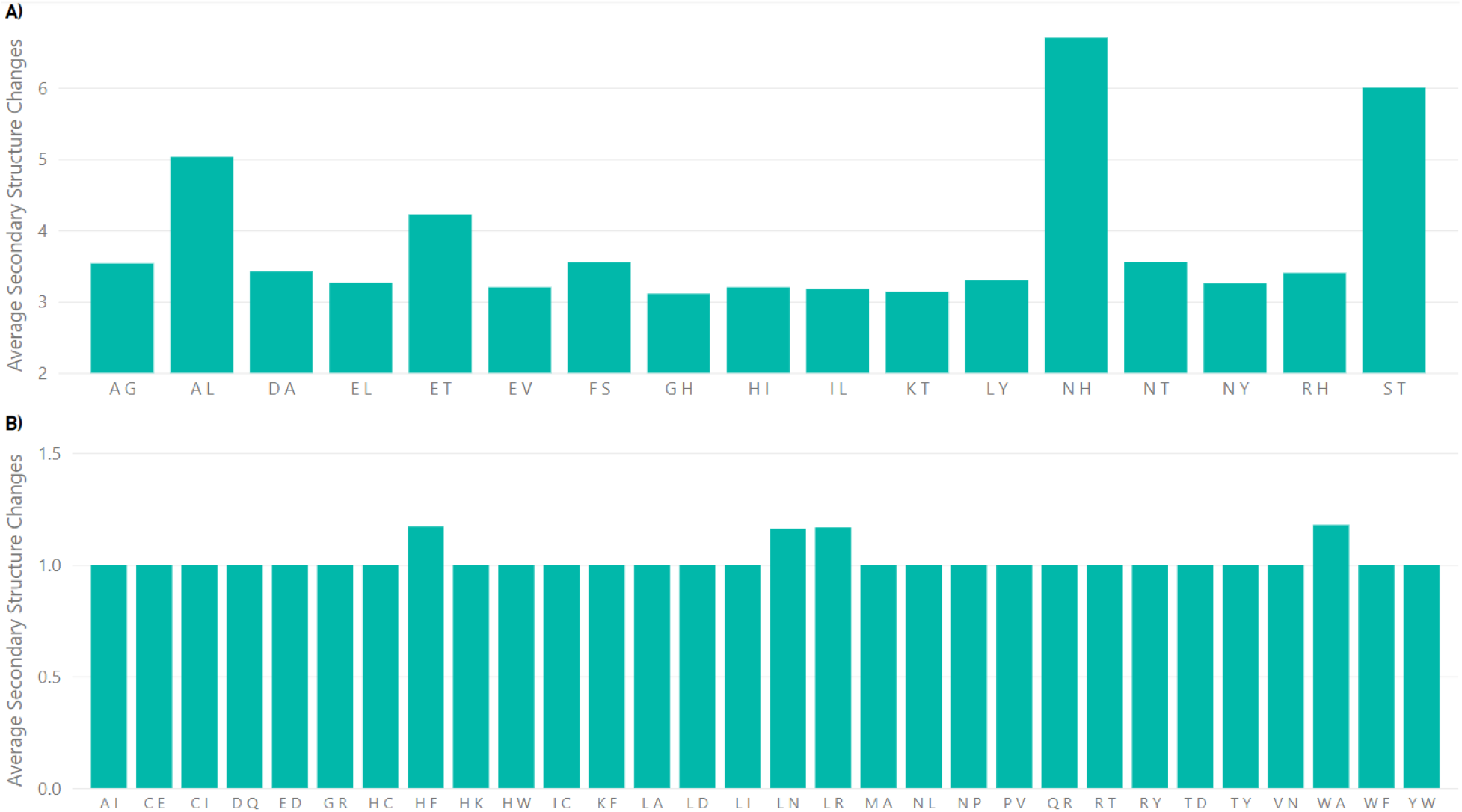
Amino acid mutations results. This data shows mutations in two letter codes. The first letter is the wild-type amino acid and the second letter is the mutated amino acid. A) Most common disruptive mutations in our dataset. B) Least common disruptive mutations that appear in our dataset.

As previously seen in mutation stability results, secondary structural changes due to single amino acid mutations occur less frequently in experimental PDB data than in predicted structures. The SSE classes that produce the previously mentioned discrepancies can be seen in detail in Fig. 18. In our dataset, *π*-helices commonly transition into *α*-helices, loosening the helix structure. Less frequently, *π*-helices dissolve into hydrogen-bonded turns or bends. They never transition into *β*-sheets, bridges, 3-10 helices, or coils, as such changes would require significant energy, likely exceeding what a single mutation can induce. This thermodynamic constraint appears to be absent in structure prediction methods, limiting their ability to produce more realistic predictions.

**Figure 18.**
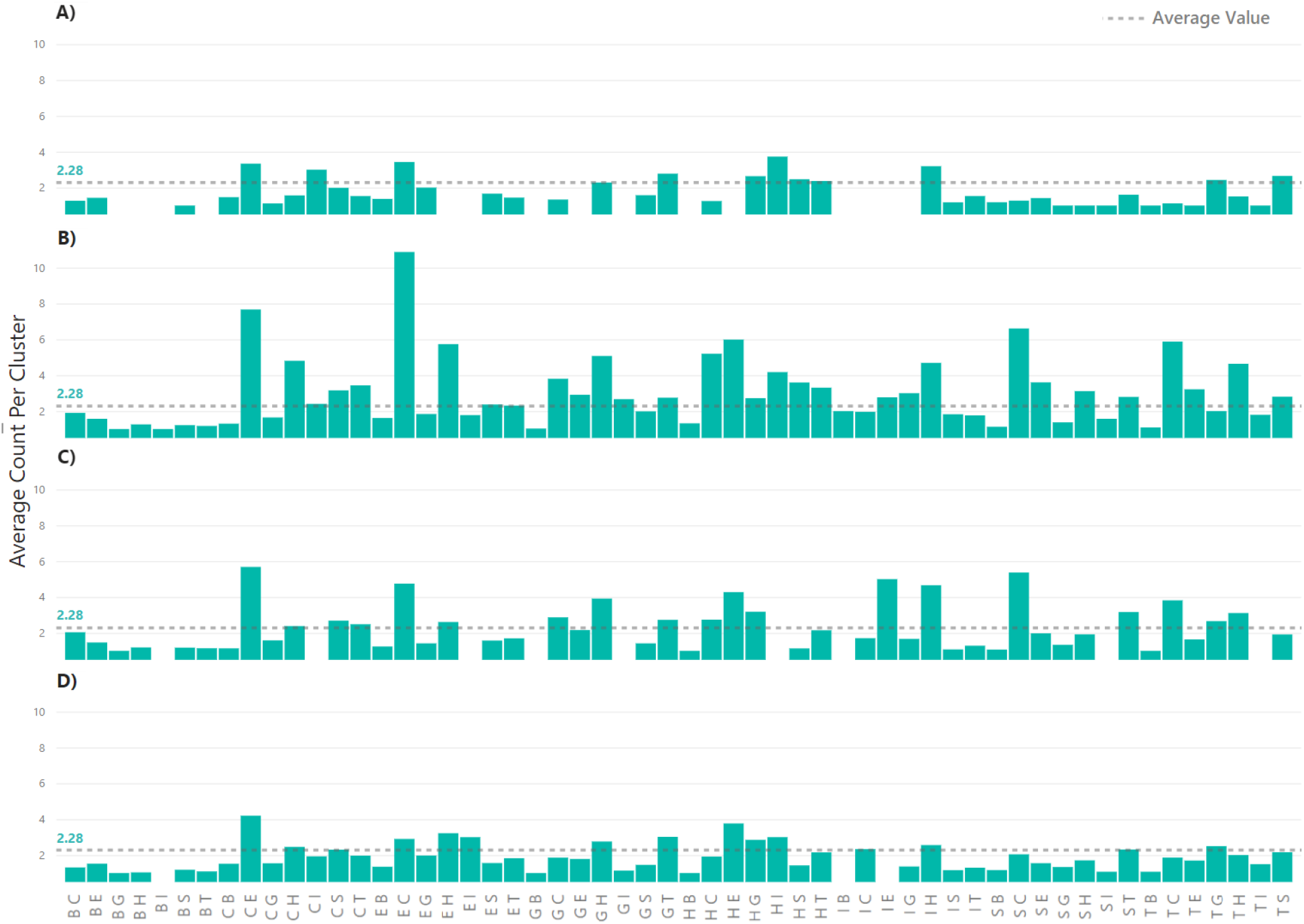
Secondary structure mutations results. All bars show the average amount of times a secondary structure mutation occurred in a cluster. Secondary structure mutations follow a two letter code. First letter is the secondary structure assigned to the wild-type amino acid, while the second letter is the secondary structure assigned to the mutated amino acid. A) PDB data from experimentally obtained structures. B) “Low’ performing structure prediction methods. C) “Average’ performing structure prediction methods. D) “Top’ performing structure prediction methods.

### Prediction difficulty for methods

Most structure prediction methods struggle to predict stable mutations (i.e. single amino acid mutations that do not cause a change in the structure), even though such mutations occur in reality. Transport proteins are a notable exception, as these proteins are inherently stable, and minor prediction errors do not significantly affect their structure. This stability allows transport proteins to often be correctly predicted, aligning with their actual behavior. However, transport proteins also appear among incorrectly predicted structures, highlighting inconsistencies in prediction accuracy for these proteins.

Incorrectly predicted proteins frequently include membrane and structural proteins, which often form complexes and contain *β*-sandwich structures characterized by anti-parallel *β*-sheets. These features are also common in proteins with stable mutations. Therefore, the beta structures found in these types of proteins could be the factor leading to inaccurate predictions for stable mutations.

These findings are illustrated in Fig. 19.

**Figure 19.**
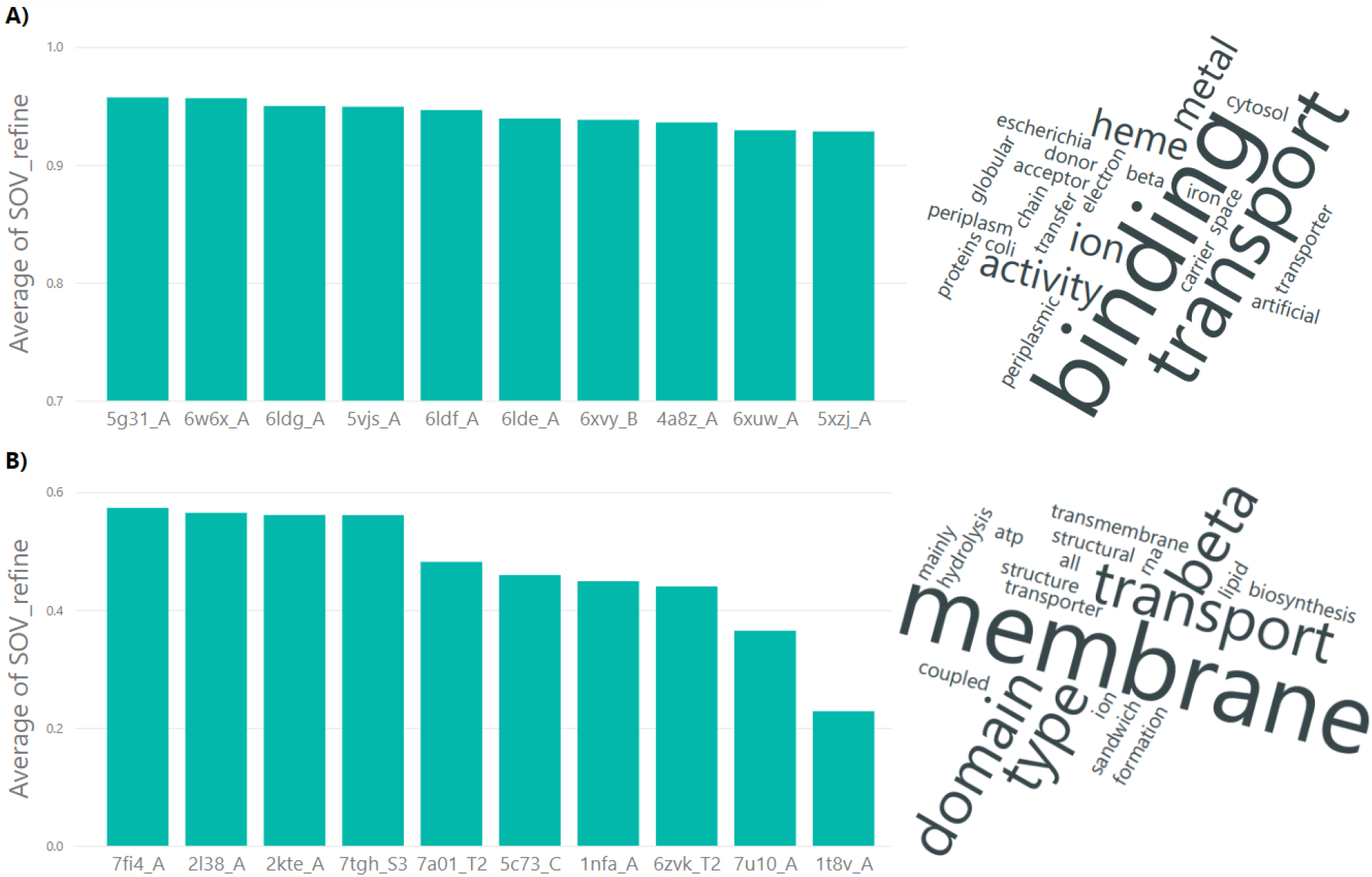
Overall protein structure prediction results. Structural properties are an agglomeration of protein descriptors from CATH, SCOP, and PDB. The proteins are named in the following manner: PDB ID (underscore) Protein chain. A) Best overall predicted proteins and their structural properties. B) Worst overall predicted proteins and their structural properties.

### Method comparisons

In this section, we analyze the similarities and differences between methods in each performance category by examining their best- and worst-predicted proteins using three secondary structure prediction metrics.

Among the top-performing methods, the best-predicted protein chains often include immunoglobulin-like *β*-sandwich domains [16], one of the most common structural motifs. These domains are present in a wide variety of proteins, including those in the extracellular matrix, muscle proteins, immune system proteins, cell-surface receptors, and enzymes.

The best-predicted protein chains for average-performing methods often include transport proteins involved in nuclear and cytoplasmic transfer. Some also relate to lipid binding, suggesting an association with the lipocalin family. The lipocalin family, characterized by an antiparallel *β*-barrel structure surrounding its binding site, transports small hydrophobic molecules such as lipids and binds to complexed iron molecules and heme.

The best-predicted protein properties for low-performing methods include heme binding and periplasmic activity, suggesting an association with periplasmic heme-binding proteins. In bacteria, these proteins are part of the heme acquisition system, transferring heme across the periplasmic space from the outer membrane for energy acquisition. These results are shown in Fig. 20.

**Figure 20.**
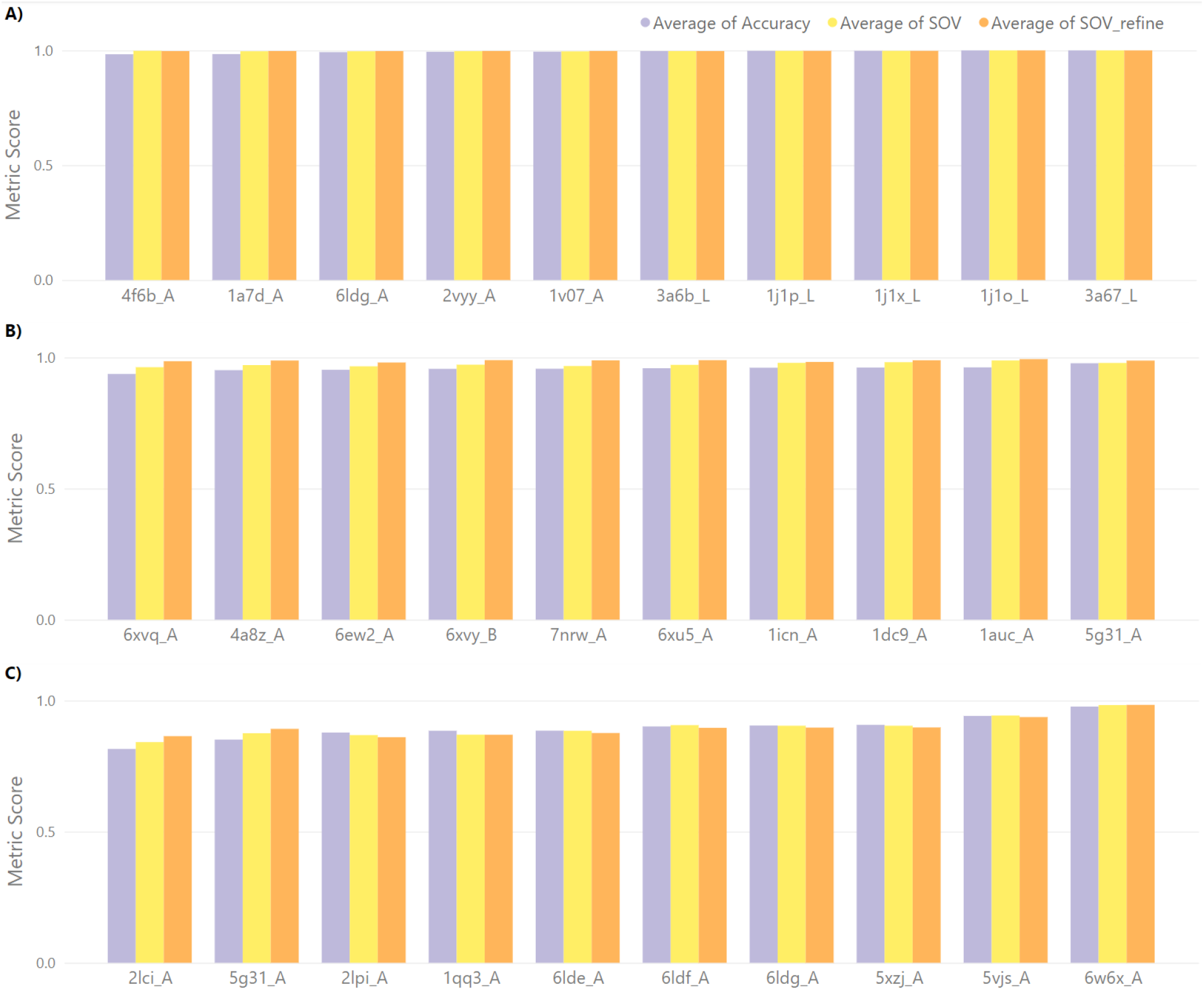
Best results per method category. Best predicted proteins for each method performance category. There are only a few proteins in multiple performance categories. A) Top performing methods. B) Average performing methods. C) Low performing methods.

The worst-predicted proteins for top performing methods are similar to the overall mispredicted proteins. These include membrane and structural proteins with a connection to RNA, suggesting an association with membrane-associated RNA-binding proteins. These proteins play a role in organelle-coupled translation, facilitating efficient protein localization within the cell.

The worst-predicted proteins for average performing methods are primarily associated with viral proteins and some membrane activity, suggesting a link to surface proteins critical for viral infection. The prediction challenges may stem from the diverse binding capabilities of these proteins. Additionally, the receptor-binding site flexibility of surface proteins, which is not captured in the static, crystallized structures within the PDB, could further contribute to the increased difficulty in prediction.

The worst-predicted proteins for low performing methods also include transport and metal ion binding properties, such as heme binding. Interestingly, these properties were also present in the best-predicted proteins, indicating inconsistency in the performance of low-performing methods when predicting certain protein types. As a result, no clear association between prediction performance and specific protein types could be identified. While low-performing methods exhibit some overlapping protein properties, no significant patterns were observed. The worst-predicted proteins for different method categories are shown in Fig. 21.

**Figure 21.**
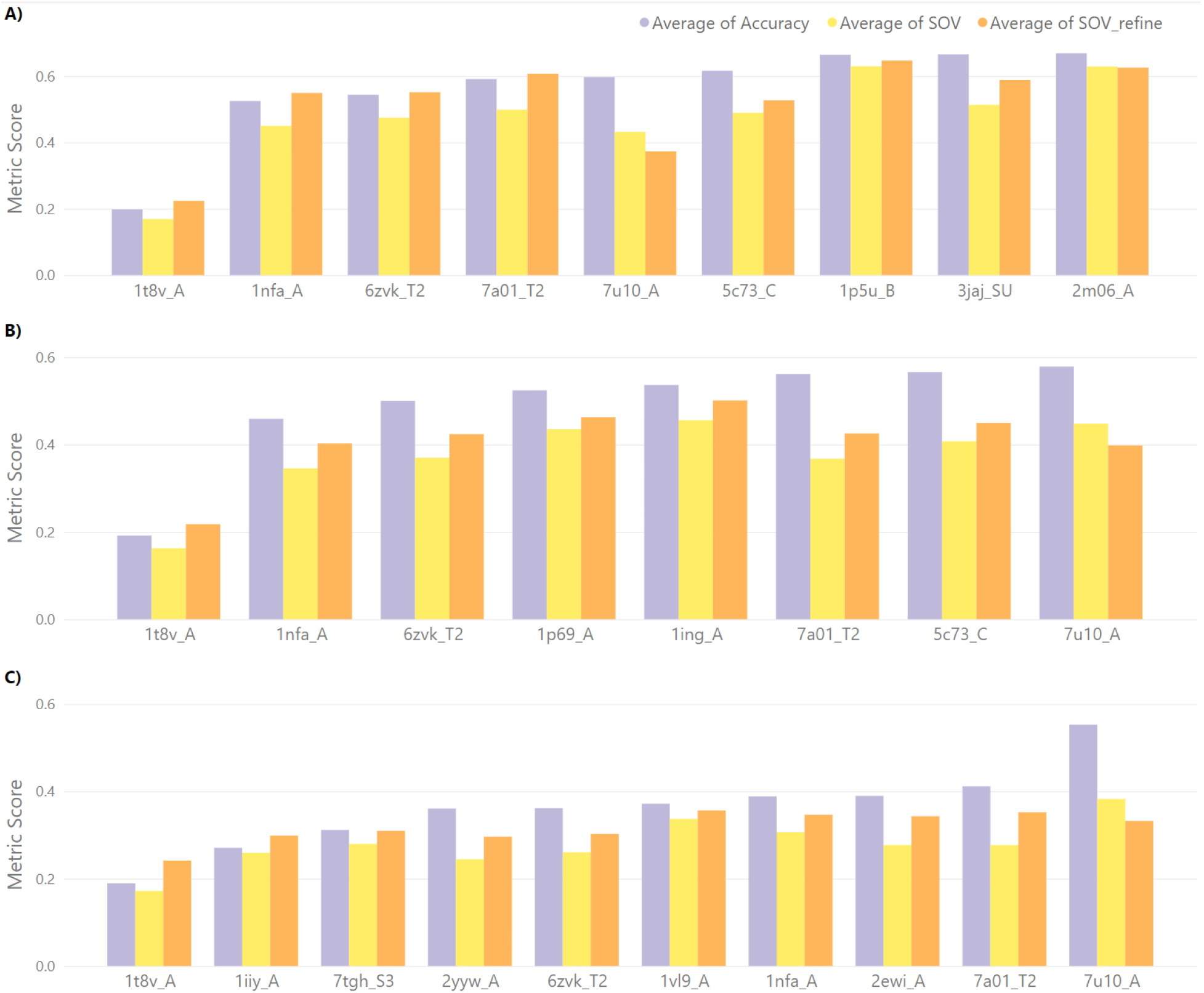
Worst-predicted proteins per method category. The worst-predicted proteins along with their properties for the different method performance categories. Many proteins are equally incorrectly predicted among all performance categories. A) Top performing methods. B) Average performing methods. C) Low performing methods.

### Methods strengths and weaknesses

The structure prediction methods analyzed in this work employ distinct methodologies, resulting in varying performance outcomes. This section highlights the protein properties that posed challenges for each method.

AlphaFold2 struggles with the same protein properties observed in the overall worst-predicted proteins for top performing methods. This is unsurprising, as AlphaFold2 and ColabFold produce similar results due to their closely related algorithms. Their similarity biases the top performing category by contributing a disproportionate number of similar results.

Both AlphaFold2 and ColabFold perform well overall, but their predictions for mutation stability are notably weak, underscoring the need for further research on stable mutation prediction. Between the two, ColabFold is optimized for efficiency, making it the more favorable choice for this study.

ESMFold, like AlphaFold2 and ColabFold, struggles with transport proteins. However, as a language model, ESMFold does not require MSAs, significantly reducing its prediction time by an order of magnitude. While its overall performance is comparable to AlphaFold2 and ColabFold, ESMFold has a slightly lower average accuracy. Nevertheless, its faster prediction speed makes it a strong alternative for handling higher workloads.

SSPro8 stands out as the only top performing method specifically designed for secondary structure prediction. Unsurprisingly, it achieves the best overall performance for our task. Analyzing its worst predictions reveals no clear pattern, suggesting that SSPro8 is a robust solution without bias toward specific protein types. The results for each top performing method are shown in Fig. 22.

**Figure 22.**
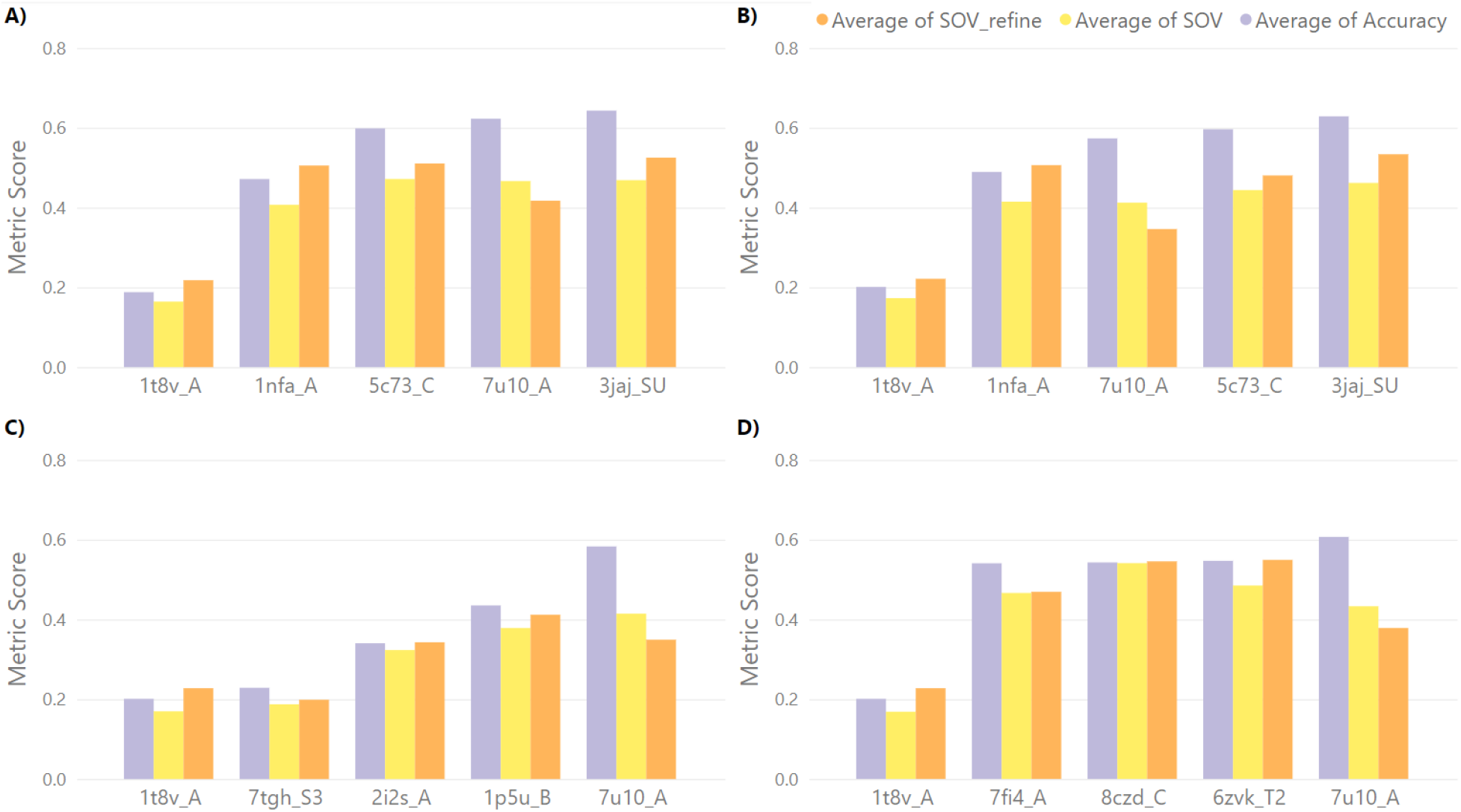
Limitations on Top performing methods. Worst performing proteins for each of the top performing methods. Alphafold2 and Colabfold have very similar performance and thus perform the same prediction mistakes. ESMFold and SSPro8 have very different methodologies to the other two methods and thus perform differently. “1t8v_A’ is commonly predicted incorrectly across all methods. Prediction methods: A) AlphaFold2, B) ColabFold, C) ESMFold, D)SSPro8.

The average performing methods, SPOT-1D and SPOT-1D-LM, struggle with predicting proteins related to viruses and RNA binding. Additionally, SPOT-1D has difficulty with metal ion binding proteins, a challenge also observed in top performing methods. Both are secondary structure prediction methods that underperform compared to their tertiary structure counterparts. As a language model, SPOT-1D-LM achieves higher accuracy than SPOT-1D by not relying on MSA evolutionary information, while SPOT-1D’s reliance on MSAs mirrors the approach of AlphaFold2 and ColabFold. The results for these methods are shown in Fig. 23 A and Fig. 23 B.

**Figure 23.**
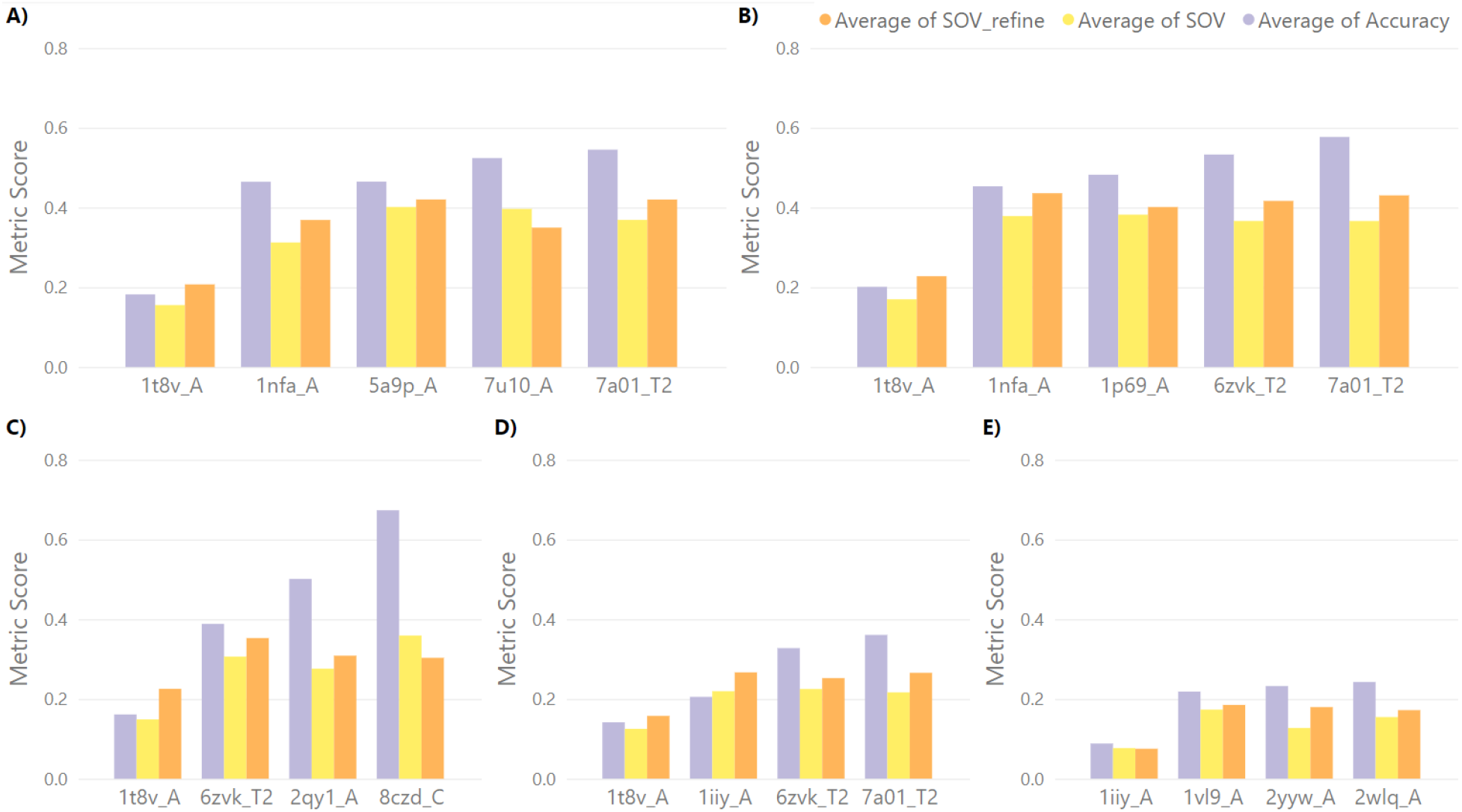
Limitations on Average and Low performing methods. Worst performing proteins for each of the “average’ and “low’ performing methods. As with top performing methods, “1t8v_A’ is commonly predicted incorrectly. Prediction methods: A) SPOT-1D, B) SPOT-1D-LM, C) SPOT-1D-Single, D) Raptor-X Property, E) RGN2.

Low performing methods lack a performance consensus, as their varied methodologies lead to differing results.

Raptor-X Property struggles with virus-binding proteins, but no consistent pattern was found among other poorly predicted proteins. Despite its low performance, it is the fastest prediction method among all analyzed.

SPOT-1D-Single is arguably the best among the low performing methods, with slightly higher overall performance than RGN2 and Raptor-X Property. However, it has difficulty predicting membrane and lipid transport proteins, where average-performing methods excel.

RGN2, a 3D structure prediction method, is fast but ranks low in performance. Its speed depends on a structure relaxation process, which can take longer for poorly predicted structures. RGN2 struggles with proteins related to regulation, metabolism, and metal ion binding.

The results for each low performing method are presented in Fig. 23 C, Fig. 23 D, and Fig. 23 E.

An exceptional case in our dataset involves a protein (PDB ID 1t8v, a fatty-acid binding protein) that posed significant challenges for all structure prediction methods but was predicted more accurately by RGN2, a low-performing method. As shown in Fig. 24, RGN2 outperformed all other methods for chain A of this protein. Notably, this protein has few homologous sequences, with only 14 entries exceeding 90% similarity in our non-redundant dataset aligned using protein-BLAST.

**Figure 24.**
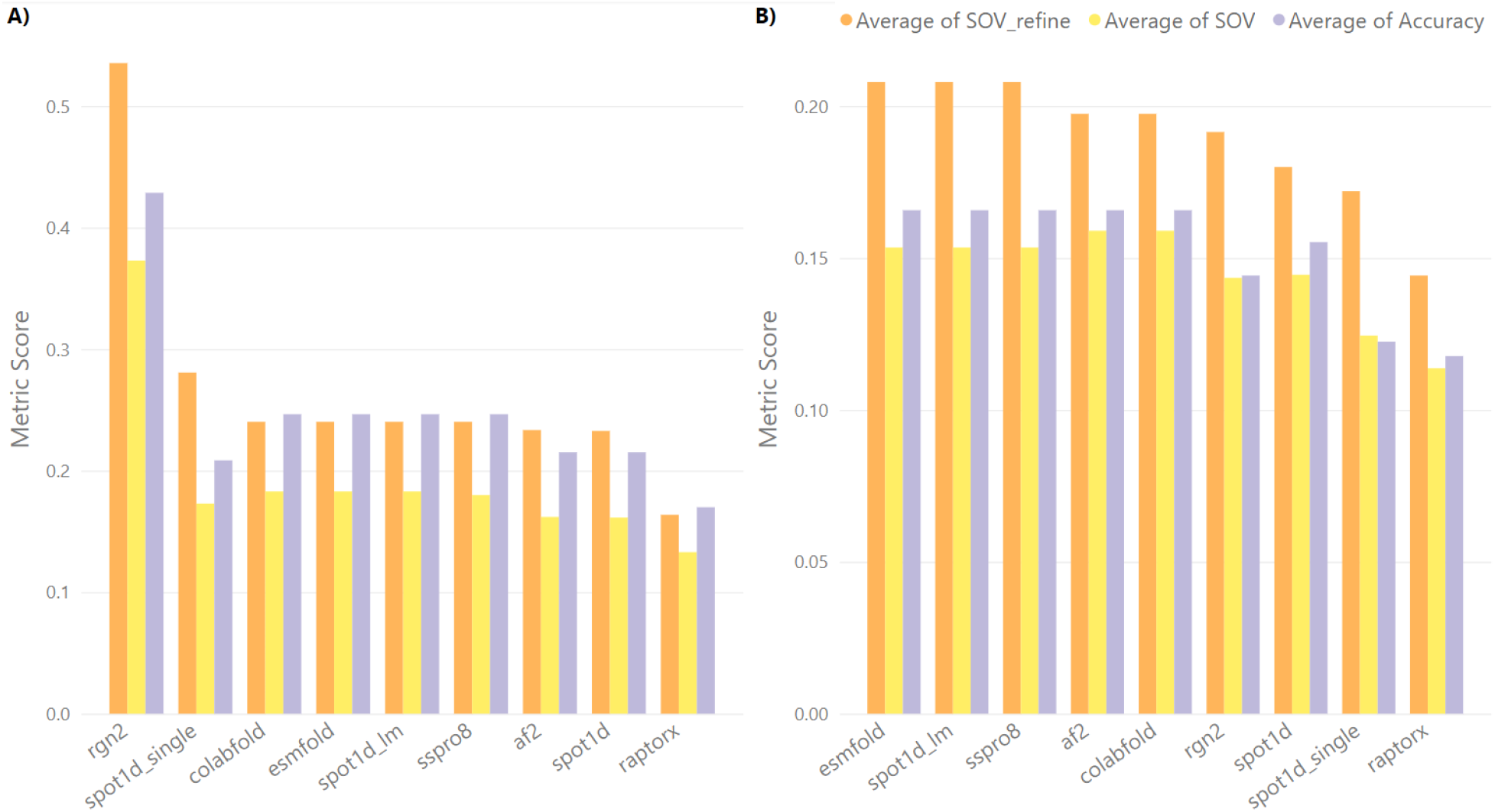
Exceptional Case: RGN2. A challenging protein to predict for all methods (PDB ID 1t8v_A), where the low performing method RGN2 outperforms all others. A) Performance difference from all other methods on local vicinity.B) No difference or very low difference to other prediction methods for distant vicinity.

### Temperature factor and confidence results

The Temperature factor [67] in crystallographic data measure the attenuation of the X-rays by thermal motion due to atomic vibrations. This can lead to inaccuracies in the crystallographic data as the atoms’ positions become difficult to discern.

Here, we investigate any possible correlations with the Temperature factor of proteins within the PDB, and the confidence scores produced by tertiary structure prediction methods. This is done by comparing the temperature factor and confidence score in relation to the single amino acid mutation position in the protein. Unsurprisingly, no correlation could be found for the Temperature factor, as it is prone to bias from experimental conditions, e.g., thermal motion, or crystal purity.

Confidence scores for each predicted secondary structure, or location of *C*_*α*_-atom for each amino acid, also did not provide a strong correlation to the single amino acid mutation location. In some cases, the variance of the confidence score can become the maximum value near the mutation location; but in general, the confidence score of the method, which is present where the temperature factor would be in PDB data, is not predictive of mutation location. These results can be visualized in Fig. 25.

**Figure 25.**
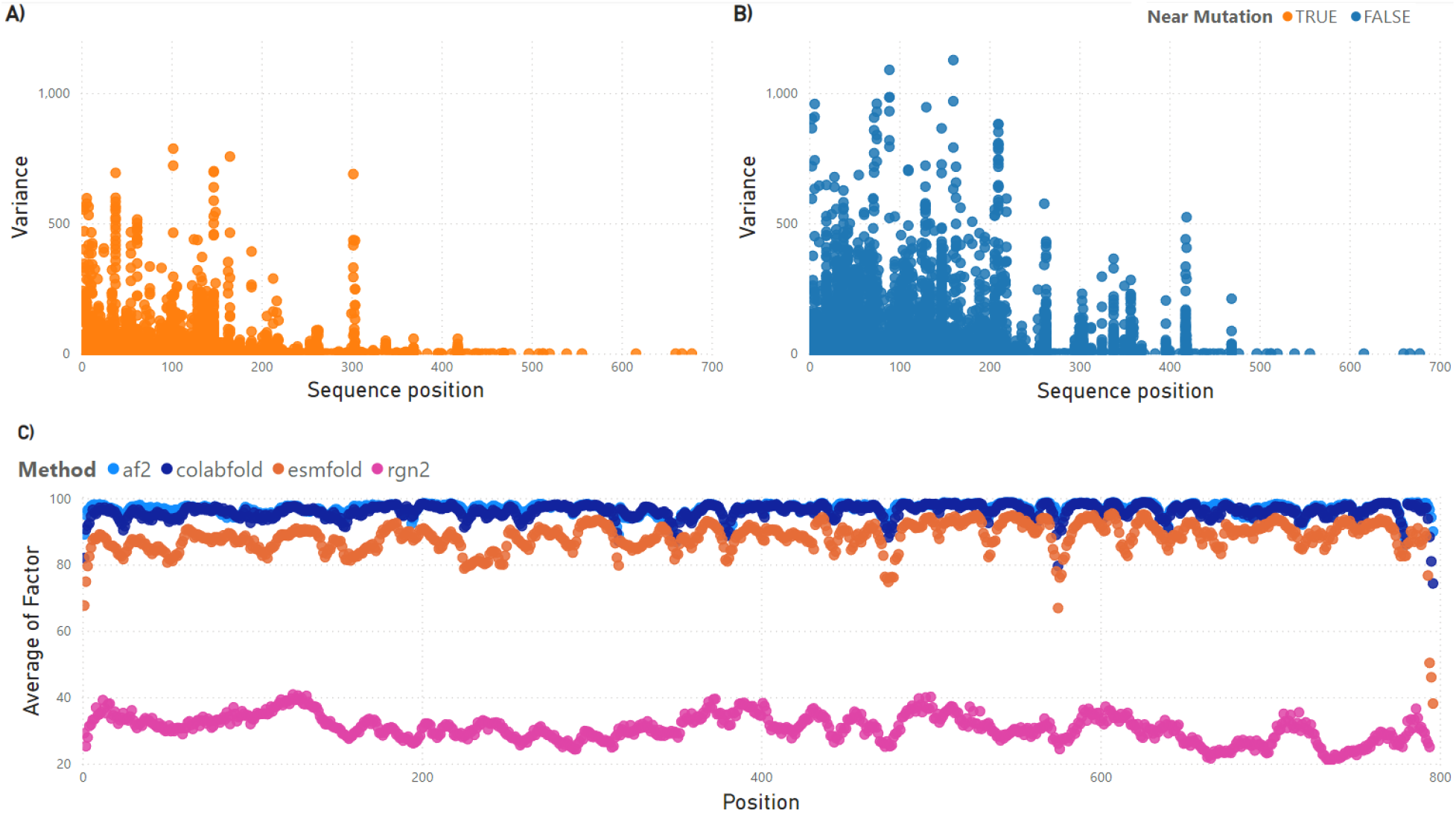
Temperature factor and confidence results. No significant correlation to single amino acid mutations found for both temperature factors in PDB data and confidence scores in predictions. A) Variance value in temperature factor when a mutation is near the sequence location. B) Variance value in temperature factor whena mutation is far from the sequence location. C) Confidence values for all tertiary structure prediction methods in mutations around position 260. This position have low variance when a mutation is near, but high variance when a mutation is not near. As seen from the figure, there is no indication that a mutation has taken place around position 260 from the confidence scores.

## Conclusion

In this work, we evaluate state-of-the-art methods for predicting backbone secondary structure changes caused by missense mutations. To our knowledge, this is the first evaluation of protein prediction capabilities at different mutation vicinity levels. For this purpose, we created a dataset of missense mutations containing primary, secondary, and tertiary structures derived from experimental data. The evaluation includes five secondary structure prediction methods and four tertiary structure prediction methods, each employing vastly different methodologies.

Our analysis reveals that all methods struggle to predict stable mutations—those that do not cause structural changes—often favoring secondary structure element changes even when experimental data does not support them. Experimental data shows logical patterns for secondary structural changes, such as 3-10 helices transforming only into *α*-helices, bends, or turns. In contrast, prediction methods lack the capability to infer the improbability of drastic changes, such as a 3-10 helix turning into a *β*-sheet from a single amino acid mutation.

Although the benchmarking dataset is limited by the small amount of available experimental mutation data, it remains a reliable evaluation tool since models generally avoid training on high-homology sequences. This ensures that the dataset effectively tests how models handle proteins with missense mutations they have not encountered during training.

The dataset provided in this work can aid in the training and testing of future models for missense mutations. Our results demonstrate that while current prediction models achieve high accuracy, they exhibit weaknesses that must be addressed when working with mutational data. With this knowledge, it should be possible to refine existing models using our dataset to develop a more stable, mutation-aware secondary structure prediction method.

Additionally, protein chains involved in complexes or requiring ligand binding to adopt their crystallized structure, as observed in PDB data, highlight the need for models that account for bound molecules in their predictions. Membrane and transport proteins in our dataset illustrate these challenges, as even top performing methods struggle to predict their structures accurately.

Finally, the prediction of low-homology proteins may benefit from an ensemble of structure prediction methods with varying homology requirements. This approach could leverage methods that do not rely on homology, which have shown the potential to outperform top-performing methods in cases where high homology information is unavailable.

We are currently developing a refined model with promising results. This model reduces prediction variance, offering more consistently accurate results compared to the evaluated methods.

## Supporting information

### Supplementary Information Document containing supplementary figures and tables

Contains further information on secondary structure classification procedures, metrics calculations, protein statistics, and fine-grained mutation results. This document is attached to the preprint below the main article’s references.

## Acknowledgments

We would like to express our sincere gratitude to Microsoft and University of Alberta for providing the computational resources required for this project. We are grateful for funding from Natural Sciences and Engineering Research Council of Canada for both the University of Alberta and the University of Victoria. Microsoft AI for Health supported us through their Azure grant, while University of Alberta allowed us to use their “Industry Sandbox and AI Computing’ entrepreneurship resources. Their generous contribution enabled us to perform large-scale predictions and analyses that were essential to our research.

## Footnotes

1 https://pdbj.org/

2 https://sparks-lab.org/server/spot-1d-lm/

3 https://github.com/ivanpmartell/SSMetrics

4 https://github.com/ivanpmartell/SSMetrics

5 https://sw-tools.rcsb.org/apps/MAXIT

