## Supplementary Info, Figures, and Tables for "Missense mutations: Backbone structure positional effects"

#### Supplementary information

#### 1 Secondary structure classification

Proteins are composed of one or more structural domains that can fold independently of each other. Different combinations and arrangements of secondary structures can result in different protein folds, which have functional and evolutionary implications [5]. Folds are defined by the arrangement and connectivity of secondary structures in the protein. Protein secondary structure is the local spatial conformation of the protein's backbone atoms. This conformation occurs through the pattern of hydrogen bonds between the amino hydrogen and carboxyl oxygen atoms in the backbone. Secondary structures in protein form as an intermediate before the protein folds into its three dimensional tertiary structure. The two most common secondary structures are $\alpha$ helices and $\beta$ sheets, which were first theorized by Pauling and Corey [49].

- $\alpha$ helices are right-handed spiral conformations of polypeptide chains. In $\alpha$ helices, every backbone amino $(N - H)$ group donates a hydrogen bond to the backbone carbonyl $(C = O)$ group, which is located four residues prior. This creates a bond between these groups and links them into its spiral conformation.
- $\beta$ sheets are also formed by hydrogen bonding between carbonyl and amino groups that make up the protein backbone and cause the molecule to bend and fold into a pleated sheet form.

Secondary structures were first classified into three categories, including the two most common structures and referring to everything else as coil. Afterwards, the need for a more detailed classification lead to the creation of algorithmic structure categories by Kabsch and Sander [37] which were subsequently refined into the following classes,

- 3-turn helix (3 10 helix). A helix-like structure with a minimum length of 3 residues. This class is denoted by the letter G.
- 4-turn helix ($\alpha$ helix). A helix-like structure with a minimum length of 4 residues. This class is denoted by the letter H.
- 5-turn helix ($\pi$ helix). A helix-like structure with a minimum length 5 residues. This class is denoted by the letter I.
- hydrogen bonded turn (3, 4 or 5 residue turn). Forms a turn-like structure and is denoted by the letter T.
- extended strand in a $\beta$-sheet conformation forming a pleated sheet structure. This class is denoted by the letter E.
- residue in an isolated $\beta$-bridge (single pair $\beta$-sheet hydrogen bond formation). This class is denoted by the letter B.
- bend (the only non-hydrogen-bond based assignment). This class is denoted by the letter S.
- coil (none of the above), denoted by the letter C.
- PPII helix (polyproline helix). A helix-like structure made up of repeating proline residues. This class is denoted by the letter P.

The previous list consists of nine categories, as the polyproline helix was later added to a newer version of the DSSP software remade by the PDB team.

#### 2 Secondary structure assignment

We used DSSP to assign secondary structures to the protein MMCIF' structure data. The macromolecular crystallographic information file (mmCIF) format was selected because the PDB has required it for all new submissions since 2019 [2]. As a result, structures submitted to the PDB after 2019 are no longer available in the older PDB' format, rendering older software incompatible with recent data. DSSP has been updated to process the mmCIF format and has been extensively tested by the community.

DSSP's algorithm for assigning secondary structures to mmCIF-formatted input has evolved from its original version, which processed 'PDB' files. The main differences include both the file formatting and the secondary structure classification, which varies from the traditional 8 classes, as shown in Table S1.

When producing mmCIF output, DSSP expands the input file by rewriting it and appending the secondary structure information. However, this rewriting process can introduce formatting errors, particularly when the input file contains quotations with the character **'**. To address these issues, we provide a script that properly re-formats DSSP's mmCIF output.

| DSSP Class (Q8) | PDB Class | mmCIF Class | Description |
|---|---|---|---|
| B | B | STRN | $\beta$-bridge |
| C | ' '(space) | OTHER | Loop or Coil |
| E | E | STRN | Strand |
| G | G | HELX_RH_3T_P | 3-10 helix |
| H | H | HELX_RH_AL_P | $\alpha$-helix |
| I | I | HELX_RH_PI_P | $\pi$-helix |
| S | S | BEND | Bend |
| T | T | TURN_TY1_P | Turn |
| | P | HELX_RH_PP_P | PPII-helix |

Table S1: DSSP class conversion by input format. Note that classes $\beta$-bridge and Strand are indistinguished by the mmCIF DSSP algorithm.

As noted earlier and shown in Table S1, the mmCIF format classifies both the $\beta$-bridge and Strand under the same STRN class, making it difficult to distinguish between them. Kabash and Sander, in their original DSSP publication, refer to $\beta$-bridge as an "isolated bridge", which is formed by a single hydrogen bond similar to those found in a $\beta$-sheet (DSSP's Strand class).

According to Kabash and Sander's description of secondary structures, an isolated bridge is part of the repeating hydrogen-bonding patterns "turn" and "bridge." Repeating turns form "helices," repeating bridges form "ladders," and connected ladders form "sheets." Based on this understanding, we convert the mmCIF output back to the original classes used in the 'PDB' format, which are recognized by all secondary structure prediction tools. Specifically, when we encounter an isolated STRN, we convert it to its corresponding $\beta$-bridge (**B**) class.

It's also important to note that the PolyProline (PPII) class is a newer addition in the mmCIF format. However, since prediction methods rely on the Q8 assignment, the PPII class is not used in our analysis. Additionally, the OTHER class does not appear in the output, as DSSP only generates proper secondary structural classes. The OTHER class can be inferred when a residue lacks a structural classification.

By applying these conversions, we can accurately assess mmCIF-formatted structure files in terms of the original Q8 classification scheme.

#### 3 Mutation extraction

Finding mutations from an a series of aligned sequences and assigning a wild-type or mutation value to a sequence is non-trivial. To find mutations in our preprocessed data, we created a similar method from Weblogo [18] which use information theory to provide the significance of each mutation. The height of each mutation in the logo is characterized by the frequency of the amino acid at a specific position ($p_i$) and subsequently through Shannon entropy,

$$S = -\sum_i p_i \log_2 p_i \tag{S1}$$

Equation S1 ranges from $[0, 1]$ with a domain of frequencies of $[0, 1]$ and has a characteristic bell shaped curve with a maximum on $0.5$. Therefore, values for frequencies that are equally spaced apart from the maximum value give the same result. Unfortunately, we require a method that can distinguish between such values since the wild-type frequency might be equally spaced apart from the mutation frequency (e.g. $0.25$ and $0.75$), but clearly one is more frequent than the other. Therefore we decided to modify the equation as follows,

$$S = -\sum_i p_i \log_2 (1 - p_i) \tag{S2}$$

This was done to obtain mutation positions easily since positions without mutations will return 0 or Infinity. Infinity values are subsequently turned to 0. Any non-zero value will be valid for a mutation or wild-type amino acid in a position where a mutation has occurred. By using sparse matrices, it is then simple to systematically find these non-zero values and assign its sequence either a wild-type (maximum value) or mutation (others) category for a location in the sequence.

#### 4 Protein data statistics for Mutational Sufficiency

Proteins were filtered to ones containing only single amino acid mutations. We show that the filtering process does not significantly alter the distribution of proteins lengths before and after the filtration procedure in the Fig. S1 and Fig.S2. Most changes occur in long proteins with over 600 amino acids in length. These proteins contained a higher amount of mutations as expected from their length. Clusters with a high amount of proteins also decrease as duplicate single amino acid mutations found in several proteins inside a cluster were filtered out by the Mutation Sufficiency process.

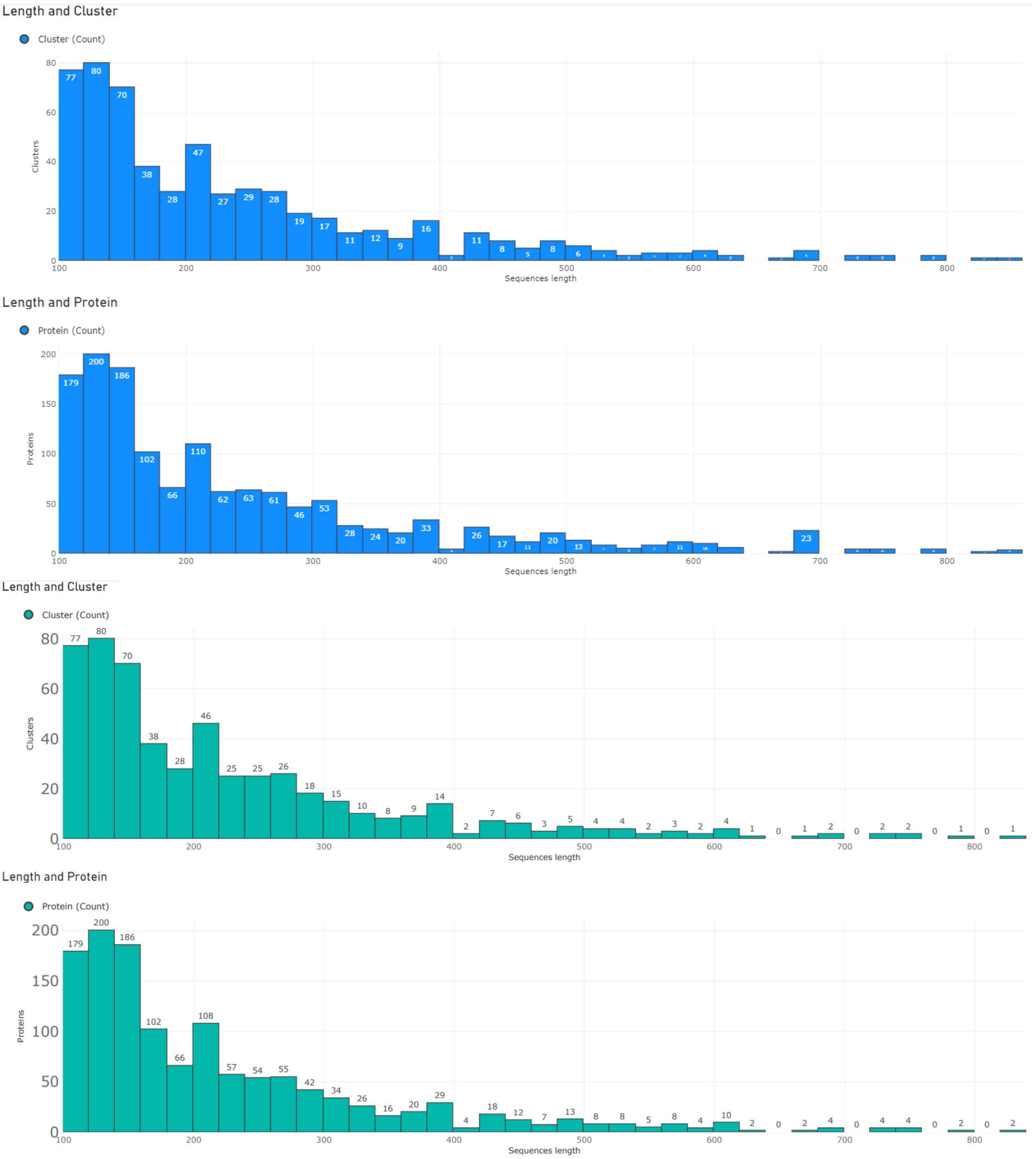


Figure S1: **Sequence lengths by cluster and protein** Top (blue): Amount of clusters for sequence lengths of proteins, and amount of proteins with certain sequence length before filtering. Bottom (green): After $MutationSufficiency$ filtering.

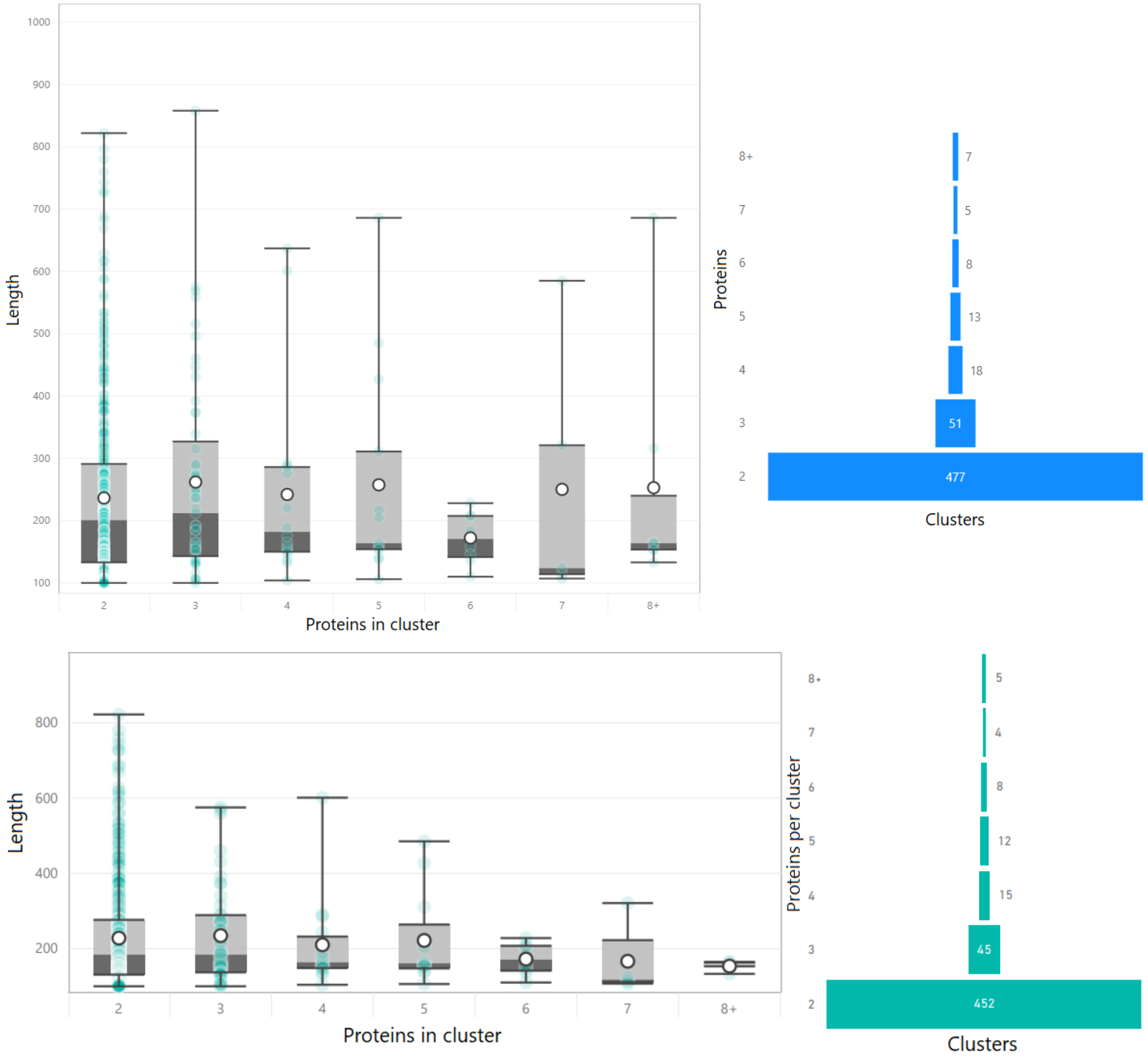


Figure S2: **Length of proteins in clusters per amount of proteins.** Top (blue): Before filtering. Bottom (green): After $MutationSufficiency$ filtering.

#### 5 Mutation performance from prediction methods

Section containing secondary structure metrics obtained for all prediction methods tested in the main text. 'Top' performing methods are contained in Tables S2 and S3. 'Average' performing methods are shown in Table S5, and finally 'Low' performing methods are shown in Table S4.

Table S2: **Single amino acid mutation benchmark on secondary structure for 'top' performing methods.**

| Method | Type | Vicinity | Accuracy | SOV99 | SOV_refine |
|---|---|---|---|---|---|
| af2 | 1d | distant | 0.928 | 0.925 | 0.938 |
| | | global | 0.925 | 0.922 | 0.935 |
| | | local | 0.914 | 0.914 | 0.923 |
| | 2d | distant | 0.927 | 0.924 | 0.937 |
| | | global | 0.925 | 0.922 | 0.935 |
| | | local | 0.911 | 0.910 | 0.919 |
| | 3d | distant | 0.926 | 0.923 | 0.937 |
| | | global | 0.925 | 0.922 | 0.935 |
| | | local | 0.916 | 0.905 | 0.918 |
| | contact | distant | 0.926 | 0.923 | 0.936 |
| | | global | 0.925 | 0.922 | 0.935 |
| | | local | 0.918 | 0.910 | 0.922 |
| colabfold | 1d | distant | 0.929 | 0.926 | 0.938 |
| | | global | 0.927 | 0.924 | 0.936 |
| | | local | 0.919 | 0.919 | 0.928 |
| | 2d | distant | 0.929 | 0.925 | 0.938 |
| | | global | 0.927 | 0.924 | 0.936 |
| | | local | 0.915 | 0.914 | 0.924 |
| | 3d | distant | 0.928 | 0.924 | 0.937 |
| | | global | 0.927 | 0.924 | 0.936 |
| | | local | 0.920 | 0.910 | 0.923 |
| | contact | distant | 0.927 | 0.923 | 0.937 |
| | | global | 0.927 | 0.924 | 0.936 |
| | | local | 0.922 | 0.915 | 0.927 |

Table S3: **Single amino acid mutation benchmark on secondary structure for 'top' performing methods.**

| Method | Type | Vicinity | Accuracy | SOV99 | SOV_refine |
|---|---|---|---|---|---|
| esmfold | 1d | distant | 0.896 | 0.891 | 0.908 |
| | | global | 0.893 | 0.888 | 0.904 |
| | | local | 0.882 | 0.879 | 0.891 |
| | 2d | distant | 0.895 | 0.891 | 0.907 |
| | | global | 0.893 | 0.888 | 0.904 |
| | | local | 0.874 | 0.870 | 0.883 |
| | 3d | distant | 0.895 | 0.890 | 0.908 |
| | | global | 0.893 | 0.888 | 0.904 |
| | | local | 0.877 | 0.863 | 0.881 |
| | contact | distant | 0.894 | 0.889 | 0.907 |
| | | global | 0.893 | 0.888 | 0.904 |
| | | local | 0.880 | 0.868 | 0.885 |
| sspro8 | 1d | distant | 0.943 | 0.940 | 0.951 |
| | | global | 0.940 | 0.937 | 0.948 |
| | | local | 0.932 | 0.931 | 0.940 |
| | 2d | distant | 0.942 | 0.939 | 0.950 |
| | | global | 0.940 | 0.937 | 0.948 |
| | | local | 0.928 | 0.926 | 0.935 |
| | 3d | distant | 0.941 | 0.938 | 0.950 |
| | | global | 0.940 | 0.937 | 0.948 |
| | | local | 0.934 | 0.926 | 0.940 |
| | contact | distant | 0.940 | 0.938 | 0.949 |
| | | global | 0.940 | 0.937 | 0.948 |
| | | local | 0.937 | 0.931 | 0.942 |

Table S4: **Single amino acid mutation benchmark on secondary structure for 'low' performing methods.**

| Method | Type | Vicinity | Accuracy | SOV99 | SOV_refine |
|---|---|---|---|---|---|
| raptorx | 1d | distant | 0.605 | 0.553 | 0.576 |
| | | global | 0.600 | 0.547 | 0.571 |
| | | local | 0.584 | 0.543 | 0.558 |
| | 2d | distant | 0.604 | 0.553 | 0.576 |
| | | global | 0.600 | 0.547 | 0.571 |
| | | local | 0.563 | 0.526 | 0.539 |
| | 3d | distant | 0.603 | 0.550 | 0.579 |
| | | global | 0.600 | 0.547 | 0.571 |
| | | local | 0.571 | 0.508 | 0.534 |
| | contact | distant | 0.600 | 0.547 | 0.578 |
| | | global | 0.600 | 0.547 | 0.571 |
| | | local | 0.583 | 0.524 | 0.550 |
| rgn2 | 1d | distant | 0.627 | 0.607 | 0.630 |
| | | global | 0.622 | 0.602 | 0.625 |
| | | local | 0.598 | 0.583 | 0.600 |
| | 2d | distant | 0.626 | 0.605 | 0.628 |
| | | global | 0.622 | 0.602 | 0.625 |
| | | local | 0.582 | 0.568 | 0.585 |
| | 3d | distant | 0.627 | 0.604 | 0.631 |
| | | global | 0.622 | 0.602 | 0.625 |
| | | local | 0.584 | 0.554 | 0.582 |
| | contact | distant | 0.626 | 0.602 | 0.631 |
| | | global | 0.622 | 0.602 | 0.625 |
| | | local | 0.593 | 0.561 | 0.592 |
| spot1d_single | 1d | distant | 0.666 | 0.622 | 0.649 |
| | | global | 0.661 | 0.616 | 0.642 |
| | | local | 0.641 | 0.610 | 0.628 |
| | 2d | distant | 0.664 | 0.621 | 0.648 |
| | | global | 0.661 | 0.616 | 0.642 |
| | | local | 0.623 | 0.594 | 0.613 |
| | 3d | distant | 0.663 | 0.618 | 0.651 |
| | | global | 0.661 | 0.616 | 0.642 |
| | | local | 0.630 | 0.576 | 0.604 |
| | contact | distant | 0.660 | 0.615 | 0.649 |
| | | global | 0.661 | 0.616 | 0.642 |
| | | local | 0.643 | 0.590 | 0.619 |

Table S5: **Single amino acid mutation benchmark on secondary structure for 'average' performing methods.**

| Method | Type | Vicinity | Accuracy | SOV99 | SOV_refine |
|---|---|---|---|---|---|
| spot1d | 1d | distant | 0.793 | 0.783 | 0.810 |
| | | global | 0.789 | 0.780 | 0.807 |
| | | local | 0.775 | 0.771 | 0.792 |
| | 2d | distant | 0.792 | 0.782 | 0.809 |
| | | global | 0.789 | 0.780 | 0.807 |
| | | local | 0.761 | 0.754 | 0.777 |
| | 3d | distant | 0.791 | 0.778 | 0.808 |
| | | global | 0.789 | 0.780 | 0.807 |
| | | local | 0.768 | 0.744 | 0.776 |
| | contact | distant | 0.790 | 0.776 | 0.808 |
| | | global | 0.789 | 0.780 | 0.807 |
| | | local | 0.774 | 0.750 | 0.780 |
| spot1d_lm | 1d | distant | 0.825 | 0.812 | 0.836 |
| | | global | 0.821 | 0.808 | 0.833 |
| | | local | 0.807 | 0.797 | 0.817 |
| | 2d | distant | 0.824 | 0.810 | 0.835 |
| | | global | 0.821 | 0.808 | 0.833 |
| | | local | 0.797 | 0.786 | 0.807 |
| | 3d | distant | 0.822 | 0.807 | 0.836 |
| | | global | 0.821 | 0.808 | 0.833 |
| | | local | 0.804 | 0.779 | 0.809 |
| | contact | distant | 0.821 | 0.806 | 0.835 |
| | | global | 0.821 | 0.808 | 0.833 |
| | | local | 0.807 | 0.783 | 0.811 |

#### 6 Protein properties

In the main text, we include properties for each of the proteins of interest and to show the strengths and weaknesses of the prediction methods. These properties were agglomerated from CATH, SCOP and PDB descriptor data. We generated word clouds to obtain the most common words, which resulted in the descriptors for the proteins. This section contains all the figures with their corresponding protein property word clouds.

The bar plots are shown in the main text, but we decide to include both the plots and word clouds together to more easily match the properties to their corresponding proteins. The larger the word in the word cloud, the more common the property is for the respective bar plot proteins. Figures for protein properties: S3, S4, S5, S6, S7, S8, S9.

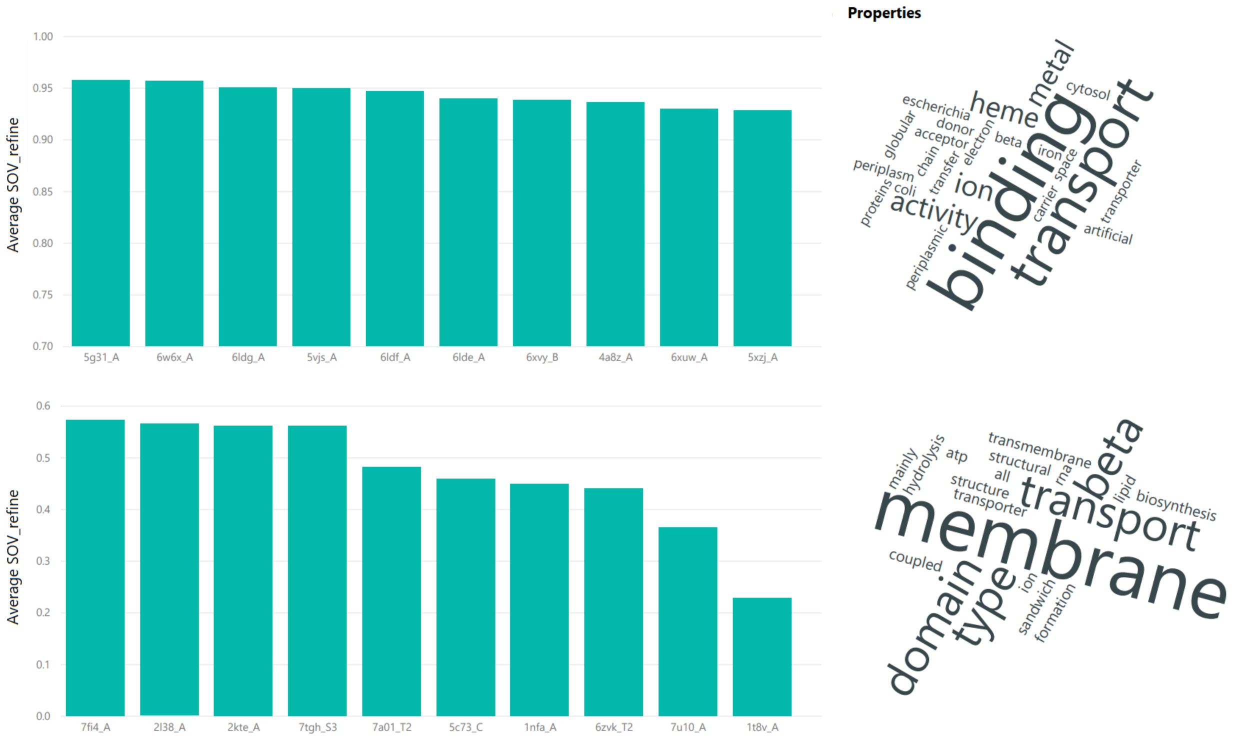


Figure S3: **Overall protein structure prediction results.** Top: Best overall predicted proteins and their properties. Bottom: Worst overall predicted proteins and their properties.

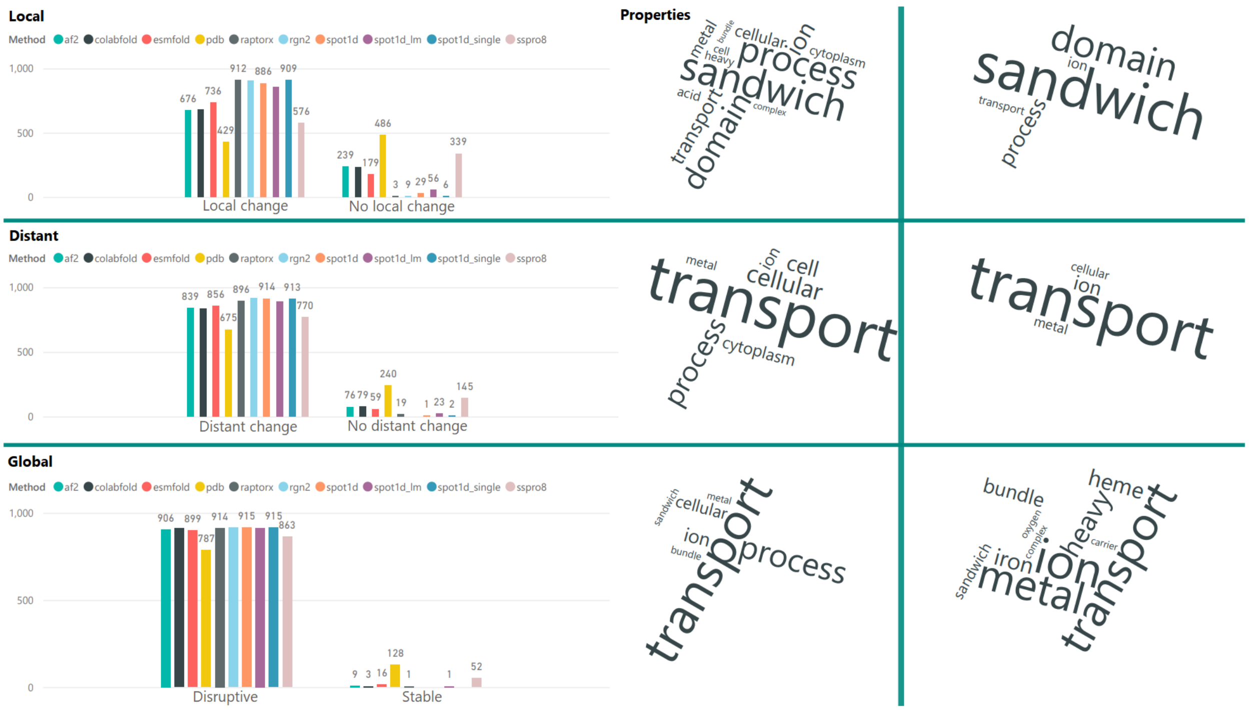


Figure S4: **Mutation stability results.** Disruptive and Stable mutations. Stabilizing mutations occur more often in PDB data than in prediction methods, as the latter almost always predicts destabilizing mutations. The exception is SSPro8 while still missing two thirds of stabilizing mutations.

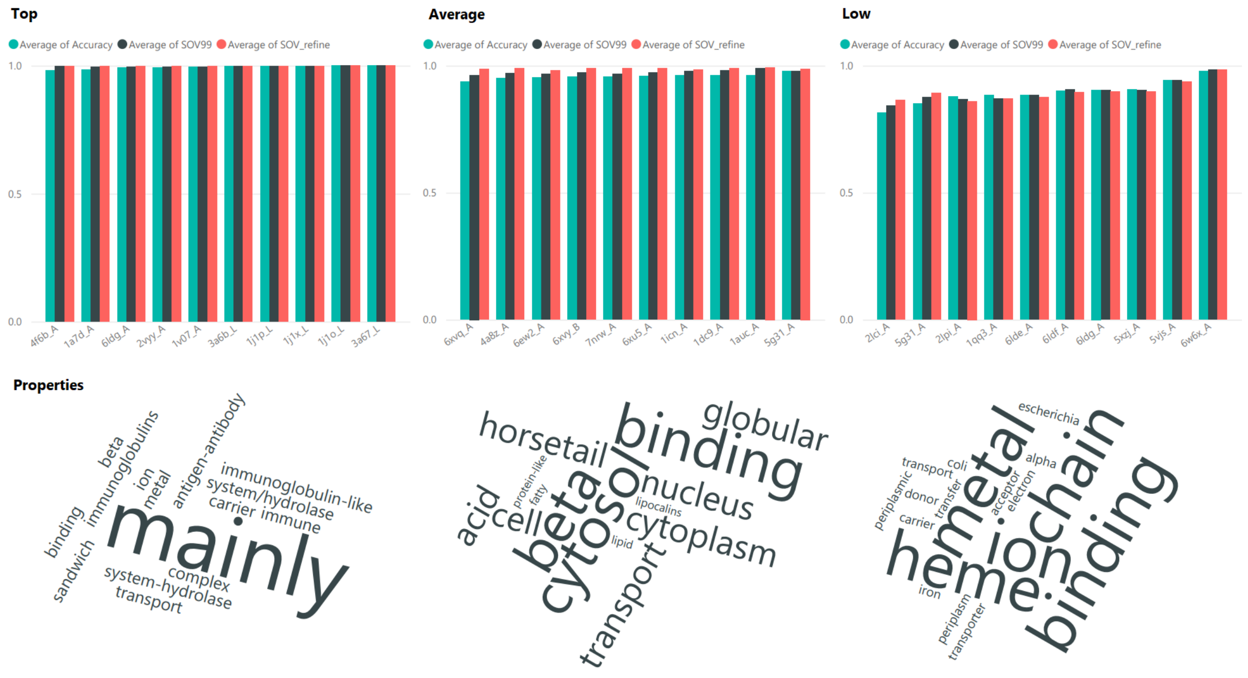


Figure S5: **Best results per method category.** Best predicted proteins along with their properties for each method category (top-performing, average performing and low performing). From left to right, Top performing methods, Average performing methods, and Low performing methods.

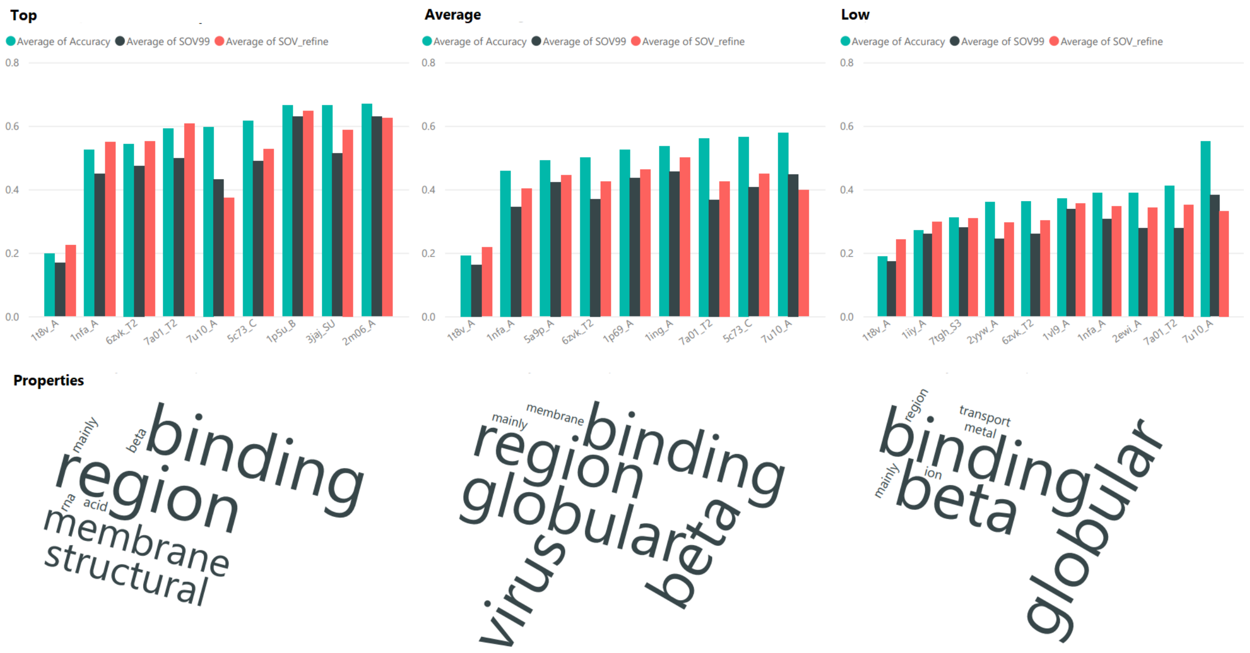


Figure S6: **Worst-predicted proteins per method category.** The worst-predicted proteins along with their properties for the different method categories. From left to right, Top performing methods, Average performing methods, and Low performing methods.

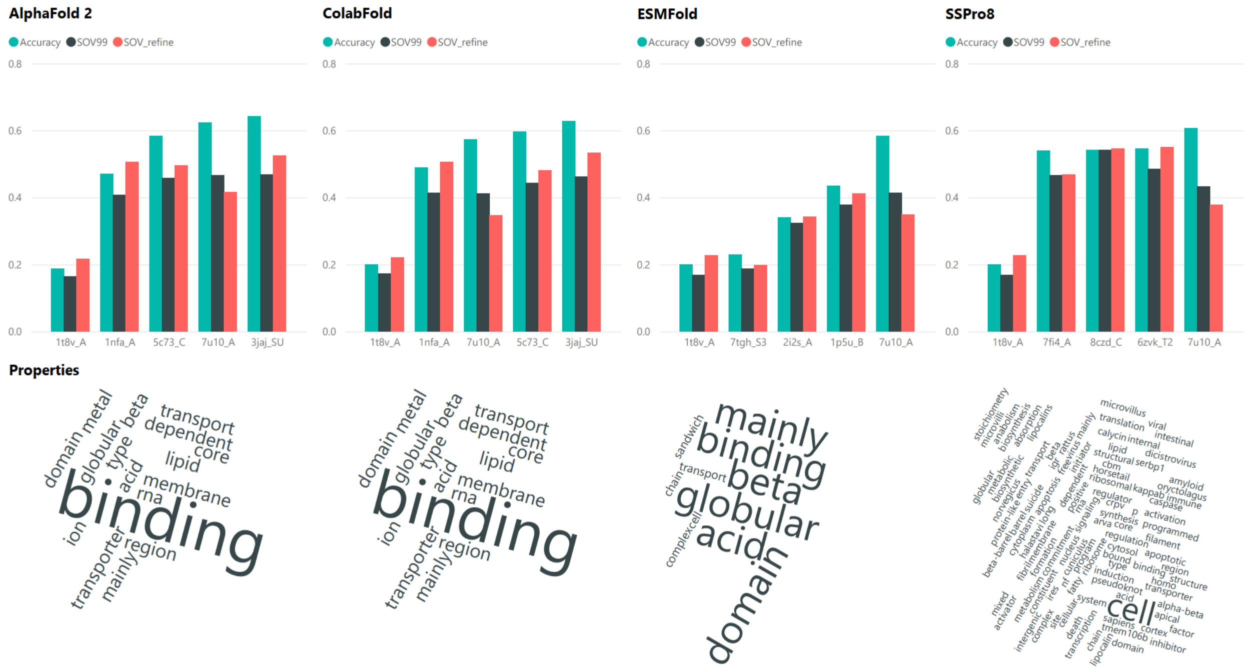


Figure S7: **Top performing methods.** Worst performing proteins for each of the top performing methods. From left to right: AlphaFold2, ColabFold, ESMFold, and SSPro8.

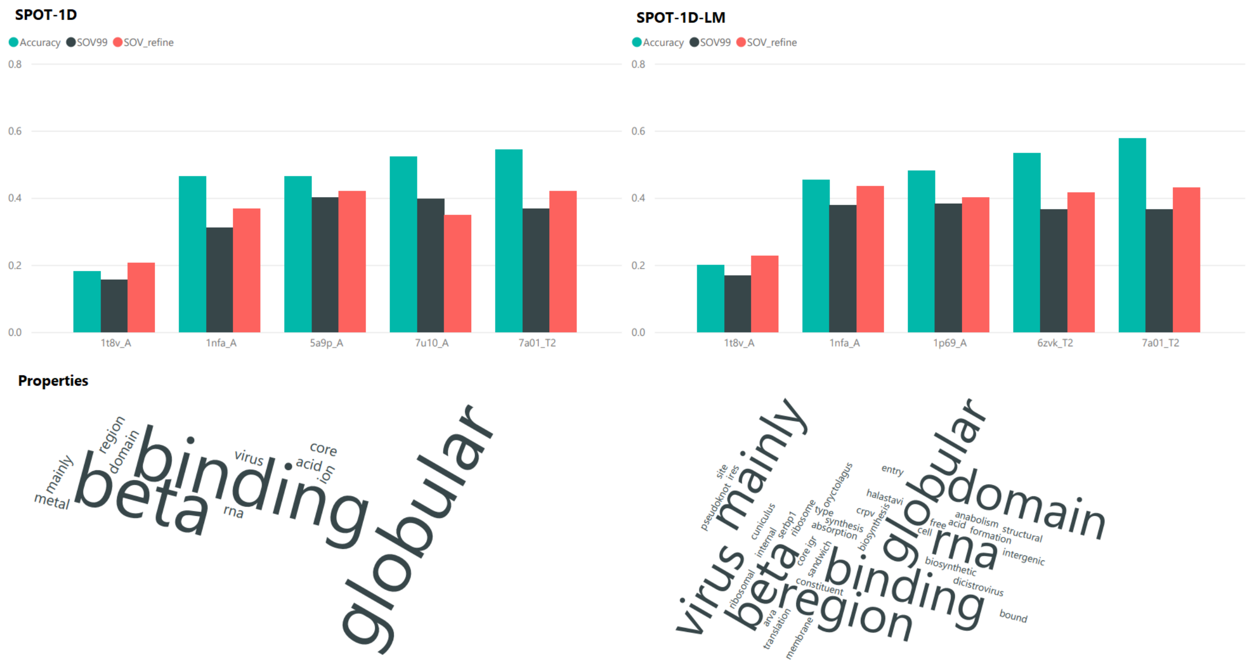


Figure S8: **Average performing methods.** Worst performing proteins for each of the average performing methods. Left: SPOT-1D, Right: SPOT-1D-LM.

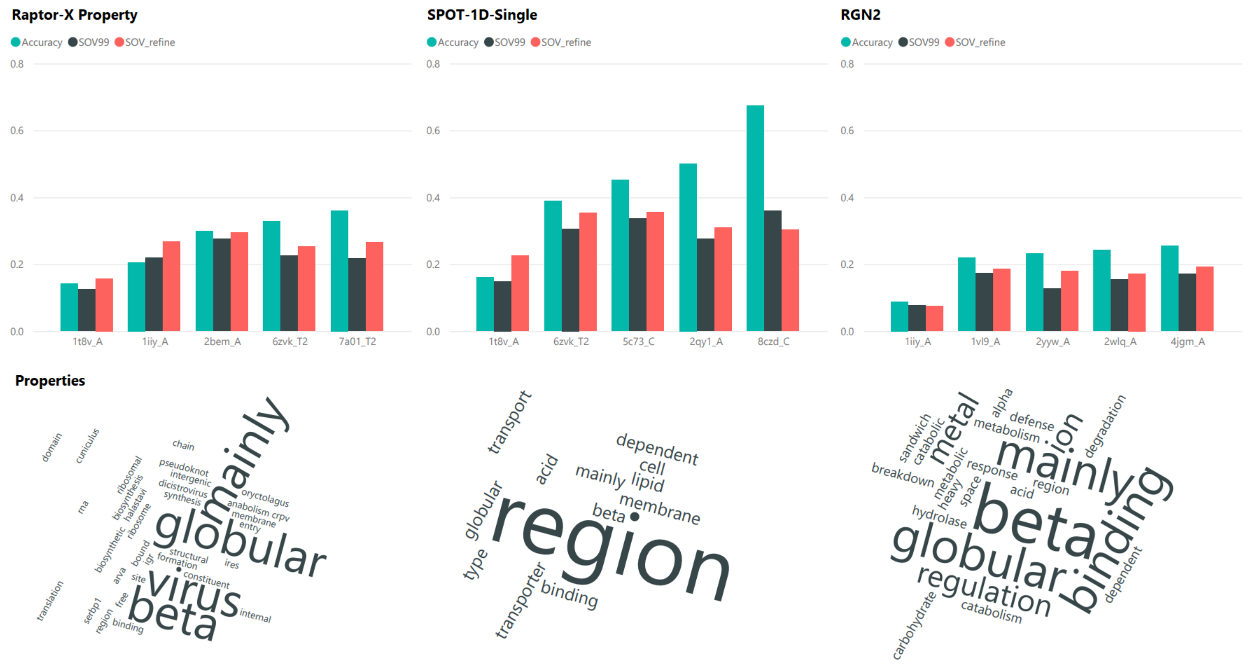


Figure S9: **Low performing methods.** Worst performing proteins for each of the low performing methods. From left to right: Raptor-X Property, SPOT-1D-Single, and RGN2.

#### 7 Details on prediction methods

Fig. S10 provides a detailed view of the prediction method results for SOV_REFINE, where boxes represent the 25th to 75th percentiles and whiskers indicating values within 1.5 interquartile range (IQR). The spread reveals that even top-performing methods struggle with some proteins, achieving below 50% in SOV_REFINE.

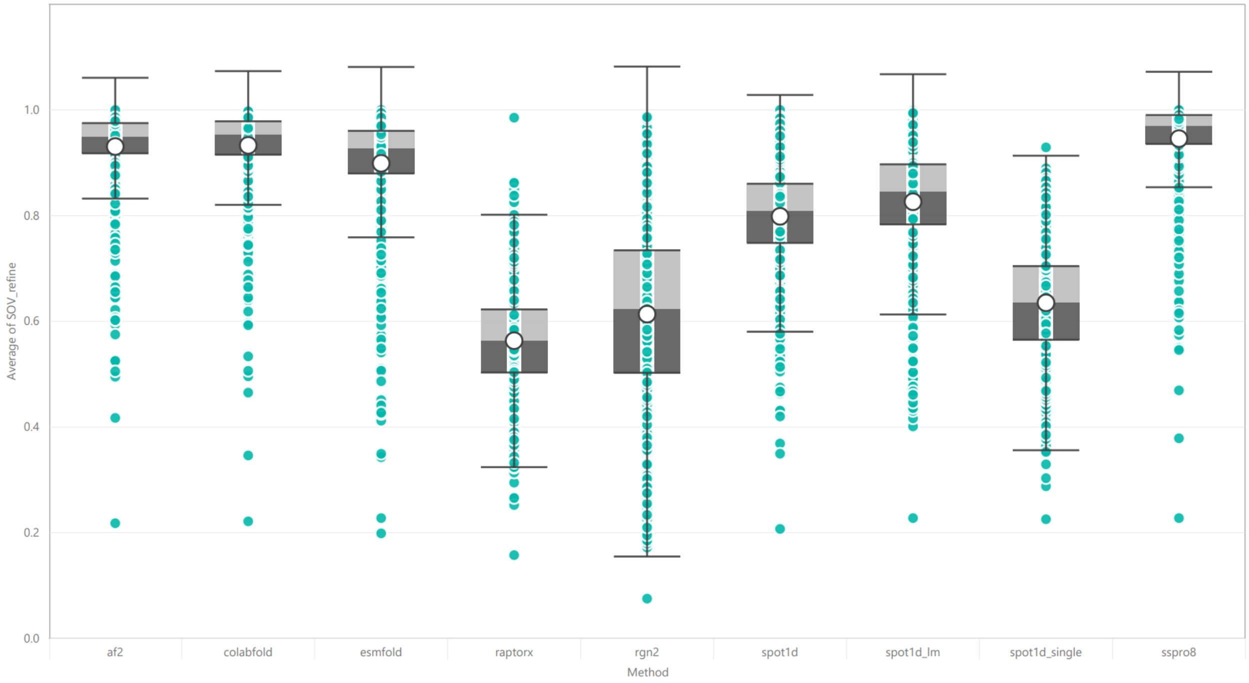


Figure S10: **SOV_REFINE of each structure prediction method.** The boxes range between 25 and 75 percentiles, while the whiskers encompass 1.5 IQR. The white circles depicts each method's mean SOV_REFINE score.

#### 8 Secondary structure prediction tools

Protein secondary structure prediction is an active area of research; significant progress has been made in the past few years through machine learning models. Secondary structure prediction has been defined multiple times, but the most common involve three-state and eight-state classification. Three-state classification contains 3 types of secondary structures, while eight-state classification contains 8 types of secondary structures allowing for more fine-grained structures. The highest three-state classification accuracy without relying on structure templates is around 85% [76]. These improvements came from increasingly larger databases of protein sequences and structures for training, the use of template secondary structure information and more powerful deep learning techniques. As we are approaching the theoretical limit of three-state prediction of 90%, focus shifted on solving the eight-state prediction problem, which is much more complicated and challenging. The theoretical limit of secondary structure prediction for eight classes is not well established, but current template-less methods manage around 70% [56]. Although controversy exists around the eight classes as they are not equally distributed and some are more difficult to predict than others, further improvements are still being achieved.

The field's improvements started to slow down at the fifth iteration of the CASP competition, which took place in 2002. This meeting was the last CASP meeting that accepted secondary structure predictions. Their methods abstract [13] details each ranked group's methodology for the different predicted categories. The top 10 predictors' results were posted in their meeting manuscript [3]. The models were evaluated on using segment overlap (SOV) accuracy measure [78], which is currently still being utilized in the field.

Current secondary structure prediction models are heavily influenced by these methods, including SSPro, PsiPred, and its derivatives. During the early 2000s, machine learning models were the primary predictors being developed because of their increased prediction accuracy. These models take into account co-evolutionary data for each protein in order to predict the secondary structure. Since CASP5 there have been multiple reviews surveying the status quo in the field [32, 62, 34].

PsiPred [11] is one of the long-standing secondary structure prediction software with over 20 years of development. PsiPred was created as a web server but is also offered as a standalone software. It unfortunately only predicts three-class secondary structure, but its PSSM-based methodology has had a significant impact on eight-class secondary structure prediction methods. PsiPred was originally a multi-layer neural network based method that utilizes position specific scoring matrices (PSSM) generated by PSI-BLAST [35]. PSSM gives PsiPred the co-evolutionary information required to predict a secondary structure by characterizing protein domains. Later PsiPred was updated to a deeper neural network architecture with two hidden layers rather than just one, and with rectifier activations rather than sigmoid. The other change is that the input window has been extended from 15 residues to 33 thanks to using sparse connections between the input layer and first hidden layer [11].

ssPro8 [44] utilizes a three-stage workflow. As with other classic tools, it uses PSI-BLAST to derive multiple sequence alignment and profile probabilities. Also, it uses an ensemble of 100 Bidirectional Recursive Neural Networks (BRNNs) trained on the data to generate a first set of probability predictions for each secondary structure class. Finally, ssPro8 derives secondary structure predictions from the ensemble by using sequence-based structural similarity in regions with more than 45% sequence similarity.

RaptorX Property Predictor [73] was created as a web server, but later offered as a standalone software. It uses a deep learning method with a Convolutional Neural Field architecture [74]. This architecture combines the advantages of both Conditional Random Fields and Convolutional Neural Networks, which captures not only a complex sequence-structure relationship, but also models secondary structure correlation among adjacent residues.

SPOT-1D [29] is a model with an ensemble of nine Bidirectional Recurrent Neural Networks and Residual Networks hybrid models to identify and propagate short and long term dependencies throughout the sequence. This model utilizes predicted data from sequence data consisting of two evolutionary profiles from three iterations of PSI-BLAST and from HHBLITS. The predicted data are used to create an amino acid contact map through SPOT-Contact. Finally, the amino acid sequence is also utilized to predict their physicochemical properties. All these data are utilized as input to the ensemble of neural networks which produces the final secondary structure prediction. Even though SPOT-1D was not part of the latest reviews [32, 62, 34], it was selected since it was compared to other models and found to be well performing among the best protein secondary structure prediction software.

SPOT-1D-Single [59] is an ensemble of three neural network architectures. Similar to SPOT-1D, the architectures are variants of Residual Networks and Bidirectional Recurrent Neural Networks. The main difference is that the input is only a single amino acid sequence, without the evolutionary data from PSI-BLAST, HHBLITS or additional data from SPOT-Contact.

SPOT-1D-LM [60] is an ensemble of large language models that have recently improved prediction accuracy for sequential tasks is many fields. The pre-trained large language models utilized here consist of ESM-1b and ProtTrans which turn the initial single amino acid sequence into features for a deep learning model with the same ensemble architecture as SPOT-1D-Single. The ensemble architecture then outputs the secondary structure prediction. The main advantage for these large language models is the included evolutionary information built into them from their training phase. SPOT-1D-LM managed to get comparable prediction performance to models with evolutionary information, such as SPOT-1D. This, of course with the added speed benefit from not requiring computationally expensive PSI-BLAST and other evolutionary information procuring software.

Therefore, we consider a mix of methods; some that obtain evolutionary (template) information from alignment methods such as PSI-BLAST and HHBlits, methods that contain evolutionary information embedded in the model itself such as with large language models, and template-less models that only utilize the input sequence to make a prediction.

### 9 Tertiary structure prediction tools

Deep learning algorithms have enabled unprecedented progress towards the prediction of tertiary structure from a protein sequence. Recent advances in protein structure prediction are due to AlphaFold2, which has taken the problem of protein structure prediction to another level, reaching similar performance to experimental methods like X-ray crystallography and NMR spectroscopy for proteins with high number of homologous proteins. This allows Alphafold2 to extrapolate the 3D structure from a protein's homologs. Homology is the similarity due to shared ancestry between a pair of structures. This similarity is calculated from the protein's sequence and done through multiple sequence alignment (MSA) of known proteins.

Proteins can be characterized by the angles of certain backbone atoms. Ramachandran angles are torsion angles $\phi$, $\psi$, and $\omega$ that describe rotation of the backbone around the bonds between $N - C\alpha$, $C\alpha - C'$ and $C' - N$ respectively. Angle $\omega$ is rarely used as its value is most often $180°C$, although the discarding of $\omega$ might lead to structure reconstruction inaccuracies. These angles are usually plotted into a graph to obtain statistics about secondary structures of the protein. For computational reconstruction, the use of a library of known protein structures is needed to identify which combination of $\phi$ and $\psi$ angles corresponds to each amino acid residue in the sequence. Once all of the angles have been assigned, a three-dimensional model of the protein that is consistent with those angles can be created. This leads to sidechain atoms being estimated from the $C\alpha$ atoms locations.

Frenet-Serret frames are used in protein modelling to generate the backbone structure of a protein. The backbone geometry is based on two local parameters (curvature and torsion), and these parameters have fixed values for each residue ($C\alpha$). The Frenet-Serret formulas describe motion along the backbone using the reference frame of the backbone itself. This means that the frames require a minimum of 4 $C\alpha$ atoms locations to create the angles needed for reconstruction.

AlphaFold2 [46] is a neural network-based system that can predict protein structures with atomic accuracy. Its protein tertiary structure prediction is done using an amino acid sequence and aligned sequences of homologs as inputs. It processes these inputs through repeated layers of Evoformer blocks, which are neural network layers with attention and multiplicative updates to exchange information between the MSA and the pair representation. The pair representation encodes the spatial and evolutionary relationships between residues as edges in a graph. The Evoformer blocks produce a structural hypothesis that is continuously refined throughout the neural network's training.

The structure module introduces an explicit 3D structure in the form of rotations and translations for each residue of the protein. These representations are initialized in a randomized state and then updated iteratively using invariant point attention (IPA) and equivariant updates. IPA is a geometry-aware attention operation that augments the queries, keys and values with 3D points in the local frame of each residue. The equivariant updates allow the network to implicitly reason about the side-chain atoms and enforce peptide bond geometry. The structure module breaks the chain constraint to allow simultaneous local refinement of all parts of the structure.

The network predicts the side-chain angles, the per-residue accuracy (pLDDT), and the global similarity score (pTM) using small per-residue networks on the final activations. The pLDDT score is the predicted lDDT metric score that the neural network calculates for its predicted amino acids. The pTM score is the predicted TM-score that is calculated by the neural network to assess the predicted structure to the experimentally obtained structure. The final loss term, called frame-aligned point error (FAPE), compares the predicted atom positions to the true positions under many different alignments. This loss term prioritizes the orientational correctness of the residues and provides correct chirality, or asymmetric conformation of a molecule, for AlphaFold2.

The network is trained using supervised learning on PDB data, but also uses self-distillation to enhance accuracy. Self-distillation uses a trained network to predict structures for unlabelled protein sequences and then trains a new network from scratch using a mixture of PDB data and predicted structures. The network also uses a BERT-style objective to predict masked elements of the MSA sequences, which encourages it to learn phylogenetic and covariation relationships. BERT stands for Bidirectional Encoder Representations from Transformers [20]. It is an open-source deep learning method that is used for various language tasks by using a transformer deep learning model architecture that is primarily used for processing sequential data like text or time series [70].

AlphaFold2 is able to handle missing physical context and produce accurate models for challenging cases such as intertwined proteins that fold in the presence of unknown ligands, or molecule that binds to the protein. However, AlphaFold2 performs worse for proteins that have few intra-chain or homotypic (identical subunit) contacts compared to heterotypic (dissimilar subunit) contacts, which typically occur for bridging domains within larger complexes. AlphaFold2 also depends on MSA depth and its accuracy decreases when there are fewer than 30 aligned sequences.

ESMFold [42] shows that language models trained on millions of protein sequences can learn patterns that enable them to predict the three-dimensional structure of proteins at an atomic resolution. It is a fast and accurate deep learning end-to-end single sequence structure predictor, that leverages the evolutionary patterns captured by a language model. ESMFold simplifies the neural architecture and eliminates the need for MSA and templates, resulting in up to 60x speed-up compared to state-of-the-art methods while maintaining high resolution accuracy.

ESMFold predicts the tertiary structure of a protein by inputting the protein sequence into ESM-2, a large-scale transformer language model trained on UniRef [68] protein sequences with masked language modeling objective. ESM-2 produces internal representations that capture evolutionary patterns and structural information. The internal representations are passed to a folding module, which consists of a series of folding blocks and an equivariant transformer structure module. The folding blocks alternate between updating a sequence representation and a pairwise representation, while the structure module outputs 3D coordinates and confidences for each atom in the protein. The folding head performs three steps of recycling, which iteratively refines the predicted structure by feeding back the coordinates into the folding blocks. Recycling improves the accuracy and confidence of the predictions. The final output is an atomic-level structure prediction with predicted-lDDT (plDDT) scores that indicate the confidence and quality of the prediction.

RGN2 [47] combines a protein language model (AminoBERT) that learns latent structural information from unaligned proteins with a recurrent geometric network (RGN) that uses Frenet-Serret formulas, an alternative to ramachandran angles used in previous methods, to generate the backbone geometry of a protein. RGN2 predicts the tertiary structure of a protein by first encoding the amino acid sequence into a high-dimensional vector representation using AminoBERT, a transformer-based language model trained on millions of natural protein sequences. AminoBERT learns to fill in missing residues and permute segments of sequences, capturing global and local information about protein structure. Then, RGN2 uses a recurrent neural network to decode the AminoBERT representation into a sequence of rotation matrices that describe the local geometry of each residue using the Frenet-Serret formulas. These formulas define an orthonormal reference frame at each $C\alpha$ atom along the backbone curve, which can be easily constructed by applying the rotation matrices sequentially. This parameterization is simple, compact, and invariant to translation and rotation of the whole protein. Finally, RGN2 refines the predicted structures using an AF2Rank-based protocol that imputes the backbone and sidechain atoms and optimizes the energy function. The refinement step is not differentiable but improves the quality of the predictions. RGN2 outperforms existing methods that rely on multiple sequence alignments (MSAs) for orphan and designed proteins, which have no known sequence homologs.

#### 10 Alphafold2 batch processing

The prediction of a 3D structure by alphafold requires two main subprocesses. First, multiple sequence alignments (MSA) have to be done to obtain features that are utilized as an input for the following subprocess. Secondly, the neural network component takes the MSA features to produce and refine a predicted 3D structure of the target protein sequence. For long sequences, the first subprocess requires a high amount of RAM (∼128gb), an ordinary amount of CPU processing power (∼8 threads), and no GPU processor. The second subprocess requires an ordinary amount of RAM (∼32gb), a low amount of CPU processing power (∼4 threads), and a high amount of GPU processing power. The differing computing needs from the two subprocesses can be exploited by isolating them and computing them in separate clusters. This was done by ParaFold [81] for use of high performance computing clusters and further refined with concurrent computing by members of the community. Unfortunately, the original project has not been updated with the community improvements. We have updated the project by fixing and adding unfinished functionality for ease of use. Our improvements can be obtained at `https://github.com/ivanpmartell/ParallelFold`.

#### 11 RGN2 local processing

RGN2 was originally made for google colab [6]. This is a service that lets you run python notebooks with a GPU on a server for free. We converted the python notebook code into python files for use in a local environment. The files include run 'run_aminobert.py' and 'run_rng2.py'. They are run successively starting with the aminobert script to obtain a tertiary structure prediction. The prediction is then run through an atomic relaxation (emrefinement [7]) software to remove any inconsistencies in the prediction. We noticed that the original relaxation script from RGN2 had an error which meant that some proteins were not relaxed and finalized to produce a prediction file. We fixed the error in the 'ter2pdb.py' file, which we also include. All files can be found under the 'extra/rgn2_local_files' folder in our main repository for this project.

[6] `https://colab.research.google.com/`

[7] `https://zhanggroup.org/ModRefiner/`
